# Sex-specificity of the *C. elegans* metabolome

**DOI:** 10.1101/2022.08.11.503636

**Authors:** Russell N. Burkhardt, Alexander B. Artyukhin, Erin Z. Aprison, Brian J. Curtis, Bennett W. Fox, Andreas H. Ludewig, Amaresh Chaturbedi, Oishika Panda, Chester J. J. Wrobel, Siu S. Lee, Ilya Ruvinsky, Frank C. Schroeder

## Abstract

Recent studies of animal metabolism have revealed large numbers of novel metabolites that are involved in all aspects of organismal biology, but it is unclear to what extent metabolomes differ between sexes. Here, using untargeted comparative metabolomics for the analysis of wildtype animals and a series of germline mutants, we show that *C. elegans* hermaphrodites and males exhibit pervasive metabolomic differences. Several hundred small molecules are produced exclusively or in much larger amounts in one sex, including a host of previously unreported metabolites that incorporate building blocks from nucleoside, carbohydrate, lipid, and amino acid metabolism. A subset of male-enriched metabolites is specifically associated with the presence of a male germline, whereas enrichment of other compounds requires a male soma. Further, we show that one of the male germline-dependent metabolites, an unusual dipeptide incorporating *N*,*N*-dimethyltryptophan, accelerates the last stage of larval development in hermaphrodites. Our results serve as a foundation for mechanistic studies of how the genetic sex of soma and germline shape the *C. elegans* metabolome and provides a blueprint for the discovery of sex-dependent metabolites in other animals.

## Introduction

Recent studies of metabolism and related physiological responses in humans and rodent models have revealed dramatic differences between sexes^1–5^. For example, targeted metabolomic analysis of human serum revealed significant differences for one-third of annotated metabolites^6^. However, few studies in any model system have leveraged the power of untargeted metabolomic approaches based on high-resolution mass spectrometry (HRMS) to uncover novel, unannotated metabolites associated with sex.

HRMS-based untargeted metabolomic analyses in several species have revealed vast metabolic diversity, including large numbers of metabolites whose chemical structures have not yet been determined^7,8^. Interpretation of the resulting large datasets usually relies on comparative analyses of samples from different biological conditions, which enables identifying metabolites that are significantly associated with a context of interest and thus can be prioritized for detailed chemical characterization^8–11^. HRMS-based comparative metabolomics of different sexes thus has the potential to uncover unannotated metabolites whose identification can advance mechanistic understanding of sex-specific phenotypes and complement transcriptomics and proteomics.

In the model nematode *C. elegans*, discovery-oriented metabolomic analyses^12,13^ have almost exclusively focused on the predominant sex, the self-fertile hermaphrodites, which account for >99% of the populations under standard conditions^14^, whereas the metabolomes of the much less abundant males have been studied only to a limited extent. Notably, over 5500 genes, or ~1/3 of the entire protein-encoding transcriptome, are differentially expressed between the two sexes^15,16^. Correspondingly, there is growing evidence for major sex-specific differences in metabolism, disease response, and other phenotypes in *C. elegans*^17–21^.

An intriguing subset of metabolomic differences between the sexes is comprised of excreted small molecules with which animals communicate. Several recent studies describe diverse life history traits affected by such sex-specific pheromones. The best-studied case involves a pair of ascaroside derivatives that have nearly identical chemical structures (Figure 1a) but are enriched either in hermaphrodites (ascr#3, **1**) or in males (ascr#10, **2**)^22^. At physiological concentrations, the male-specific ascr#10 exerts effects that appear opposite to the effects of ascr#3^23^, including reduced exploratory behavior of hermaphrodites^24,25^, improved aspects of germline function^26–28^, and shortened lifespan^27,29^. More recently, we reported an unusual fatty acid-amino acid conjugate, nacq#1 (**3**), that also shortens the hermaphrodite lifespan^29^, therefore contributing to a phenomenon of male-induced demise, whereby hermaphrodites die more quickly in the presence of male-excreted compounds^19,30,31^. In addition to reducing hermaphrodite lifespan, nacq#1 accelerates the last stage of larval development, resulting in faster sexual maturation^29^.

**Figure 1.**
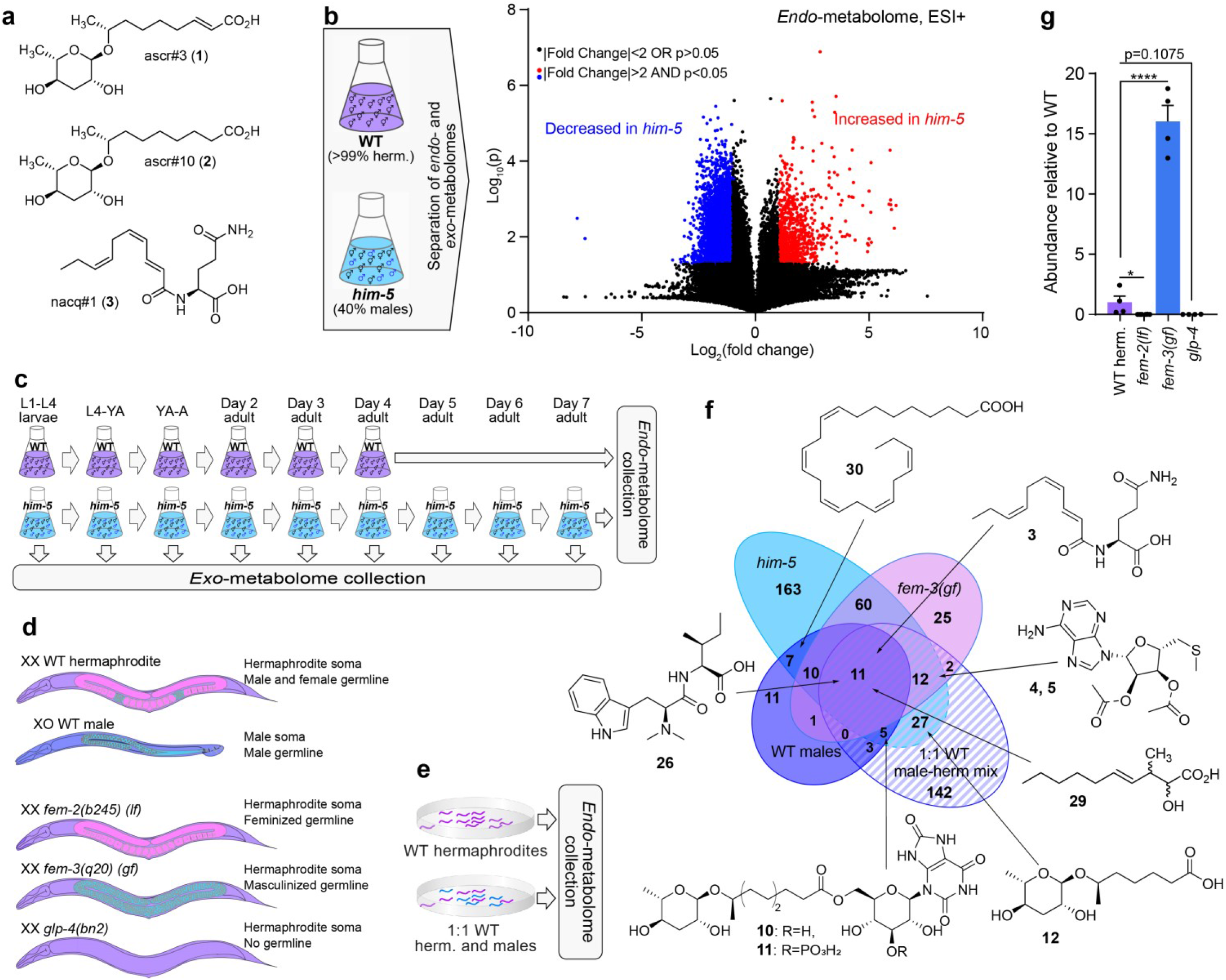
Metabolomic profiling of male *C. elegans*. **a.** Known sex-biased metabolites in *C. elegans*. Whereas ascr#3 (**1**) is more abundant in hermaphrodites, ascr#10 (**2**) and nacq#1 (**3**) are more abundant in males. **b.** Comparison of the metabolomes of WT (N2) and *him-5(e1490)* mutants. Volcano plot shows data obtained in positive ionization (ESI+) mode. **c.** Experimental setup for monitoring metabolism of WT and *him-5* animals with periodic harvesting of the *exo*-metabolome. **d.** Simplified body plans of WT hermaphrodites and males as well as germline-feminized *fem-2(b245) (lf)*, germline masculinized *fem-3(q20) (gf)*, and germline deficient *glp-4(bn2)* mutants. **e.** Schematic of hermaphrodite and male-enriched WT animals grown on plates for metabolomics. **f.** Comparison of male-enriched metabolites detected from *him-5* mutants, *fem-3(gf)* mutants, 1:1 mixtures of WT males and herms, and hand-picked WT males. **g.** Production of nacq#1 (**3**) in germline mutants relative to WT hermaphrodites. *, *p*<0.05; ****, *p*<0.0001.

Studies of ascr#10 and nacq#1 demonstrated that sex-specific metabolites can have major effects on *C. elegans* life history and suggest that identification of additional sex-specific compounds will contribute significantly to a mechanistic understanding of various aspects of *C. elegans* biology. Here, we report a comprehensive untargeted survey of differences between male and hermaphrodite metabolomes, including the roles of male and female germlines in the observed differences. The metabolomic data from this study provide a resource for future detailed exploration of sex-biased metabolic pathways and phenotypes.

## Results

### Approaches toward sex-specific metabolomics

One reason for the relatively scant knowledge of the *C. elegans* male metabolome is the low abundance of males in the wildtype (WT) laboratory strain, N2 Bristol. Under standard laboratory conditions, WT cultures consist almost entirely of self-fertilizing hermaphrodites that carry two X chromosomes, whereas males with a single X chromosome constitute <1% of the population^32^. Although mating between hermaphrodites and males can yield up to 50% male progeny, this form of reproduction occurs primarily on plates and less effectively in liquid culture. The low male frequency and the requirement for plates for mating make it impractical to prepare large samples of animals as needed for in-depth metabolomic analyses.

Therefore, we began our analyses with a comparison of the metabolomes of WT cultures, consisting mostly of hermaphrodites, and *him-5(e1490)* mutant cultures, which contain up to 40% males (Figure 1b). The *him-5(e1490)* animals carry a mutation that increases the likelihood of nondisjunction of X chromosomes during meiosis, resulting in a higher incidence of males, independent of mating^32^. To distinguish metabolites that are primarily excreted, we separately analyzed extracts of the culture media (*exo*-metabolome) and the worm bodies (*endo*-metabolome). Next, we considered that metabolism undergoes profound changes during development and throughout adult life. Elucidation of biological functions and biosynthesis of sex-specific compounds could be aided by the knowledge of the developmental stages during which they are produced. To this end, we examined the *exo*-metabolome of *him-5* cultures by collecting the culture media during different time periods throughout larval development and adulthood (Figure 1c).

Metabolome samples were analyzed by liquid chromatography coupled to a high-resolution mass spectrometer (LC-HRMS). Bioinformatic analysis of the resulting data was conducted using the Metaboseek platform, which provides a comprehensive toolset for comparative metabolomics (see Methods,^8^). Our analysis of the *exo*- and *endo*-metabolomes of WT and *him-5* cultures yielded 110,184 significant features (unique pairs of mass-to-charge (*m/z*) ratios and retention times) including 8,225 features at least two-fold more abundant in the male-enriched *him-5* cultures and 22,538 MS features that are at least two-fold more abundant in WT cultures (Figure 1b). Manual curation to remove isotopes, adducts, and fragments yielded almost 300 unique compounds that are up-regulated in male-enriched *him-5* cultures, and a similar number of compounds enriched in WT relative to *him-5*. Following comparative analysis of the LC-HRMS data, we acquired MS2 fragmentation spectra for all male-enriched metabolites (Supplementary Tables 1-3).

### Prioritization of sex-dependent compounds

Both the germline and somatic tissues differ between the two sexes and could therefore contribute to the detected differences between *him-5* and WT animals. The self-fertile *C. elegans* hermaphrodites produce a cache of ~300 sperm, severalfold fewer than males, before switching to oogenesis^33^. To begin disentangling potential sources of metabolic differences, we took advantage of three conditional mutations that at the nonpermissive temperature alter aspects of germline biology (Figure 1d). The XX individuals (these are normally hermaphrodites) carrying the *fem-3(q20)* gain-of-function (*gf*) allele have a masculinized germline that produces a dramatically increased amount of sperm and no eggs, but do not show overt masculinization in the soma^34^. Conversely, the XX individuals carrying the *fem-2(b245)* loss-of-function (*lf*) allele have a feminized germline that produces eggs but no sperm^35^. The third mutation we used in this study, *glp-4(bn2)*, severely limits germline development (~1% of the wildtype number of germline nuclei), while maintaining an apparently normal soma ^36^, although a likely null allele causes larval lethality^37^. In addition, we included samples of hand-picked WT males and a 1:1 mixture of WT (heat-shock derived) males and hermaphrodites (Figure 1e). These small, plate-derived samples did not permit extensive metabolomic analyses, but were used to verify that many of the compounds enriched in *him-5* and *fem-3(gf)* mutants were likewise enriched in WT males (Figure 1e).

Comparison of metabolomes of *him-5* animals and germline-masculinized *fem-3(gf)* animals revealed that 93 of the 295 *him-5*-enriched compounds were also up-regulated in *fem-3(gf)* cultures (Figure 1f), suggesting that production of this subset of metabolites is associated with the presence of the male germline, whereas production of the remaining 202 *him-5*-enriched compounds may depend on the male soma or other changes in metabolism in *him-5* mutants. The majority of metabolites enriched in *fem-3(gf)* mutants relative to WT animals were also enriched relative to germline-feminized *fem-2(lf)* and germline-less *glp-4* mutants, further supporting that these metabolites depend on the presence of the male germline (Supplementary Tables 1-3). An example of a *him-5*-enriched metabolite that depends on the presence of the male germline is the developmental regulator nacq#1, which was previously shown to be produced in much higher amounts by males relative to hermaphrodites^29^. Quantification of nacq#1 in the germline mutants showed that this metabolite was abundantly produced in masculinized *fem-3(gf)* animals, but absent in feminized *fem-2(lf)* and germline-less *glp-4* animals (Figure 1g). Comparison of the metabolomes of *fem-3(gf), him-5*, and WT cultures revealed 11 additional *him-5*-enriched compounds that were missed in the initial comparison of the *him-5* and WT metabolomes (Supporting Tables 1-3).

For chemical characterization of male-enriched metabolites, we initially prioritized compounds that were consistently associated with the presence of the male germline and increased in *him-5* as well as *fem-3(gf)* compared to WT cultures. Analysis of the molecular formulae and MS2 networks for the resulting set of compounds revealed several different families of likely structurally related metabolites. For in-depth analysis we then selected representative and most abundant members of these families that could also be detected in samples of WT males.

### Male-enriched nucleoside derivatives

Among prioritized male-enriched metabolites in the *endo*-metabolomes, we detected several families of compounds whose molecular formulae and MS2 fragmentation patterns suggested that they represent unusual nucleoside derivatives. These included two isomeric compounds (*m/z* 340.1074, C_13_H_18_N_5_SO_4_^+^, 6.69 and 7.30 min) that fragmented in a nearly identical manner (Figure 2a), with the later-eluting compound being the predominant isomer. MS2 fragments consistent with methylmethylenesulfide (*m/z* 61.0104, C_2_H_5_S^+^) and adenine (*m/z* 136.0608, C_5_H_6_N_5_^+^) suggested that the two compounds represent *O*- acetylated derivatives of *S*-methylthioadenosine (MTA), a common metabolite downstream of *S*-adenosylmethionine (SAM) ^38^. Acetylation of MTA yielded a mixture of two isomers of the same retention time as the natural compounds (acemta#1, **4**, and acemta#2, **5**), confirming their structures (Figures S1, S2).

**Figure 2.**
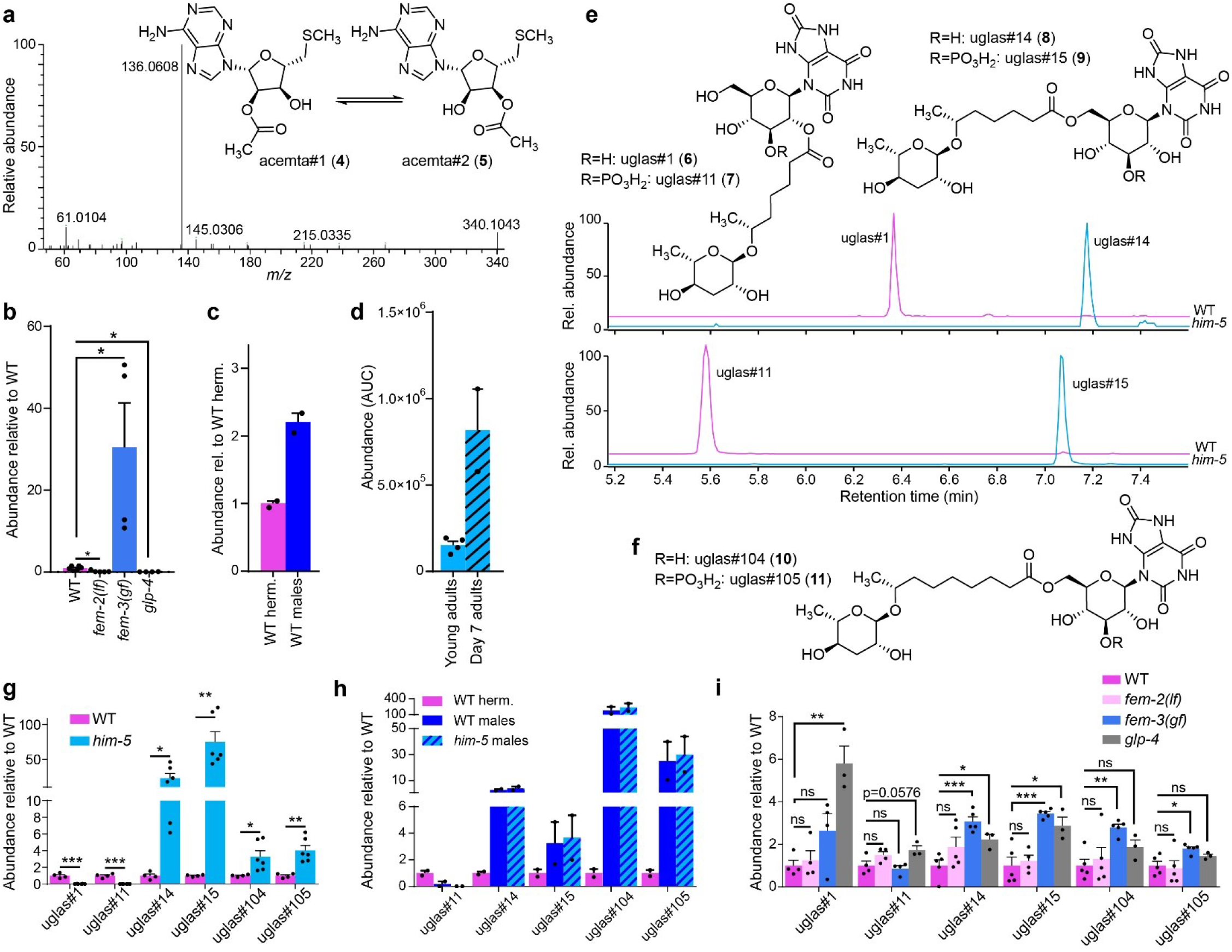
Enrichment of nucleoside-derived metabolites in males. **a.** MS2 fragmentation of acemta#1/2 in positive (ESI+) ionization mode. **b.** Relative abundance of acemta#2 levels in *endo*-metabolome extracts from WT, *fem-2(lf), fem-3(gf)*, and *glp-4* mutants. **c.** Comparison of acemta#2 levels in hand-picked WT males with hand-picked WT hermaphrodites. **d.** Comparison of acemta#2 levels in young (Day 1) and old (Day 7 *him-5* adults. **e.** Sex-specificity of uric acid glucoside derivatives incorporating the 7-carbon side chain ascaroside, ascr#1, and corresponding structures. Shown are ion chromatograms for *m/z* 667.1869 in ESI-ionization mode. **f.** Male-specific uric acid glucoside derivatives incorporating a 9-carbon side chain ascaroside, uglas#104 (**10**) and uglas#105 (**11**). **g-i.** Comparison of uric acid glucoside-containing ascarosides in N2 and *him-5* (**g**), WT hermaphrodites, WT males, and *him-5* males (**h**), and WT and germline mutants (**i**). *, *p*<0.05; **, *p*<0.005; ***, *p*<0.0005; ns, not significant.

Production of acemta#1 and acemta#2 appears to be associated with the presence of the male germline, as the two compounds are undetectable in feminized *fem-2(lf)* and germline-less *glp-4* animals, whereas they are much enriched in masculinized *fem-3(gf)* animals compared to WT hermaphrodites (Figure 2b). acemta#1 and acemta#2 were also enriched in samples of hand-picked males (Figure 2c). In contrast to ascaroside pheromones and nacq#1^22,29^, these nucleoside derivatives accumulate primary in the worm body, and their abundance is increased in older worms (Figure 2d). Notably, the abundance of the putative precursor of acemta#1/2, MTA, is not significantly increased in *him-5* or *fem-3(gf)* mutants (Figure S3).

In addition to acemta#1 and acemta#2, we detected a family of compounds whose MS2 spectra suggested that they represent hexose-based nucleosides, including a series of putative uric acid derivatives. Molecular formulae and MS2 spectra of the two most dramatically enriched (75- and 23-fold over WT in the *him-5 endo*-metabolome) members of this family (*m/z* 587.2211, C_24_H_36_N_4_O_13_ and *m/z* 667.1880, C_24_H_37_N_4_PO_16_) suggested that they represent isomers of the recently described uglas#1 (**6**) and uglas#11 (**7**), two gluconucleosides incorporating uric acid and the ascaroside ascr#1 (Figure 2e,f) ^39^. However, uglas#1 and uglas#11, which bear an ascr#1 moiety in the 2-position of the glucose, had significantly earlier retention times than their male-enriched isomers and were virtually absent in *him-5* mutants. Based on the recent identification of a series of 6’-*O*-acylated glucosides from *C. elegans*^40,41^, we hypothesized that in the male-enriched isomers of uglas#1 and uglas#11 the ascaroside moiety is attached to the 6-position of the uric acid glucoside. Comparison of MS2 spectra and retention times of a synthetic sample of the 6’-*O*-substituted isomer of uglas#1, named uglas#14 (**8**), established that the male-enriched isomer is in fact uglas#14, and thus the male-enriched isomer of uglas#11 was assigned as its phosphorylated derivative, uglas#15 (**9**, Figures 2f, S4, S5). uglas#14 and uglas#15 were accompanied by two compounds with analogous fragmentation patterns (Figures S6, S7), which appeared to incorporate the 9-carbon sidechain ascaroside, ascr#10, instead of the 7-carbon sidechain ascr#1. These compounds, named uglas#104 (*m/z* 615.2531, C_26_H_40_N_4_O_13_, **10**) and uglas#105 (*m/z* 695.2189, C_26_H_41_N_4_PO_16_, **11**), were roughly three- to four-fold enriched in *him-5 endo*-metabolome compared to WT cultures, whereas uglas#14 and uglas#15 were enriched 20-60-fold (Figure 2g). Biosynthesis of the 2’-*O*-acylated uglas#1 and uglas#11 has been shown to require the carboxylesterase *cest-1.1*, which mediates attachment of the ascaroside to the 2’-hydroxy of uric acid gluconucleosides^40^. Production of the male-upregulated 6’-*O*-acylated uglas-family metabolites is not *cest-1.1*- dependent and does not require any of the other so-far characterized *cest* homologs^40,41^, suggesting that a different carboxylesterase is involved.

All four male-enriched uglas#-family metabolites that were enriched in the *endo*-metabolomes of large *him-5* cultures were also enriched in small hand-picked samples of N2 and *him-5* males (Figure 2h). As is the case with the larger *him-5* cultures, production of the 2’-*O*-acylated uglas#11 is significantly decreased or even abolished in hand-picked samples of males relative to WT hermaphrodites, whereas abundances of the 6’-*O*-acylated isomer, uglas#15, and of uglas#104/105 were dramatically increased (Figure 2h). Similar to acemta#1 and acemta#2, the uglas#-family metabolites are predominantly retained within the worm body, and thus our experimental set-up for determining changes of abundances during the adult lifespan (see Figure 1c) allowed only for comparison of abundances in young adults and 6 day-old adults, which indicated a modest increase of uglas#104 and uglas#105 in day 6 adult worms (Figure S8). In contrast to other male-enriched metabolites, including nacq#1 and acemta#1/2, production of the male-enriched uglas#-family metabolites was not strictly dependent on the presence of a male (or female) germline and instead appears to be related to the male soma. Abundances of the male-specific uglas#-family metabolites were not reduced in feminized *fem-2(lf)* animals relative to WT, were only slightly increased in masculinized *fem-3(gf)* animals, and were either slightly enriched or not changed in germline-less *glp-4* animals relative to WT (Figure 2i).

### Male-enriched ascaroside derivatives

Several recent studies have shown that the ascaroside ascr#10, the first reported male-enriched *C. elegans* metabolite^22^, has broad effects on hermaphrodite development and physiology^24–29^. The original work reporting increased levels of ascr#10 in males was based on the analysis of mixed-stage *him-5* cultures, containing animals at different larval stages and ages of adulthood. In these *him-5* cultures, ascr#10 was found to be about 3-fold upregulated^22^. Our *exo*-metabolome survey throughout larval development and adulthood revealed that only trace amounts of ascr#10 are produced during the four larval stages (L1-L4), and that production increases dramatically in young adults. In *him-5* cultures, secretion of ascr#10 increased until day 4 of adulthood and was roughly tenfold higher than in WT cultures from young adulthood through the course of the experiment (Figure 3a,b).

**Figure 3.**
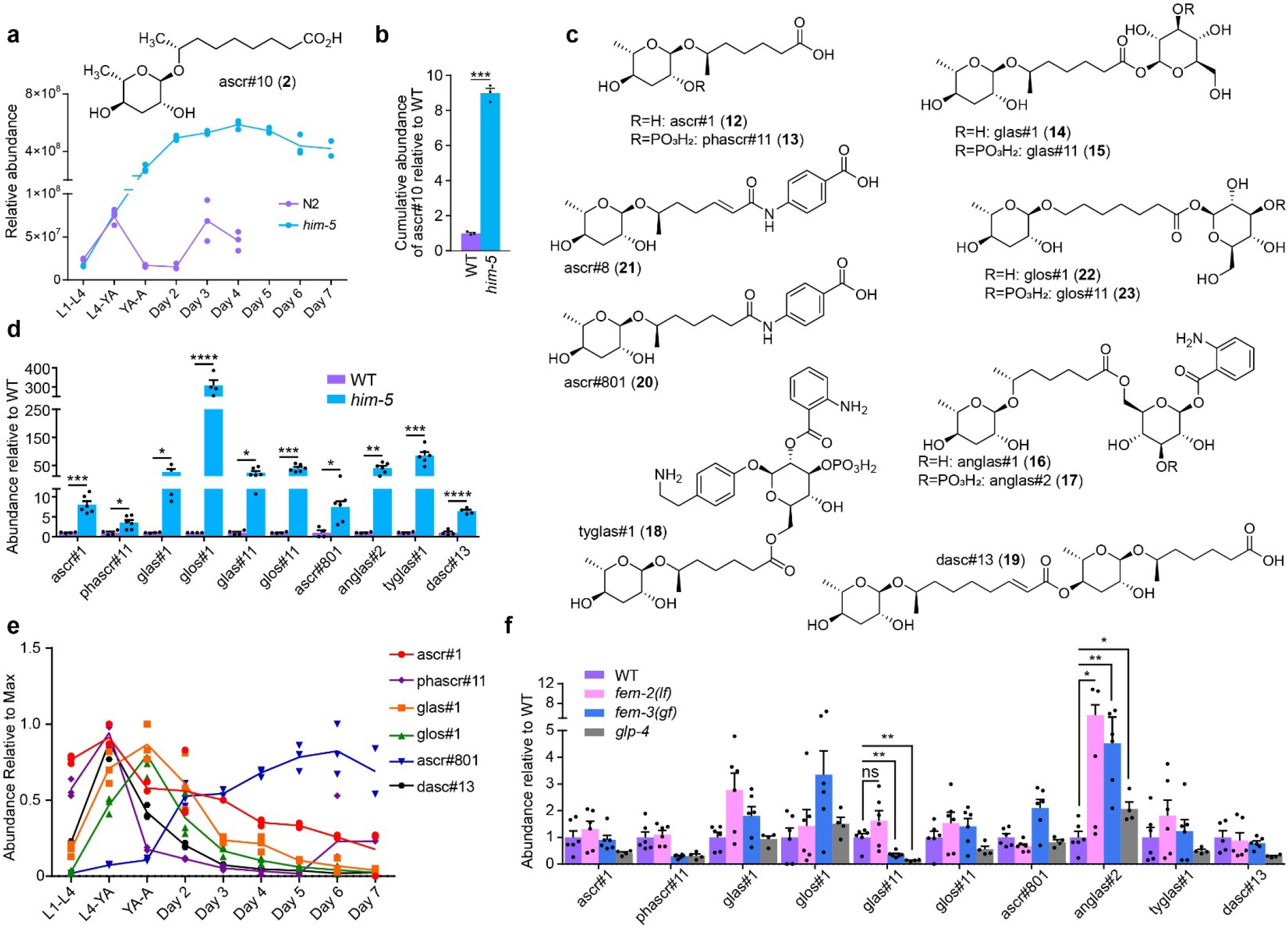
Enrichment of ascr#10 and ascr#1 derivatives in males. **a.** Developmental stage-dependent production of ascr#10 (**2**) in WT and *him-5*. **b.** Cumulative production of ascr#10 by WT and *him-5* from the L1 larval stage through Day 4 of adulthood. **c.** Chemical structures of identified male-enriched ascr#1 derivatives. **d.** Comparison of ascr#1 derivatives in the *exo*-(ascr#1, phascr#11, glas#1, glos#1, ascr#801, and dasc#13) and *endo*-metabolomes (glas#11, glos#11, anglas#2, and tyglas#1) of WT and *him-5* animals. **e.** Development-dependent production of male-enriched ascr#1 derivatives in *him-5* animals. **f.** Comparison of ascr#1 derivatives in WT and germline mutants. *, *p*<0.05; **, *p*<0.005; ***, *p*<0.0005; ****, *p*<0.0001; ns, not significant.

In addition to ascr#10, our comparative analysis of *him-5* and WT cultures revealed significantly increased amounts of the ascaroside ascr#1 (**12**) (Figure S9) and a series of structurally more complex ascaroside derivatives, primarily in the *him-5 endo*-metabolome (Figure 3c,d). These include the phosphorylated phascr#11 (**13**) and the previously described glucoside glas#1 (**14**)^42,43^ which were identified based on analysis of the MS2 spectra (Figures S10 and S11) and comparison of retention times. MS2 spectra further allowed proposing structures for several additional compounds (Figures S12-S16), including a phosphorylated derivative glas#11 (**15**), the previously described conjugate of ascr#1 with anthranilic acid glucosides, anglas#1 (**16**)^11^ and anglas#2 (**17**), a putative ascr#1-derivative of tyramine glucoside^41^, tyglas#1 (**18**), a putative dimeric ascaroside^44^, dasc#13 (**19**), and a conjugate of ascr#1 and *p*-amino benzoic acid, ascr#801 (**20**), representing a dihydro derivative of the known dauer pheromone component ascr#8 (**21**)^45^. The glucosides glas#1 and glas#11 were accompanied by later eluting isomers with near identical MS2 spectra (see Figures S11, S12) that likely represent glos#1 (**22**) and glos#11 (**23**), which feature an ω-oxygenated fatty acid chain instead of the (ω-1)-oxygenated in ascr#1, glas#1, and glas#11.

Excretion of most *him-5* upregulated ascarosides, including ascr#1 itself, peaked early in life, either around the L4-young adult molt or the first day of adulthood (Figure 3e). Similarly, ascr#1-derivatives primarily retained in the worm body were more abundant in young adults than in day-6 adults (Figure S17). An interesting exception is ascr#801, which was barely produced early in life and was most abundant in older animals (~day 6 of adulthood). Similar to the uglas#-family nucleosides described above, abundances of most *him-5* enriched ascr#1 derivatives did not correlate with the presence of a male germline (Figure 3f). However, in contrast to the uglas#-family of compounds, most of the *him-5* enriched ascr#1 derivatives were not enriched or could not be detected in samples of pure males (Figure S18), suggesting that production of ascr#1 derivatives may be upregulated in response to male-hermaphrodite interactions.

### An unusual male-enriched dipeptide

The comparison of the *exo*- and *endo*-metabolomes of *him-5* and WT animals further revealed a small family of male-enriched metabolites whose MS2 spectra suggested they represent dipeptide derivatives. The MS2 spectrum of the most abundant member of this compound family (medip#1, *m/z* 344.1978, C_19_H_26_N_3_O_3_^-^, 7.94 min) showed fragments suggesting the presence of putative indole (*m/z* 116.0512, C_8_H_6_N^-^), and (iso)leucine (*m/z* 130.0872, C6H12NO2^-^) moieties, as well as fragments consistent with loss of methylindole (*m/z* 215.1413, C_10_H_19_N_2_O_3_^-^), loss of CO_2_ and dimethylamine (*m/z* 255.1516, C_16_H_19_N_2_O-), and loss of CO_2_ (*m/z* 300.2102, C_18_H_26_N_3_O^-^) (Figure 4a). This fragmentation pattern suggested a dipeptide consisting of *N*,*N*-dimethyltryptophan and either leucine or isoleucine. To confirm these assignments and to differentiate between incorporation of leucine or isoleucine, we synthesized the two candidate compounds (Figure 4b). Dimethylation of tryptophan with formaldehyde and sodium cyanoborohydride yielded *N*,*N*-dimethyltryptophan (**24**), which was then conjugated to either *O*-tBu-leucine or *O*-tBu-isoleucine, followed by deprotection with trifluoroacetic acid^10^. Comparison of retention times and MS2 spectra of isomers **25** and **26** showed that this male-enriched metabolite represents *N,N*- dimethyltryptophan-isoleucine, that we named medip#1 (**26**, Figure 4c, S19). medip#1 is accompanied by smaller amounts of the corresponding *N*-oxide, medip#2 (**27**, Figure S20), and the monomethylated derivative medip#3 (**28**).

**Figure 4.**
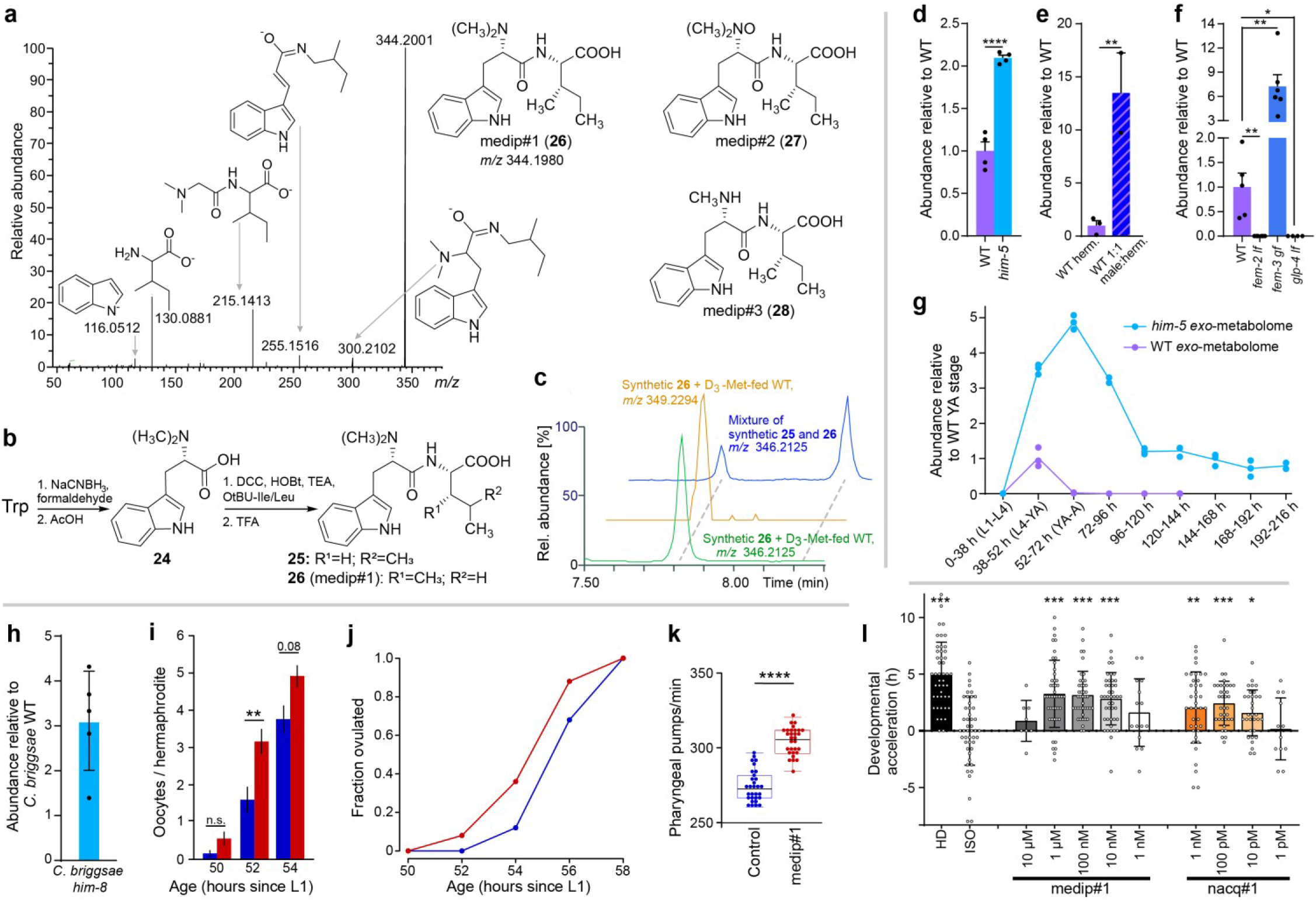
Identification, germline dependence and biological activity of dipeptide medip#1. **a.** Structures of medip#1-3 (**26**-**28**) and MS fragmentation of medip#1(**26**) in ESI-mode. **b.** Synthesis of medip#1 and related molecules. **c.** HPLC-MS traces of synthetic medip#1 with natural medip#1 from D_3_-methionine-fed animals. **d,e,f.** Comparison of medip#1 levels in WT and *him-5* animals (**d**), in WT 1:1 male:hermaphrodite cultures and pure WT hermaphrodite cultures (**e**), and in WT and germline mutants (**f**). **g.** Developmental stage-dependent production of medip#1 in WT and *him-5* animals. **h.** Upregulation of medip#1 in the *exo*- metabolome of *C. briggsae him-8* cultures. **i.** Faster acquisition of oocytes in hermaphrodites exposed to medip#1 (red) compared to paired controls (blue). **j.** Hermaphrodites exposed to medip#1 (red) ovulate earlier than controls (blue).**k.** Faster pharyngeal pumping in hermaphrodites exposed to medip#1. **l**. Time point of first egg laying of isolated worms on different concentrations of medip#1 and nacq#1 compared to untreated isolated worms (ISO, control) and grouped worms (high density, HD). *, *p*<0.05; **, *p*<0.005; ***, *p*<0.001; ****, *p*<0.0001.

Feeding of methyl-D3 methionine to *C. elegans* resulted in robust D3- and D6-labeling of medip#1-3 (Figure S21), indicating that methylation of the tryptophan moiety in these dipeptides proceeds via an *S*-adenosyl methionine- (SAM-)dependent methyltransferase. Notably, the free amino acid *N,N*-dimethyltryptophan (**24**), which we detected in *exo*- and *endo*-metabolome samples of WT and *him-5* mutants, remained unlabeled in the D3-methionine feeding experiment (Figure S22). In contrast to medip#1-3, dimethyltryptophan was not enriched in *him-5* samples relative to WT and was also detected in extracts of the *E. coli* OP50 bacteria used as food, suggesting a bacterial origin of this compound in the *C. elegans* metabolome samples. These results suggest that medip#1-3 are derived from methylation of a possibly ribosomally produced precursor peptide in *C. elegans*, rather than from a peptide-forming reaction using *N*,*N*-dimethyltryptophan.

medip#1 was enriched not only in cultures of *him-5* animals (Figure 4d), but also in cultures with equivalent numbers of males and hermaphrodites (Figure 4e), further confirming that this compound is enriched in males. Production of medip#1 was greatly increased in masculinized *fem-3(gf)* animals, but undetectable in feminized *fem-2(lf)* and germline-less *glp-4* animals, indicating its production is associated with the presence of a male germline (Figure 4f). medip#1 is produced abundantly in *him-5* cultures for several days after reaching the adult stage, whereas WT hermaphrodite cultures produced modest amounts of this compound and only transiently during the late larvae/young adult stage (Figure 4g).

### Conservation and biological activity of medip#1

Next, we asked whether male-enriched production of the metabolites identified in the preceding sections is conserved in other nematodes. A commonly used reference point for comparisons with *C. elegans* is *C. briggsae*, a member of the same genus that also typically reproduces by self-fertilization. Similar to *C. elegans him-5*, cultures of *C. briggsae CBR-him-8* mutants produce increased numbers of males (up to 30% in our hands^46^). Comparing the *exo*- and *endo*-metabolomes of *C. briggsae CBR-him-8* cultures and the *C. briggsae* WT strain AF16 revealed a similar number of differential features as in our comparison of *C. elegans* WT (N2) and *CEL-him-5* metabolomes. However, most male-enriched metabolites we identified in *C. elegans* could not be detected or were not enriched in *C. briggsae* males, consistent with other recent studies demonstrating that the metabolomes of nematodes are highly species-specific^13^. Notable exceptions include the previously reported nacq#1^29^ as well as the dipeptide medip#1 (**26**, Figure 4h), which was enriched in the *CBR-him-8 exo*-metabolome to a similar extent as in *CEL-him-5*.

Given that male-enriched production of medip#1 is conserved, we selected this compound for further biological evaluation. medip#1 is much more abundant in the *exo*- than in the *endo*-metabolomes in both *C. elegans* and *C. briggsae*, indicating that it is preferentially excreted by males and may serve as a chemical signal, potentially eliciting responses in hermaphrodites. We found that hermaphrodite larvae reached morphologically-defined adulthood faster on plates conditioned with synthetic medip#1 (Figure S23a). This acceleration is due to shortening of the last larval stage (L4) because developmental progression of earlier larval stages is not affected by medip#1 (Figure S23b). Faster development of somatic tissues was accompanied by a faster maturation of the oogenic germline as medip#1-exposed hermaphrodites acquired oocytes faster (Figure 4i) and started to ovulate earlier (Figure 4j). Development of the soma and the germline are energetically demanding. Plausibly to sustain accelerated sexual maturation, medip#1 increases the rate of food consumption (Figure 4k). The result of the totality of physiological changes induced by medip#1 in hermaphrodites is the earlier onset of reproduction (Figure 4l), which may confer a competitive advantage under specific conditions during the boom-and-bust cycles characteristic of the ephemeral environments where *C. elegans* dwell^14^.

## Discussion

Our comparative metabolomic analyses of *C. elegans* males and hermaphrodites revealed several hundred significantly male-enriched metabolites, for which we provide MS and MS2 data as a basis for future studies toward their detailed biochemical and functional characterization. It is likely that this inventory is still largely incomplete, as we used stringent peak intensity cutoffs for our analysis that may have excluded less abundant or less consistently produced metabolites. Furthermore, chromatographic conditions optimized specifically for the detection of very polar and very non-polar metabolites (e.g. lipids) will almost certainly reveal additional male-upregulated compounds. Full chemical characterization of the large number of newly detected metabolites presents a major challenge, since structure elucidation cannot yet be automated and still relies on supervised one-by-one approaches. For this study, we therefore selected representative, abundant members of several families of male-enriched compounds for detailed characterization, which uncovered a wide range of unusual chemical structures integrating building blocks from diverse metabolic pathways.

Our analysis revealed that production of a subset of male-enriched metabolites was correlated with the presence and sexual identity of the germline (Figure 5a). Biosynthesis of the dipeptides medip#1-3 and the acetylated MTA derivatives acemta#1/2 was abolished in germline-feminized *fem-2(lf)* and germline-less *glp-4* animals and substantially increased in germline-masculinized *fem-3(gf)* animals. Consistent with requirement of a male germline for their production, medip#1-3 and acemta#1/2 were absent in developing worms until germline maturation during the young adult stage (Figure 4f). Several additional families of male-enriched metabolites not discussed in the preceding sections were also associated with the presence of a male germline. This includes a stark increase in the production of very long-chain polyunsaturated fatty acids, e.g., tetracosapentaenoic acid (**30**, Figure S24), which may be derived from the marked upregulation of two fatty acid elongases, *elo-4* and *elo-7*, in males^16^. Additional male germline-dependent metabolites include the recently reported β-methyldecanoic acid derivatives, bemeth#2 (**29**) and bemeth#3^8^, as well as a number of putative modular glucosides and other compounds whose structures have not yet been elucidated (Tables S1-3).

**Figure 5.**
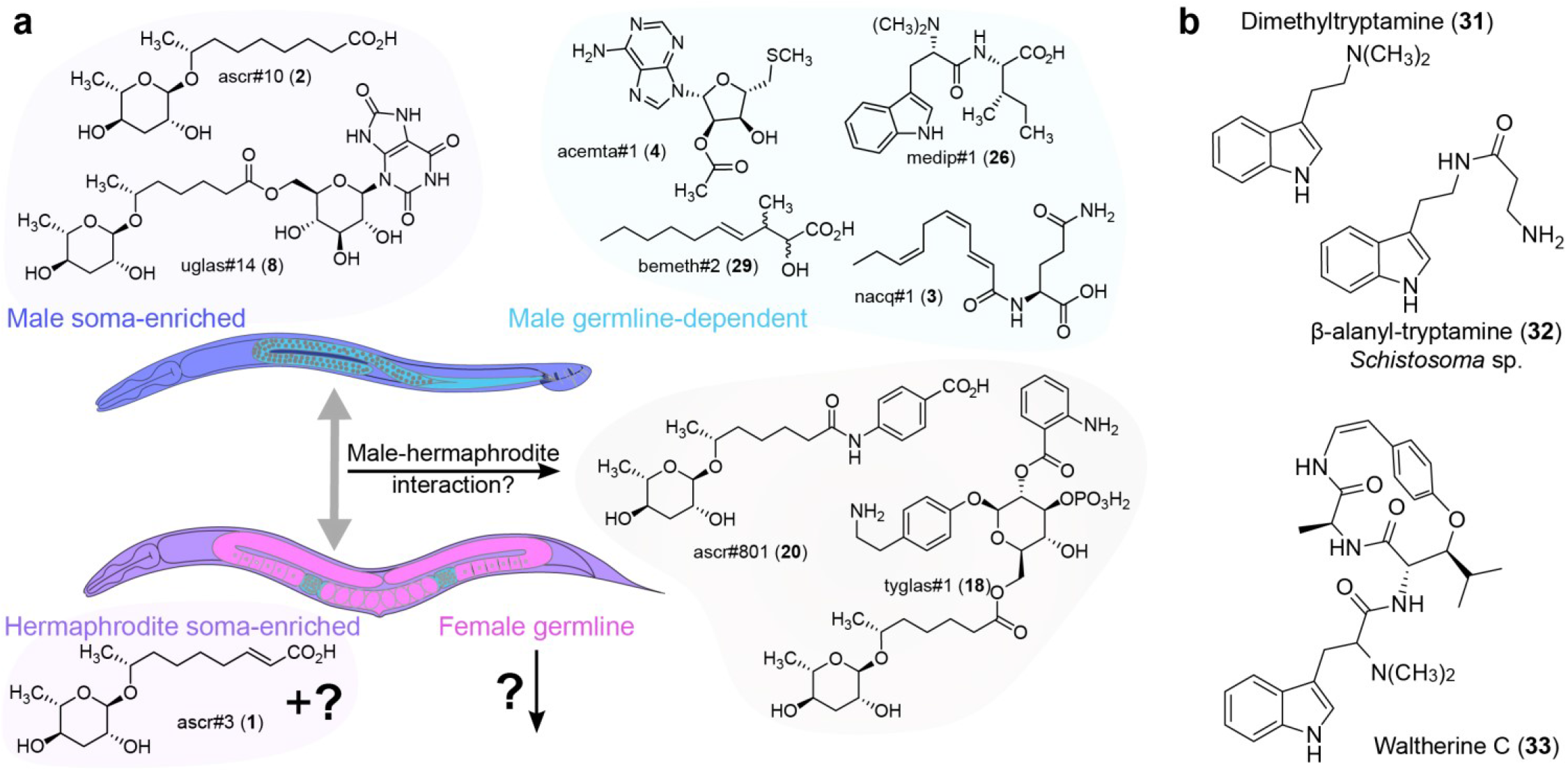
**a.** Soma- and germline dependence of the identified male-enriched metabolites and related small molecules identified from natural sources. **b.** Structures of dimethyltryptamine and dimethyltryptophan derivatives from other animals and plants.

In contrast, production of several other families of male-enriched metabolites does not depend on the presence of a male germline. For example, all male-enriched uric acid glucosides (e.g. uglas#14 and uglas#104) can still be detected in germline-feminized *fem-2(lf)* and germline-less *glp-4* animals. Notably, production of uglas#14 and uglas#104, which are more than 10-fold enriched in WT or *him*-5 males relative to WT hermaphrodites, was only slightly increased in germline-masculinized *fem-3(gf)* animals and didn’t differ significantly between *fem-3(gf)* and germline-less *glp-4* animals. *fem-3(lf)* animals have a largely hermaphrodite-like soma, and thus upregulation of uglas#-family metabolites appears to require a male soma (Figure 5a).

Yet another pattern is exemplified by the diverse family of *him-5*-enriched ascr#1 derivatives (Figure 3). Although production of these compounds was dramatically increased in either the *exo*- or *endo*-metabolomes of male-enriched *him-5* cultures, most could not be detected in samples of pure males, except for ascr#1, which was not upregulated. Thus, increased levels of ascr#1 and ascr#1-incorporating modular metabolites (Figure 3c) were observed only in cultures containing both males and hermaphrodites, but not in samples of pure WT or *him-5* males, suggesting that increased production of these compounds may result from interactions between the two sexes (Figure 5a).

Given their elaborate structures and sex-specific production, it seems likely that many of the identified metabolites serve dedicated biological functions, e.g. as signaling molecules. Previously reported male-excreted compounds in *C. elegans* affect aspects of behavior^22,24^, germline biology^26–28^, and the rate of larval development^27,29^. Among the newly identified metabolites we selected the dipeptide medip#1 for further study, because its male-enriched production is conserved in *C. briggsae* and since it is abundantly produced in a male germline-dependent manner. In addition, we were intrigued by the similarity of its chemical structure to that of other potently active biomolecules, e.g. dimethyltryptamine^47,48^ (**31**, Figure 5b). We found that at physiologically relevant concentrations medip#1 accelerates larval development of hermaphrodites by shortening the duration of the last larval stage. Another male-enriched compound, nacq#1, similarly accelerates the last larval stage^29^, consistent with the earlier finding that samples of the entire male *exo*-metabolome have the same effect^27^. Whether nacq#1 and medip#1, despite their dissimilar chemical structures, accelerate development via a shared mechanism remains to be elucidated. Both somatic and germline effects of medip#1 promote the male reproductive strategy of facilitating sexual maturation of potential mating partners. Similar observations in other species, including mammals^49^, suggest that this may be a universal feature of social communication in animals. Interestingly, a dipeptide-like compound, β-alanyl-tryptamine (**32**, Figure 5b), was recently identified from male *Schistosoma mansoni* and, as medip#1, shown to stimulate female development^50^. Like schistosomal β-alanyl-tryptamine, medip#1 could be derived from a non-ribosomal peptide synthetase (NRPS). However, the *C. elegans* genome features only one NRPS-like gene, which has been shown to be involved in the biosynthesis of a different metabolite^51^. Moreover, the finding that dietary dimethyltryptophan is not incorporated into medip#1 may suggest a ribosomal origin. Cyclopeptides of likely ribosomal origin that include dimethyltryptophan have been described from plants, e.g. waltherine (**33**)^52^.

Our results demonstrate that the male identity of the germline and the soma can regulate production of distinct sets of metabolites. Similarly, it can be expected that the hermaphrodite germline and soma contribute metabolites that are not, or only to a lesser extent, produced in males, and data deposited with this study provide a starting point for their analysis. It should be noted that correlation of a metabolite with the presence of the male or female germline may not necessarily imply that biosynthesis of that metabolite occurs, entirely or in part, within the germline itself. Alternatively, germline-dependent metabolites may result from activation or priming of somatic biosynthetic pathways by signals from the germline. Identification of genes involved in the biosyntheses of the different germline-dependent compound families will be an important next step toward uncovering their tissue origin. Taken together, our work highlights the power of untargeted comparative metabolomics in a genetically tractable model system to decipher the role of sex and the contributions of signals from different organ systems in shaping animal metabolomes.

## Acknowledgements

This research was supported in part by the National Institutes of Health (R35GM131877 and U01GM110714 to FCS.) and the Howard Hughes Medical Institute (Faculty Scholar grant to FCS). EZA and IR were supported in part by an NSF grant (IOS-1755244) to IR. Some strains used in this work were provided by the CGC, which is funded by the NIH Office of Research Infrastructure Programs (P40 OD010440). We thank Diana Carolina Fajardo Palomino and Gary Horvath for technical support and Ivan Keresztes and David Kiemle for assistance with NMR spectroscopy.

## Author Contributions

FCS supervised the study. RNB, ABA, BWF, SSL, and FCS designed the study. RNB, ABA, EZA, BWF, BJC, AHL, OP, CJW, and AC performed chemical and biological experiments. RNB, IR, and FCS wrote the paper with input from all authors.

## Data availability

All data analyzed during this study are included in the manuscript and supporting files. MS and MS/MS data are available at GNPS/MassIVE under accession number: (will be uploaded after initial review). See attached MS Data Inventory for file names and sample identities (attached as a separate file).

