## Supplementary Information for "Sex-specificity of the *C. elegans* metabolome"

**Contents**

Methods 2

Supplementary Figures 8

Supplementary Tables 31

NMR Spectra 44

Supplementary References 50

**Methods**

**Nematodes and Bacterial Strains.** Unless otherwise indicated, worms were maintained at 20 °C on Nematode Growth Medium (NGM) petri dish plates seeded with *E. coli* OP50 obtained from the *Caenorhabditis* Genetics Center (CGC)[1]. Populations were synchronized by alkaline hypochlorite treatment of gravid hermaphrodites. Isolated eggs were rocked overnight in M9 buffer at 20 °C to yield synchronized, starved L1 larvae[2]. For animals that were sensitive to alkaline hypochlorite treatment, mixed-stage animals were grown at 15 °C and then washed off the plate with 20 mL M9 buffer which was collected in 50 mL conical tubes. Tubes were left for 10 minutes at room temperature, after which all non-L1 stages settled at the bottom of the tube. The top 15 mL, containing L1 animals, was transferred to a new conical tube, rinsed twice with M9 buffer, and then rocked overnight in M9 buffer at the non-permissive temperature (25 °C) prior to experiments. The following strains were used for metabolomics experiments: wildtype Bristol N2, CB4088 *him-5(e1490)*, DH245 *fem-2(b245)*, JK816 *fem-3(q20)* (gain-of function), CB3844 *fem-3(e2006)* (loss-of-function), SS104 *glp-4(bn2)*, CB4037 *glp-1(e2141)*, JK569 *mog-3(q74)* PS7922 *cest-5.1(syb1131),* FCS51 *cest-5.1(syb1131); him-5(e1490)*, PHX3933 *cest-5.1(syb1131);cest-5.2(syb3933)*, GR1395 *mgIs49 [mlt-10::GFP-pest; ttx-1::GFP].*

***C. elegans* Liquid Cultures.** Animals were grown at a density of 3 worms / 1 µL at the indicated temperatures in S-Complete media supplemented with concentrated OP50. When animals reached adulthood, the cultures were settled as described to separate adults from any offspring. The adult animals were rinsed three times with M9 and once with water to remove residual OP50 and salts. The supernatant containing L1 larvae was centrifuged at 1,000 x g for 1 minute to remove L1 larvae and residual OP50. Adult worm pellets and conditioned media samples were snap frozen and stored at -20 °C until extraction.

**50:50 Male:Hermaphrodite Samples.** Four N2 animals were placed on 3.5 cm NGM plates seeded with OP50 and grown at 20 °C to the L4 stage. The worms were then incubated at 30 °C for 5 hours, then transitioned back to 20 °C and allowed to reproduce. Males resulting from heat shock were then picked to new plates to mate with N2 hermaphrodites to yield populations of approximately 50% males.

***C. elegans* Plate-Based Cultures.** Mixed-stage animals were grown on 10 cm NGM plates at 20 °C until nearly all of the food was depleted, at which point animals were rinsed off the plate with M9 and settled as described above. Isolated L1 larvae were transferred to new 10 cm NGM plates seeded with OP50. Worms were grown at 20 °C for 72 hours to yield a synchronized population of 1-day-old adults which were collected by washing the plate with M9. The animals were centrifuged at 1,000 x g for 1 minute and the supernatant was transferred to a new tube, snap frozen, and stored at -80 °C. The worms were rinsed twice with M9 and once with water, then frozen and stored at -80 °C until extraction.

**Hand-Picked Males and Hermaphrodites.** N2 and *him-5* animals were grown on 10 cm NGM plates seeded with OP50 as described above. After 72 hours, 2000 N2 hermaphrodites, 2000 N2 males, and 2000 *him-5* males were picked with a platinum-tipped worm pick, rinsed twice with M9 and once with water, pelleted, frozen, and stored at -80 °C until extraction.

**Developmental and Adult Time Course.** Approximately 100,000 synchronized L1 larvae obtained by alkaline hypochlorite treatment (described above) were grown at a concentration of 3 worms / 1 µL in S-Complete medium supplemented with concentrated OP50 at 20 °C. After 38 hours when the majority of worms were in the $4^{th}$ larval stage (L4), the cultures were transferred to 50 mL conical tube and allowed to settle for 10 minutes, as described above. The top 25 mL of the culture medium was transferred to a new conical tube, centrifuged at 1,000 x g for 1 minute, and then snap frozen. The animals were rinsed twice with M9 buffer, once with S-Complete buffer, and then resuspended at the same concentration in S-Complete medium and supplemented with OP50. The same protocol was followed, and conditioned medium was harvested again at 52 hours (worms in young adult stage), at 72 hours (gravid adult stage), 96 hours (day 1 adults), 120 hours (day 2 adults), and 144 hours (day 3 adults). The settling protocol was essential to separate newly hatched larvae from the aging adults; many larvae were present in the culture beginning at the 72 hour-timepoint. At the final harvest, the aged adults were pelleted, rinsed three times with M9 buffer and once with water, and then snap frozen. Developmental profiling of *him-5(e1490)* cultures was performed using the same protocol but extended by two days. Conditioned medium was collected at 38, 52, 72, 96, 120, 144, 168, and 192 hours after experiment start, and the aged adults were collected at the final time as described.

**Procedure for Liquid Growth.** N2 and *him-5* L1 animals were synchronized, and 100,000 animals were grown in S-Complete at 25 °C as described above. After 56 hours the culture was settled, the pellet rinsed three times with M9 and once with water, the supernatant was centrifuged to remove L1s and OP50, and both pellet and supernatant were frozen at -80 °C.

**Procedure for $D_3$-Methyl-Methionine Feeding.** Synchronized *him-5(e1490)* L1 larvae were prepared by alkaline hypochlorite treatment as described, and five separate cultures of 70,000 animals each were grown in S-Complete media supplemented with concentrated OP50. Cultures were grown without addition or were supplemented with L-Methionine (Sigma-Aldrich M9625) or L-$D_3$-Methyl-Methionine (Cambridge Isotope Laboratories DLM-431) to achieve a final concentration of 10 mM. Two isotopic labeling experiments were performed in which animals were either supplemented beginning at the L4 stage (after 38 hours in culture) or at the beginning of the experiment.

**Extraction Procedure.** Frozen worm pellets (*endo*-metabolome) and conditioned media (*exo*-metabolome) were lyophilized, and pellets were homogenized with a tissue grinder. Media and pellets were extracted in 30 mL and 13 mL pure methanol, respectively, for 16 hours with shaking. The methanol extracts were separated from insoluble material by centrifugation, dried *in vacuo*, and resuspended in 50 µL of methanol for plated cultures and 150 µL of methanol for liquid cultures.

**Lipid Extraction Procedure.** Frozen pellets were lyophilized and homogenized as above, then extracted with 12 mL of a 9:1 ethyl acetate:ethanol mixture for 16 hours with stirring. The organic extracts were treated as the methanol extracts above and resuspended in 150 µL of ethanol for analysis by HPLC-MS.

**UHPLC-HRMS.** Liquid chromatography was performed using a Vanquish Horizon UHPLC controlled by Chromeleon software (ThermoFisher Scientific) coupled to an Orbitrap Q-Exactive HF high-resolution mass spectrometer controlled by Xcalibur software (ThermoFisher Scientific) or a Dionex Ultimate 3000 UHPLC coupled to an Orbitrap Q-Exactive high-resolution mass spectrometer controlled by the same software. UHPLC separation was achieved using a Thermo Hypersil GOLD C18 column (2.1×150 mm 1.9 µm particle size) maintained at 40 °C. Solvent A: 0.1% formic acid in water; solvent B: 0.1% formic acid in acetonitrile. A/B gradient started at 1% B for 3 min after injection and increased linearly to 98% B at 20 min, followed by 5 min at 98% B, then back to 1% B over 0.1 min and finally held at 1% B for an additional 2.9 min.

For lipid extracts, reversed-phase post-column ion-pairing chromatography was performed using a Vanquish Horizon UHPLC coupled to an Orbitrap Q-Exactive HF as above. Solvent A: 0.1% ammonium acetate in water; solvent B: acetonitrile. A/B gradient started at 5% B for 3 min after injection and increased linearly to 98% B at 20 min, followed by 5 min at 98% B, then back to 5% B over 0.1 min and finally held at 5% B for an additional 2.9 min. A second pump (Dionex 3000) controlling a solution of 800 mM ammonia in methanol was run at a constant flow rate of 0.015 mL/min for the duration of the method and mixed via micro-splitter valve (Idex #P-460S) with the eluate line from the column.

Mass spectrometer parameters: spray voltage (-3.0 kV, +3.5 kV), capillary temperature 380 °C, probe heater temperature 400 °C; sheath, auxiliary, and sweep gas 60, 20, and 2 AU, respectively. S-Lens RF level: 50, resolution 120,000 at *m/z* 200, AGC target 3E6. Each sample was analyzed in negative and positive electrospray ionization modes. Parameters for MS/MS (dd-MS2): MS1 resolution: 60,000, AGC Target: 1E6. MS/MS resolution: 30,000, AGC Target: 2E5, maximum injection time: 50 ms, isolation window 1.0 *m/z*, stepped normalized collision energy (NCE) 10, 30; dynamic exclusion: 5 seconds, top 10 masses selected for MS/MS per scan.

**Metaboseek Analysis**. UHPLC-HRMS data were analyzed using Metaboseek software after file conversion to the mzXML format via MSConvert (v3.0, ProteoWizard)[1,2]. The feature filtration criteria used were a minimum 2-fold increase in strain of interest over control (i.e., *him-5* over N2), a minimum average intensity of 100,000 arbitrary units for the feature in the strain of interest, and a *p*-value less than 0.05 as calculated by two-sided, unpaired *t*-test.

**Manual Integration and Normalization**. For compounds discussed, the area under the curve (AUC) of the compound of interest was manually obtained through integration in Thermo FreeStyle and Qual Browser. Normalization to wildtype (N2) was achieved by dividing the AUC of the compound of interest by the AUC of ascr#3, as detected in the exo-metabolome in ESI- ionization.

**Statistical Analysis of Metabolomics Data.** All *p*-values were calculated by GraphPad Prism via unpaired *t*-tests with Welch correction. All error bars are presented as standard errors of the mean (SEM).

**Conditioning Plates with medip#1.** Concentrated medip#1 was stored in ethanol at -20 °C. Dilutions were made in ethanol. For all treatments, 2 µM medip#1 solution and added to an equal volume of 1:10 dilution of OP50 overnight culture so that 1 µM medip#1 in a total volume of 20 µL was pipetted on the plate. This was quickly absorbed into the plate and a second 20 µL drop of 1:10 dilution OP50 was pipetted over the first drop. Control plates were made with ethanol alone added to OP50 bacteria as above. These plates were incubated overnight at 20 °C and used the following day.

**Developmental Acceleration.** The time point of first egg laying (Figure 4I) was determined as previously described[3]. Acceleration of reaching adult developmental morphology (Figure S23) was determined using the protocol described in[4]. Briefly, after synchronization by hypochlorite treatment, 25 L1 larvae were singled onto control plates and 25 L1 larvae were singled onto plates prepared with 1 µM medip#1 as above. Beginning at 48 hours post release from L1 arrest, worms were monitored every two hours for transition from L4 to young adult. Developmental staging was based on vulval and gonadal morphology [5]. These results were used to calculate the difference in time of development when half the population had reached adulthood between worms exposed to 1 µM medip#1 and their paired controls.

**Census of the Reproductive System.** A census of oocytes in the gonad and embryos in the uterus was performed using an established protocol[6]. Approximately 30 synchronized L1 larvae prepared by alkaline hypochlorite treatment were pipetted onto either control NGM plates or NGM plates prepared with 1 µM medip#1, as described above. Every two hours, beginning at 48 hours release from post L1 arrest, twenty-five worms from each condition were examined on a Leica DM5000B compound microscope. The numbers of oocytes that completely spanned the gonad in both the anterior and posterior arms, as well as the number of fertilized embryos in the uterus, were counted. In addition, the fraction of worms that had ovulated at least once was noted.

**Pharyngeal Pumping Rate.** N2 hermaphrodites were synchronized by alkaline hypochlorite treatment and reared on control NGM plates for 72 hours. At that time, ten hermaphrodites were transferred to either control NGM plates or 1 µM medip#1 plates prepared as above, and then allowed to acclimate for one hour. After acclimation, individual animals were monitored and the number of pharyngeal pumps occurring in 20 seconds was counted. This was done three times for each individual, waiting at least 20 seconds between counts. The pharyngeal pumping rates for 30 worms were measured for 1 µM medip#1 and control.

**Generation of *mlt-10* Molting Curves.** The molting curve was generated using the protocol described in [7] except that the timing of the experiment was limited to 22 to 36 hours post release from L1 arrest, covering L1 thorough L3 larval stages. The experiment relied on monitoring the level of GFP in GR1395 *mgIs49 [mlt-10::GFP-pest; ttx-1::GFP]* animals [8]. Worms were maintained at 20 °C on OP50 *E. coli* under standard nematode growth conditions[1]. Populations were synchronized by alkaline hypochlorite treatment of gravid hermaphrodites. Isolated eggs were allowed to hatch overnight in M9 buffer with rotation at 20 °C [2]. Two experiments were started 12 hours apart, allowing observations of GFP expression through the transition between the L2 and L3 larval stages through the transition from L3 to L4. In each experiment, a population of GR1395 animals was synchronized by hypochlorite bleaching as described above. Between 30 and 40 L1 larvae from this population were transferred to either control plates or 1 µM medip#1 plates. Each of the two experiments consisted of 3 control and 3 treatment plates, for a total of 427 hermaphrodites (204 controls and 223 1 µM medip#1). Animals were examined every hour and scored for GFP fluorescence on a Leica MZ16F stereomicroscope.

### Chemical Synthesis

**Chemicals and General Synthetic Procedures.** Unless stated otherwise, all chemicals purchased for use were obtained from MilliporeSigma (Burlington, MA). *L*-isoleucine *tert*-butyl ester hydrochloride and *N*,*N'*-dicyclohexylcarbodiimide (DCC) were obtained from TCI (Portland, OR). 1-hydroxybenzotriazole (HOBt) was obtained from Fluka (Mexico City, Mexico). All deuterated solvents were obtained from Cambridge Isotopes (Tewksbury, MA). All non-deuterated solvents were obtained from Fisher Sci (Waltham, MA). Abbreviations used for solvents are as follows: chloroform ($CHCl_3$), dichloromethane (DCM), dimethylsulfoxide (DMSO), and methanol (MeOH).

Unless stated otherwise, all reactions were performed under argon atmosphere in flame-dried glassware. All commercially available reagents were used as purchased unless otherwise stated. All solvents were dried over activated 3Å sieves for a minimum of 24 h unless used in reactions where aqueous reagents were involved. Thin-layer chromatography (TLC) was performed with J.T. Baker Silica Gell IB2-F plastic-backed plates. Reversed-phase column chromatography was performed using Teledyne ISCO CombiFlash Rf and Rf+ systems with Teledyne ISCO RediSep Rf and Rf Gold silica columns. Nuclear Magnetic Resonance (NMR) spectra were recorded on a Varian INOVA 600 (600 MHz) or Bruker AV 500 (500 MHz) in the Cornell University NMR Facility. $^1H$ NMR chemical shifts are reported in ppm (δ) relative to the residual solvent peaks (7.26 ppm for $CDCl_3$, 3.31 ppm for $CD_3OD$, and 2.50 for $D_6$-DMSO) and $^{13}C$ NMR shifts relative to the residual solvent peaks (77.16 for $CDCl_3$ and 49.00 for $CD_3OD$, and 39.52 for $D_6$-DMSO).

***(2O)-* and *(3O)-*acetyl-*S*-methylthioadenosine (acemta#1, 4, and acemta#2, 5)**

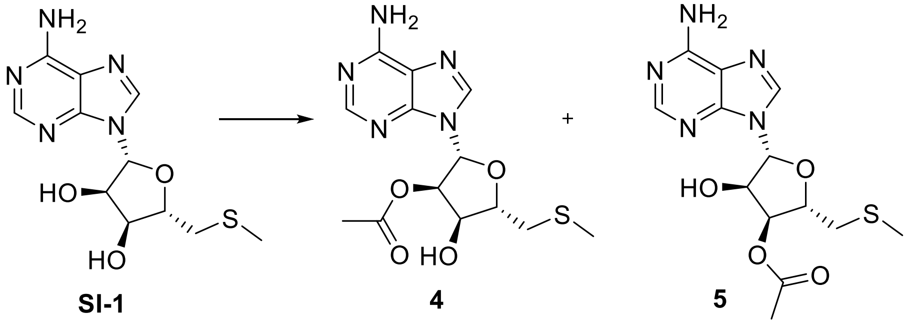


**SI-1** 4 5

*S*-Methylthioadenosine (MTA, **SI-1**, 10 mg, 0.0336 mmol) was dissolved in pyridine (150 µL) and stirred at ambient temperature. Acetic anhydride (34 µL, 0.0336 mmol) was added and the reaction stirred. After 2 h (MeOH, 2 mL) was added and the reaction stirred 30 min. The reaction was diluted with $CHCl_3$ and washed with 2% aqueous acetic acid (1×4 mL) and saturated sodium bicarbonate (1×6 mL) then dried over magnesium sulfate ($MgSO_4$), filtered, and concentrated under reduced pressure. The resulting oil was purified by flash chromatography on silica. Elution with a gradient of 0-40% DCM/MeOH yielded the diacetylated product (10 mg, 74.9%) and a 1:3 mixture of the (2*O*)- and (3*O*)-acetylated products (acemta#1, **4**, and acemta#2, **5**, 2.5 mg, 20.9%) as an oil.

**(2O)-acetyl S-methylthioadenosine (4, acemta#1):**

**$^1$H NMR ($CDCl_3$, 500 MHz):** δ (ppm) 8.31 (s, 1H), 7.98 (s, 1H), 6.09 (d, *J* = 3.4 Hz, 1H), 5.75 (dd, *J* = 5.6, 3.4 Hz, 1H), 4.83 (t, *J* = 6.3 Hz, 1H), 4.25 (q, *J* = 6.0 Hz, 1H), 2.99 (dd, *J* = 14.2, 5.2 Hz, 1H), 1.85-2.95 (m, 1H), 2.17 (s, 3H), 2.15 (s, 3H).

**$^{13}$C NMR ($CDCl_3$, 125 MHz):** δ (ppm) 170.4, 155.5, 153.1, 149.5, 139.7, 120.0, 87.2, 83.2, 76.1, 72.2, 36.4, 20.9, 16.9.

**HRMS (ESI) *m/z*:** Calculated: $(M+H)^+$ 340.1074. Actual: 340.1049. Δ ppm: -7.30.

**(3O)-acetyl S-methylthioadenosine (5, acemta#2):**

**$^1$H NMR ($CDCl_3$, 500 MHz):** δ (ppm) 8.25 (s, 1H), 8.02 (s, 1H), 5.94 (d, *J* = 6.7 Hz, 1H), 5.38 (dd, *J* = 5.8, 2.8 Hz, 1H), 4.97 (t, *J* = 6.3 Hz, 1H), 4.48 (td, *J* = 5.6, 2.8 Hz, 1H), 2.90 (t, *J* = 5.5 Hz, 2H), 2.21 (s, 3H), 2.16 (s, 3H).

**$^{13}$C NMR ($CDCl_3$, 125 MHz):** δ (ppm) 170.6, 155.6, 152.7 (2C), 139.3, 89.5, 83.5, 75.0, 73.8, 37.1, 21.1, 17.0.

**N,N-Dimethyltryptophan (24)**

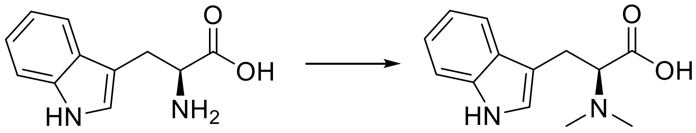


**SI-2** **24**

Based on a synthesis for *N,N*-dimethyltryptamine[9], tryptophan (**SI-2**, 1.0 g, 4.9 mmol) was stirred in MeOH (30 mL) at 0 °C under an atmosphere of argon. Sodium cyanoborohydride (1.5 g, 24 mmol) was added followed by formaldehyde solution (1.54 mL, 36%, 20 mmol) and the reaction was stirred for 36 h. The reaction was concentrated and recrystallized from MeOH to yield **24** (636 mg, 56%) as white crystals that were used without further purification.

**$^1$H NMR ($D_6$-DMSO, 500 MHz):** δ (ppm) 10.83 (s, 1H), 7.54 (d, *J* = 7.8 Hz, 1H), 7.32 (d, *J* = 7.8 Hz, 1H), 7.06 (td, *J* = 7.2, 1.0 Hz, 1H), 6.97 (td, *J* = 7.2, 1.0 Hz, 1H), 3.47 (dd, *J* = 8.0, 6.1 Hz, 1H), 3.34 (dd, *J* = 14.8, 7.9 Hz, 1H), 2.97 (dd, *J* = 14.7, 6.2 Hz), 2.41 (s, 6H).

**$^{13}$C NMR ($D_6$-DMSO, 125 MHz):** δ (ppm) 171.0, 136.1, 127.2, 123.4, 120.9, 118.3, 118.2, 111.3, 110.5, 68.4, 41.3, 24.3.

**HRMS (ESI) *m/z*:** Calculated: $(M+H)^+$ 233.1289. Actual: 233.1270. Δ ppm: 4.27.

**medip#1-*tert*-butyl ester (SI-3)**

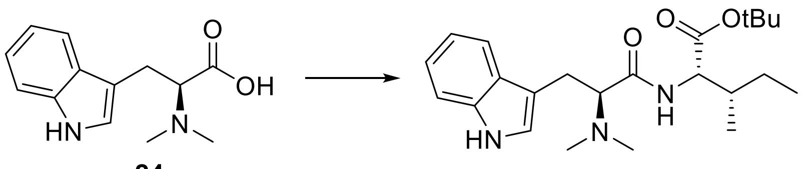


**24** **SI-3**

HOBt (11 mg, 0.0647 mmol), triethylamine (15 µL, 0.108 mmol), DCC (11.6 mg, 0.056 mmol), and isoleucine-*tert*-butyl ester (20 mg, 0.086 mmol) were added to a solution of *N,N*-dimethyltryptophan (**24**, 10 mg, 0.043 mmol) in DCM (1 mL) and DMF (1 mL) and stirred overnight. The reaction was quenched with water (1 mL), extracted with DCM (3×5 mL), dried over $MgSO_4$, filtered, and concentrated under reduced pressure. The resulting solid was purified via flash chromatography on silica gel. Elution with a gradient of 0-10% DCM/MeOH yielded a mixture of medip#1-*tert*-butyl ester (**SI-3**) and isoleucine-*tert*-butyl ester (27.8 mg), which was used without further purification.

**medip#1 (26)**

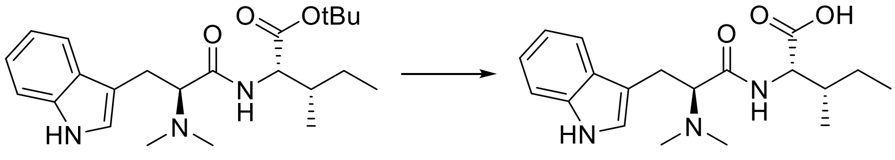


**SI-3** **26**

TFA (530 µL, 6.88 mmol) was added to a stirring solution of medip#1-*tert*-butyl ester (**SI-3**) and isoleucine-*tert*-butyl ester (27.8 mg) in DCM (530 µL) and stirred for 4 h. The reaction was then concentrated under reduced pressure. The resulting oil was purified via flash chromatography on silica gel. Elution with a gradient of 0-20% DCM/MeOH yielded medip#1 (**26**) as a white solid (14.9 mg, ~100%).

**$^1$H NMR ($CD_3OD$, 500 MHz):** δ (ppm) 7.59 (d, *J* = 8.0 Hz, 1H), 7.36 (d, *J* = 8.1 Hz, 1H), 7.22 (s, 1H), 7.12 (td, *J* = 7.3, 1 Hz, 1H), 7.05 (td, *J* = 7.3, 1 Hz, 1H), 4.29 (d, *J* = 5.5 Hz, 1H), 4.22 (dd, *J* = 8.9, 5.3 Hz, 1H), 3.50 (dd, *J* = 14.6, 5.2 Hz, 1H), 3.42 (dd, *J* = 14.6, 8.9 Hz, 1H), 2.98 (s, 6H), 1.78-1.89 (m, 1H), 1.35-1.48 (m, 1H), 1.08-1.18 (m, 1H), 0.89 (t, *J* = 7.5 Hz, 3H), 0.88 (d, *J* = 6.8 Hz, 3H).

**$^{13}$C NMR ($CD_3OD$, 125 MHz):** δ (ppm) 173.2, 168.5, 138.1, 128.2, 125.6, 120.3, 118.8, 112.6, 107.2, 69.5, 58.4, 54.8, 42.4, 38.4, 26.2, 25.7, 15.9, 11.7.

**HRMS (ESI) *m/z*:** Calculated: $(M+H)^+$ 346.2125. Actual: 346.2102. Δ ppm: 6.77.

### Supplementary Figures

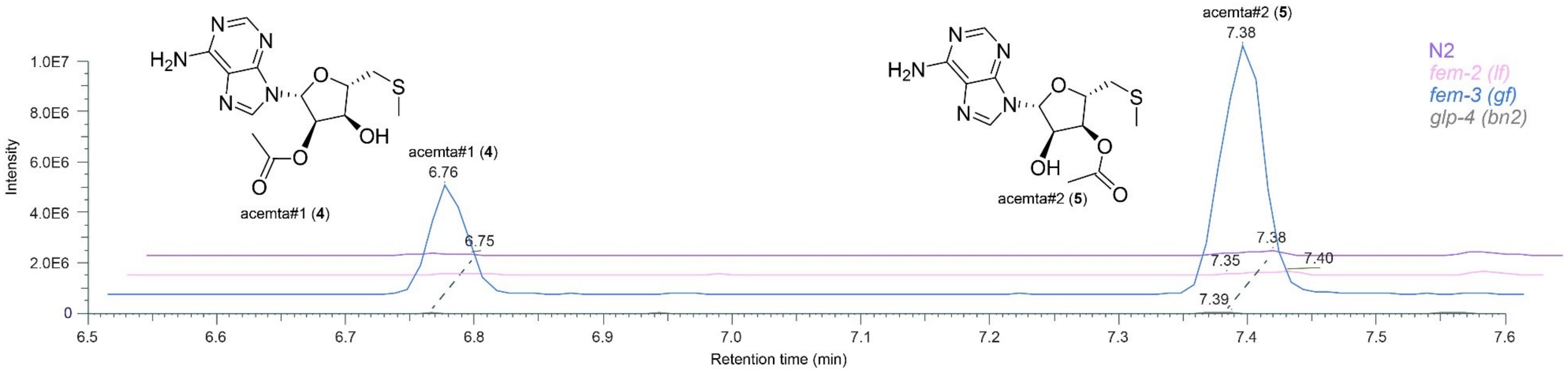


**Supplementary Figure 1. Extracted ion chromatograms (EICs) of acemta#1 (4) and acemta#2 (5).** EIC (ESI+) of *m/z* 340.1074 in N2 and *him-5* endo-metabolome samples showing peaks for acemta#1 (6.76 min) and acemta#2 (7.38 min) and MS2 spectrum for acemta#2 acquired from a *fem-3 (gf)* endo-metabolome sample.

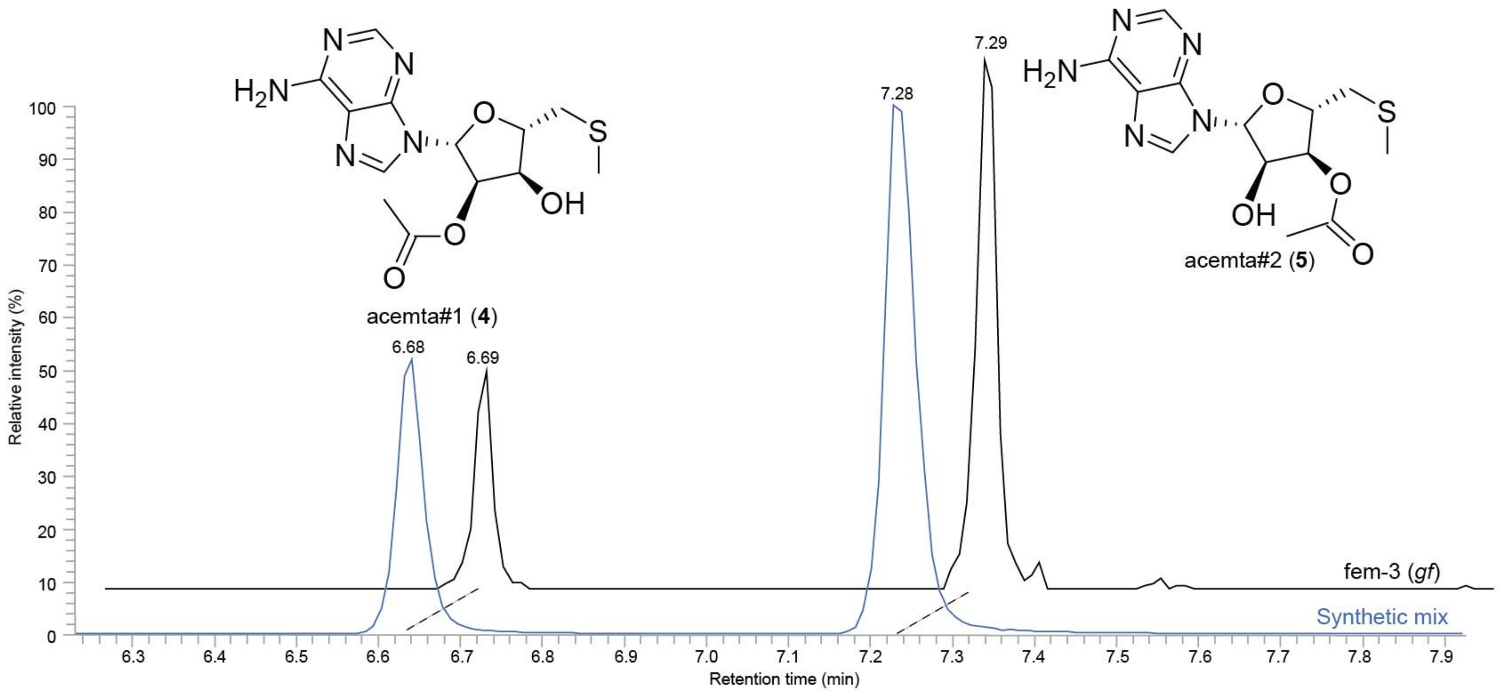


**Supplementary Figure 2. EICs of natural and synthetic acemta#1 (4) and acemta#2 (5).** EIC (ESI+) of *m/z* 340.1074 in a *fem-3 (gf)* endo-metabolome sample (black) and a synthetic standard containing a mixture of acemta#1 and acemta#2 (blue).

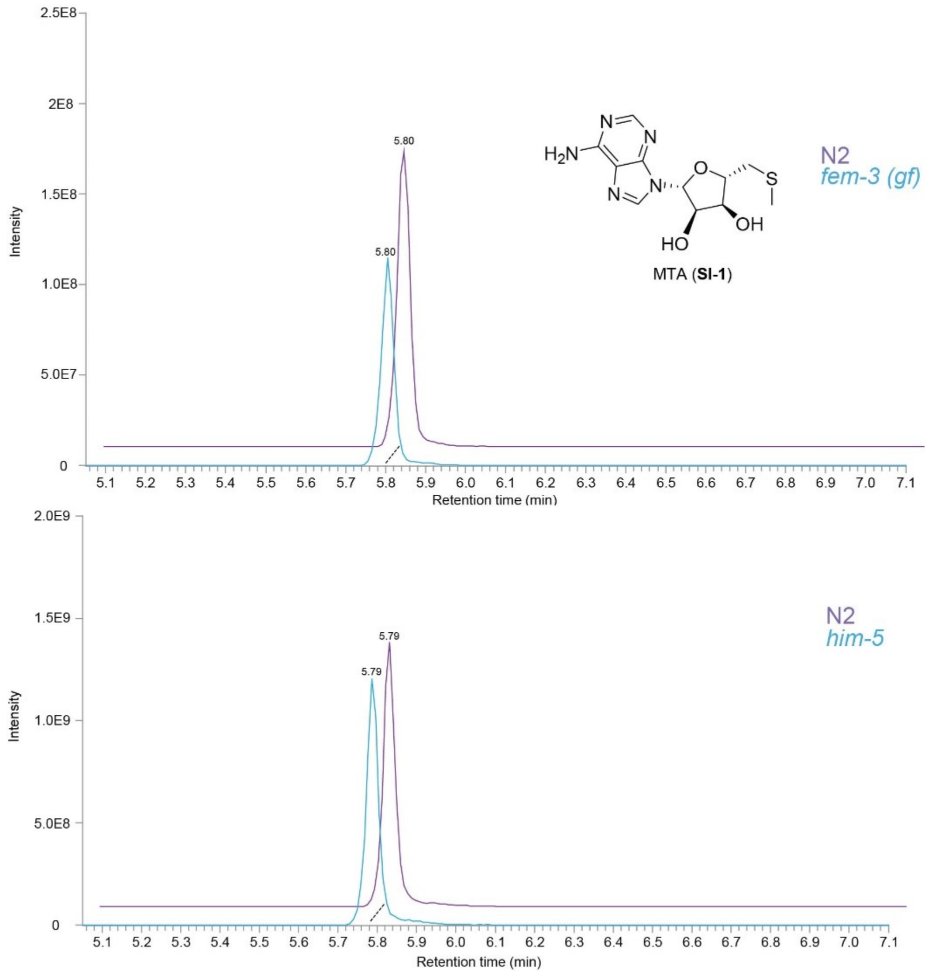


**Supplementary Figure 3.** Comparison of signal intensity of putative precursor to acemta derivatives (**4** and **5**), methylthioadenosine (MTA, **SI-1**) for WT, *fem-3 (gf)*, and *him-5* endo-metabolome samples reveals very little differences. Note: the comparisons were done via biological replicates of WT *C. elegans*, notable by the differences in signal intensity between samples.

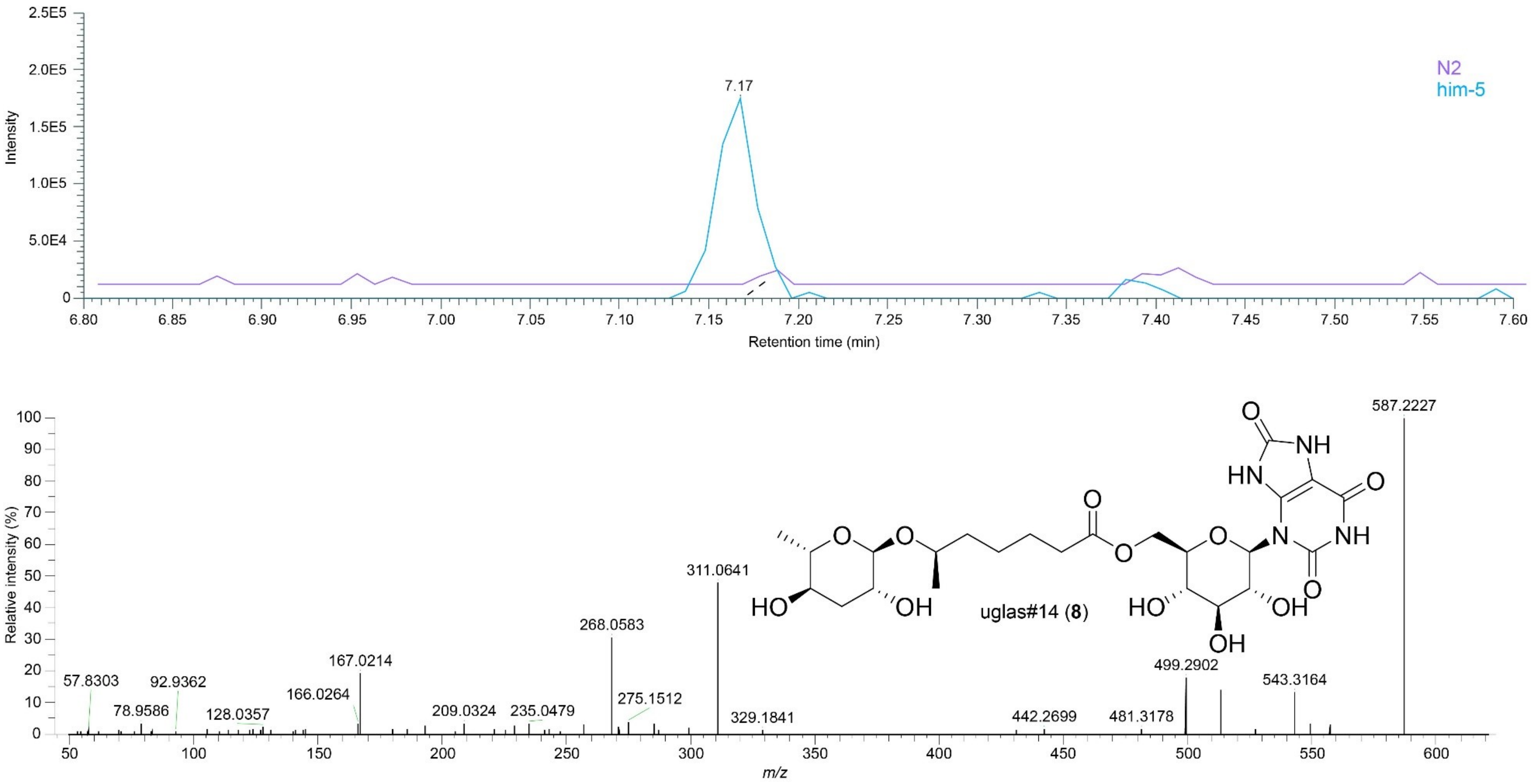


**Supplementary Figure 4. EICs and MS2 spectrum of uglas#14 (8).** EIC (ESI-) of *m/z* 587.2206 in WT and *him-5* endo-metabolome samples showing peaks for uglas#14 and MS2 spectrum (ESI-) for uglas#14 acquired from a *him-5* endo-metabolome sample.

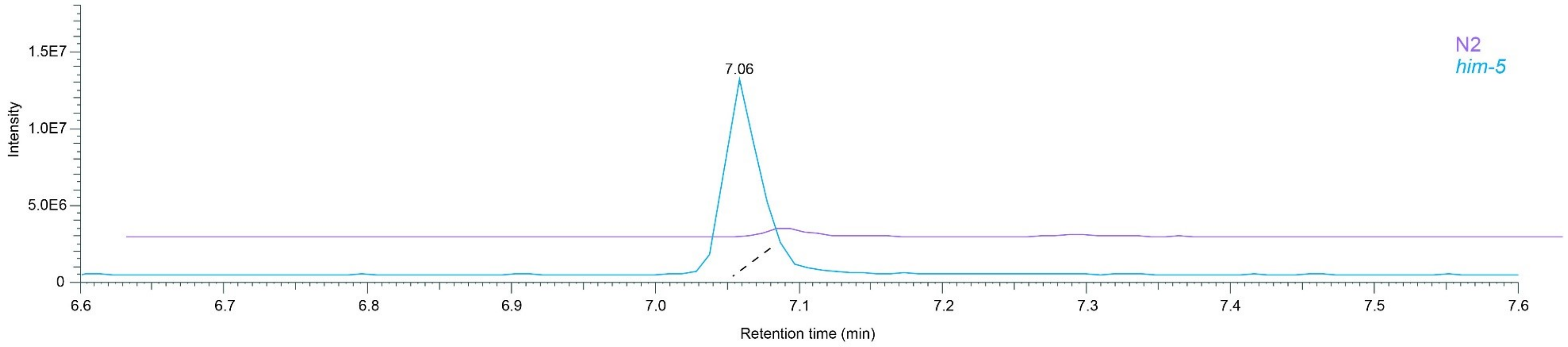


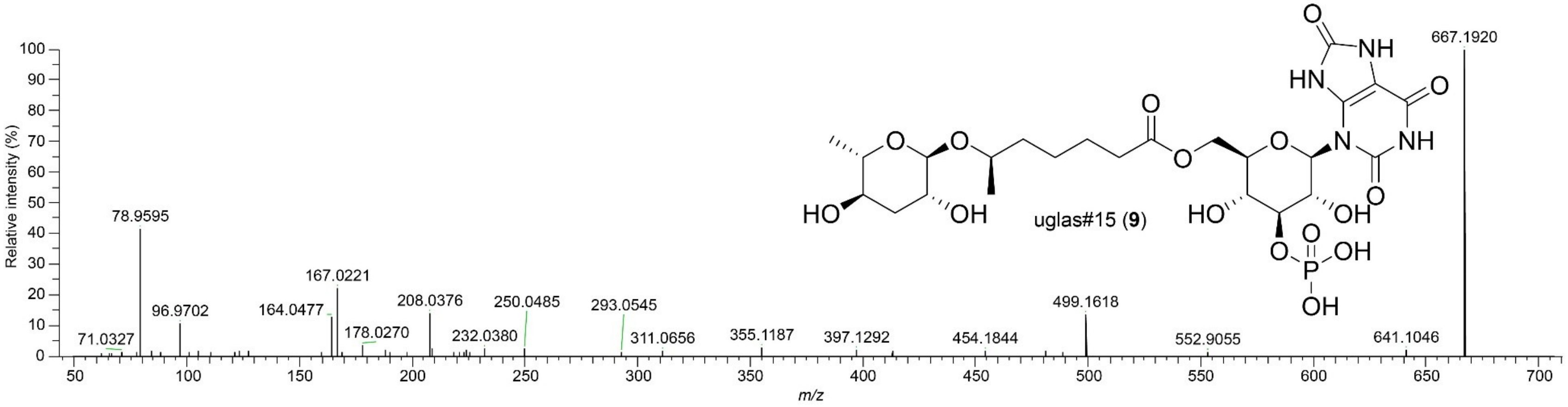


**Supplementary Figure 5. EICs and MS2 spectrum of uglas#15 (9).** EIC (ESI-) of *m/z* 667.1869 in WT and *him-5* endo-metabolome samples showing peaks for uglas#15 and MS2 spectrum (ESI-) for uglas#15 acquired from a *him-5* endo-metabolome sample.

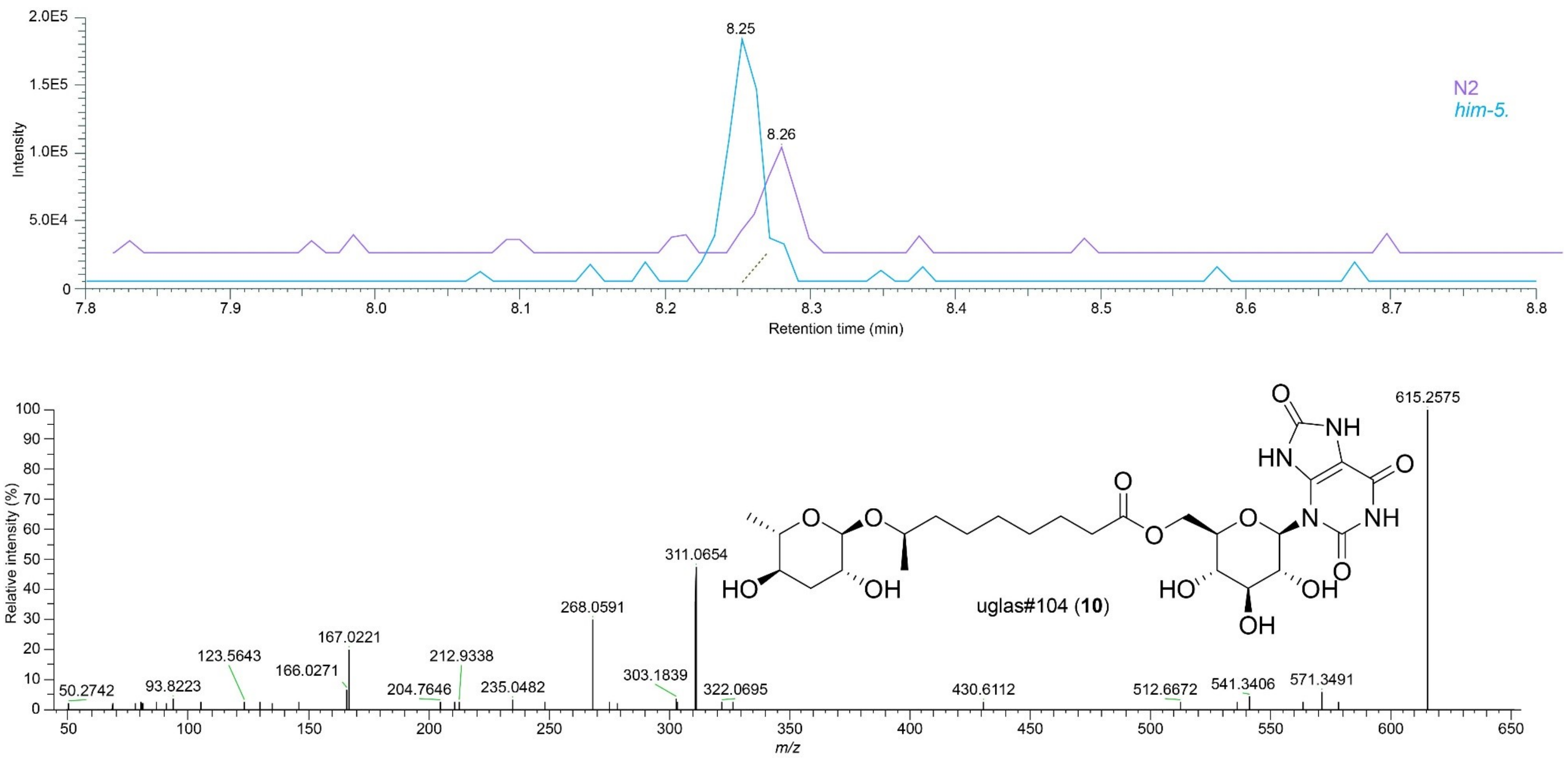


**Supplementary Figure 6. EICs and MS2 spectrum of uglas#104 (10).** EIC (ESI-) of *m/z* 615.2519 in WT and *him-5* endo-metabolome samples showing peaks for uglas#104 and MS2 spectrum (ESI-) for uglas#104 acquired from a *him-5* endo-metabolome sample.

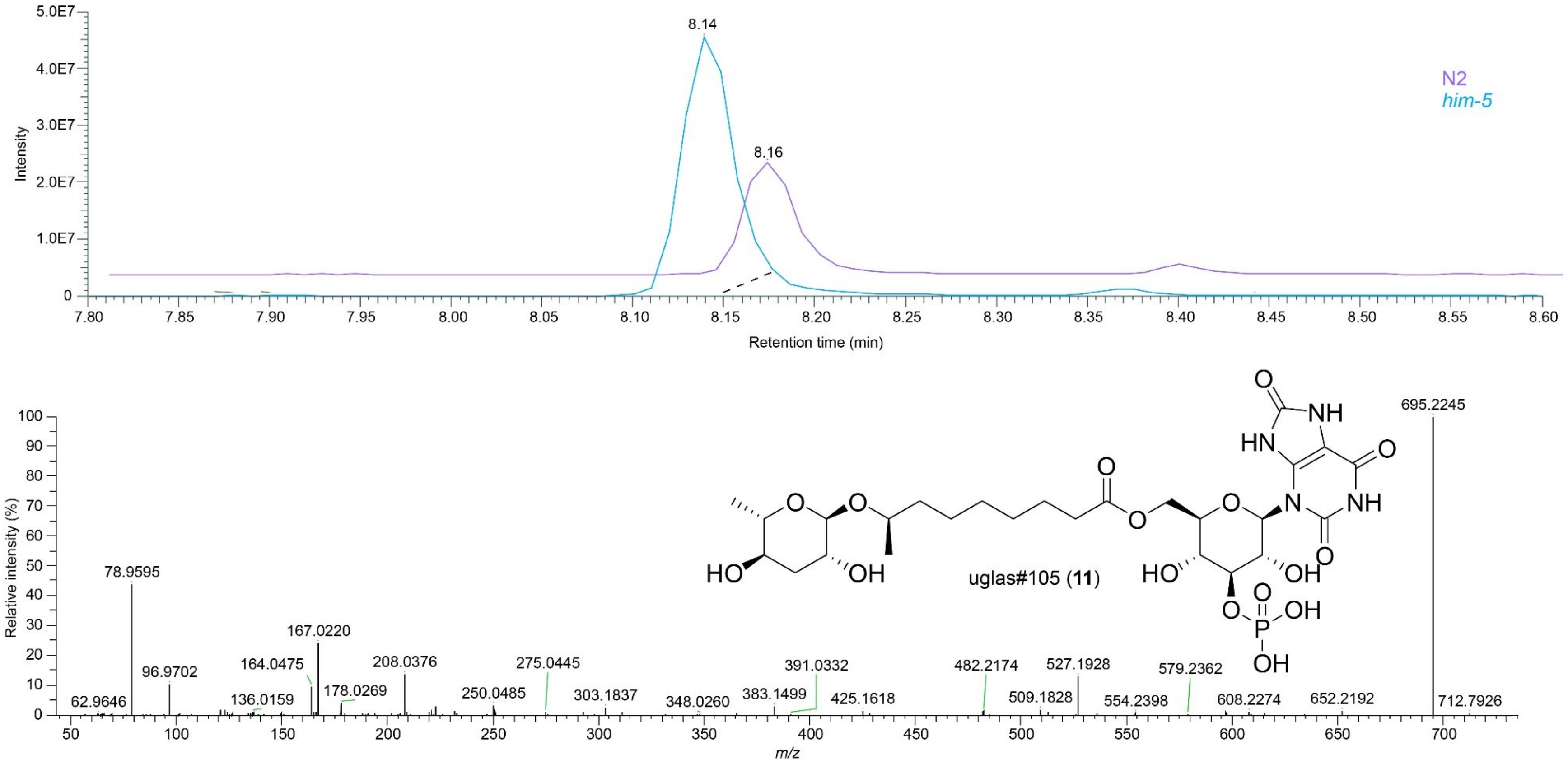


**Supplementary Figure 7. EICs and MS2 spectrum of uglas#105 (11).** EIC (ESI-) of *m/z* 695.2182 in WT and *him-5* endo-metabolome samples showing peaks for uglas#105 and MS2 spectrum (ESI-) for uglas#105 acquired from a *him-5* endo-metabolome sample.

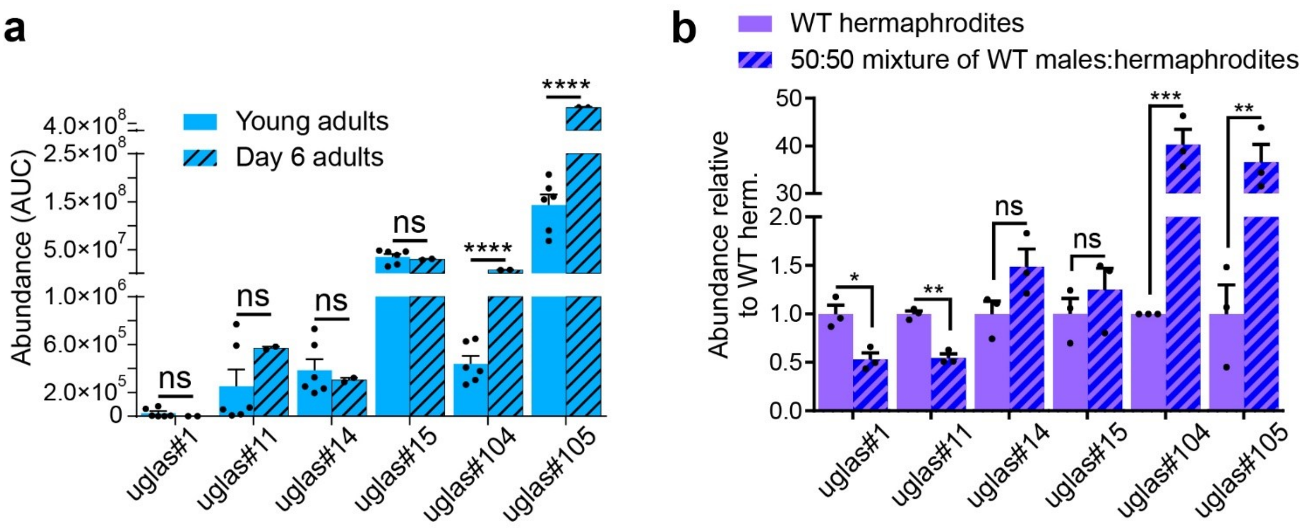


**Supplementary Figure 8.** Relative abundances of uglas-family metabolites in the *endo*-metabolomes of (a) young adults compared to day-6 adults, and (b) small plate-based samples of WT hermaphrodites and 50:50 mixtures of males and hermaphrodites.

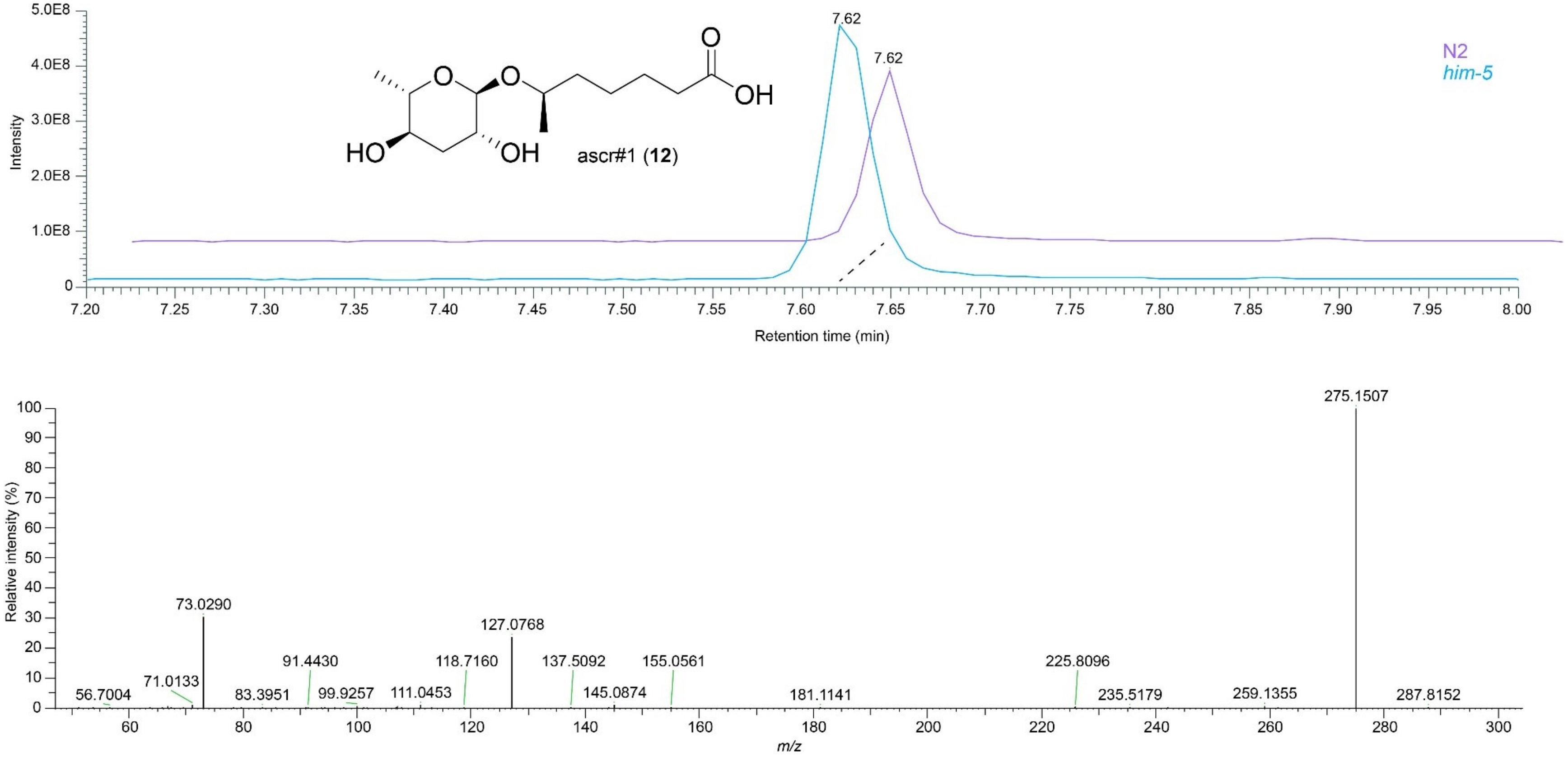


**Supplementary Figure 9. EICs and MS2 spectrum of ascr#1 (12).** EIC (ESI-) of *m/z* 275.1500 in WT and *him-5* exo-metabolome samples showing peaks for ascr#1 and MS2 spectrum (ESI-) for ascr#1 acquired from a *him-5* exo-metabolome sample.

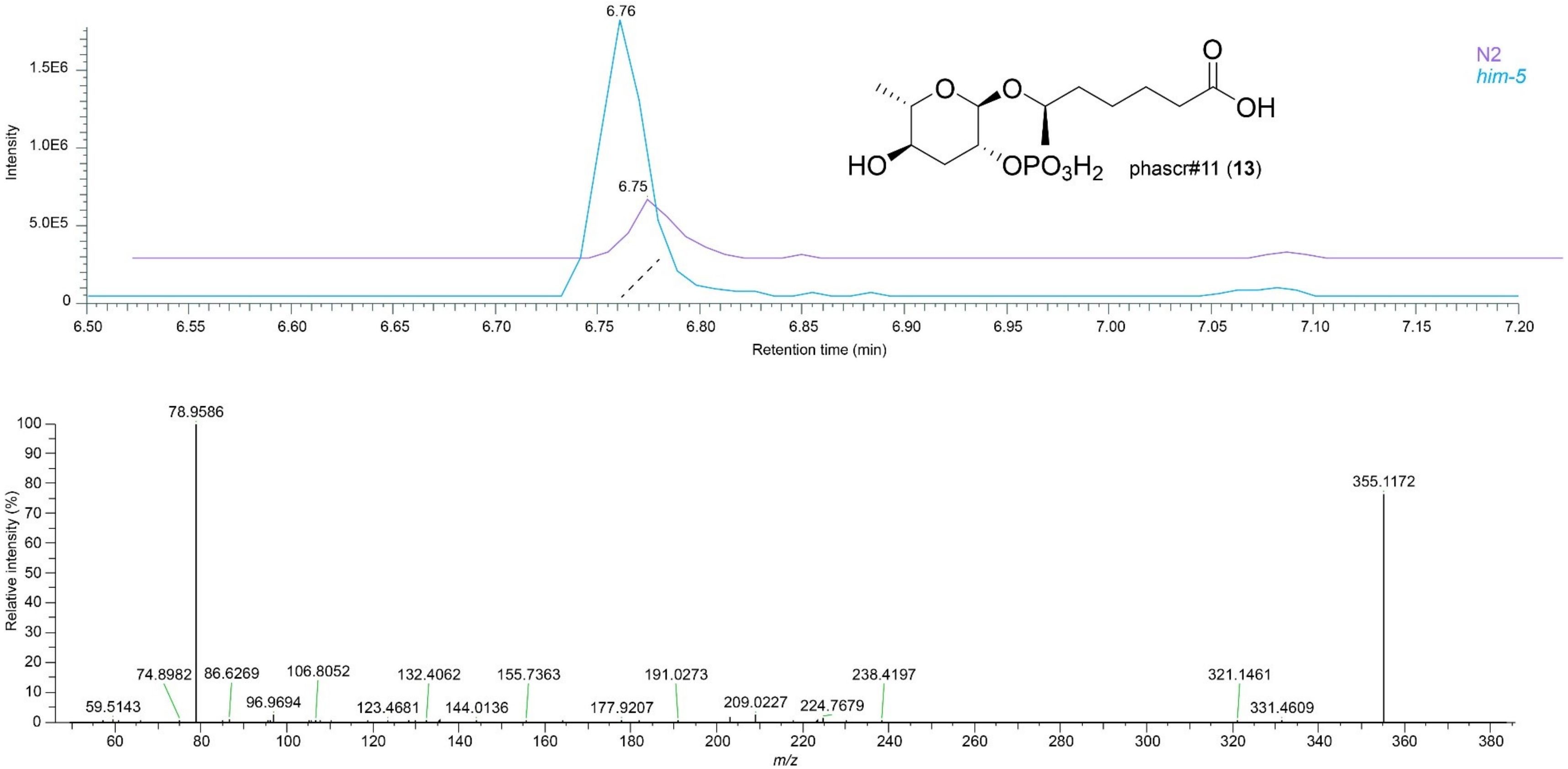


**Supplementary Figure 10. EICs and MS2 spectrum of phascr#11 (13).** EIC (ESI-) of *m/z* 355.1168 in WT and *him-5* exo-metabolome samples showing peaks for phascr#11 and MS2 spectrum (ESI-) for phascr#11 acquired from a *him-5* exo-metabolome sample[10].

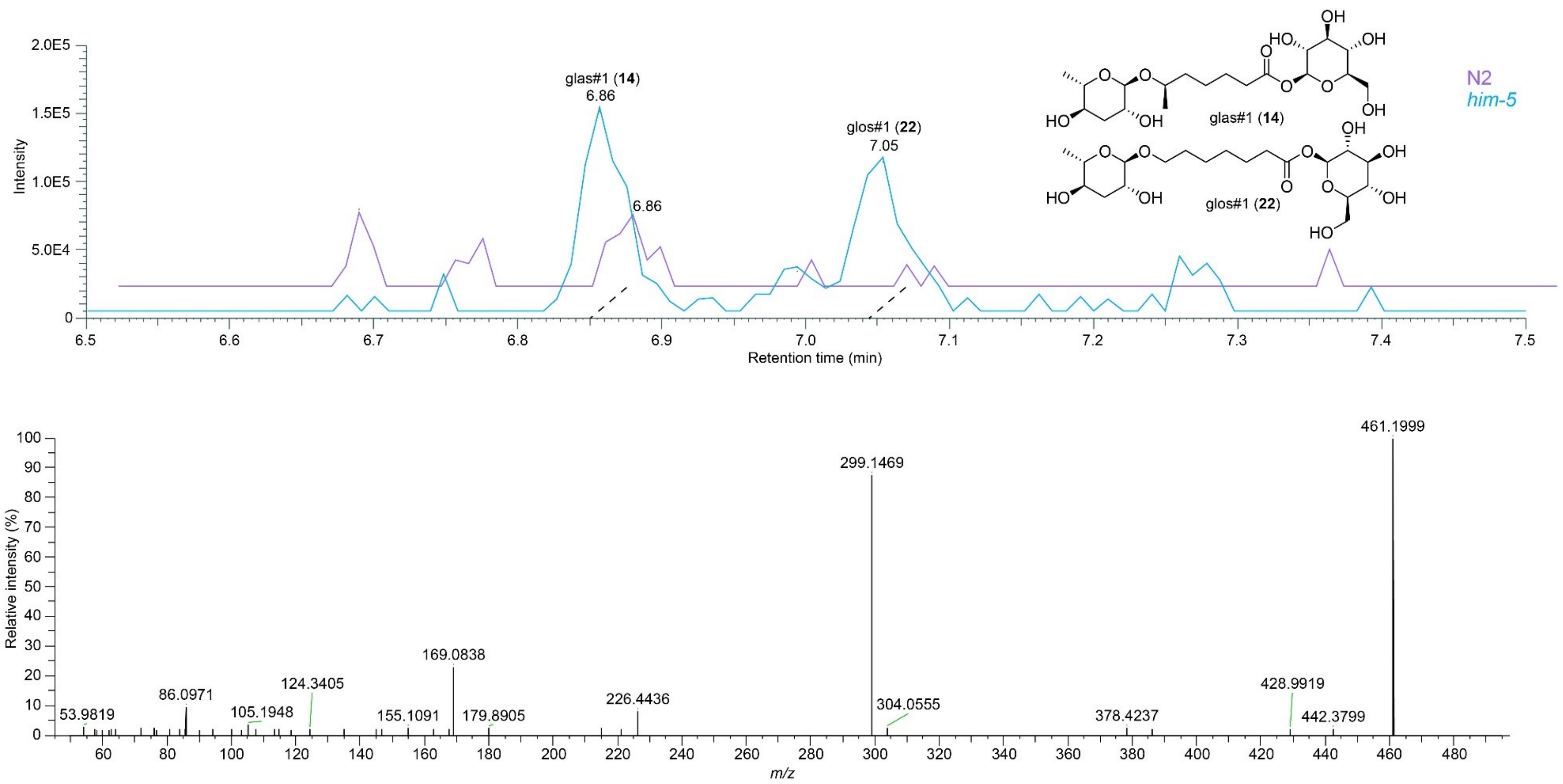


**Supplementary Figure 11. EICs and MS2 spectrum of glas#1 (14) and glos#1 (22).** EIC (ESI+) of *m/z* 461.1995 in WT and *him-5* endo-metabolome samples showing peaks for glas#1 and glos#1, and MS2 spectrum (ESI+) for glas#1 acquired from a *him-5* endo-metabolome sample.

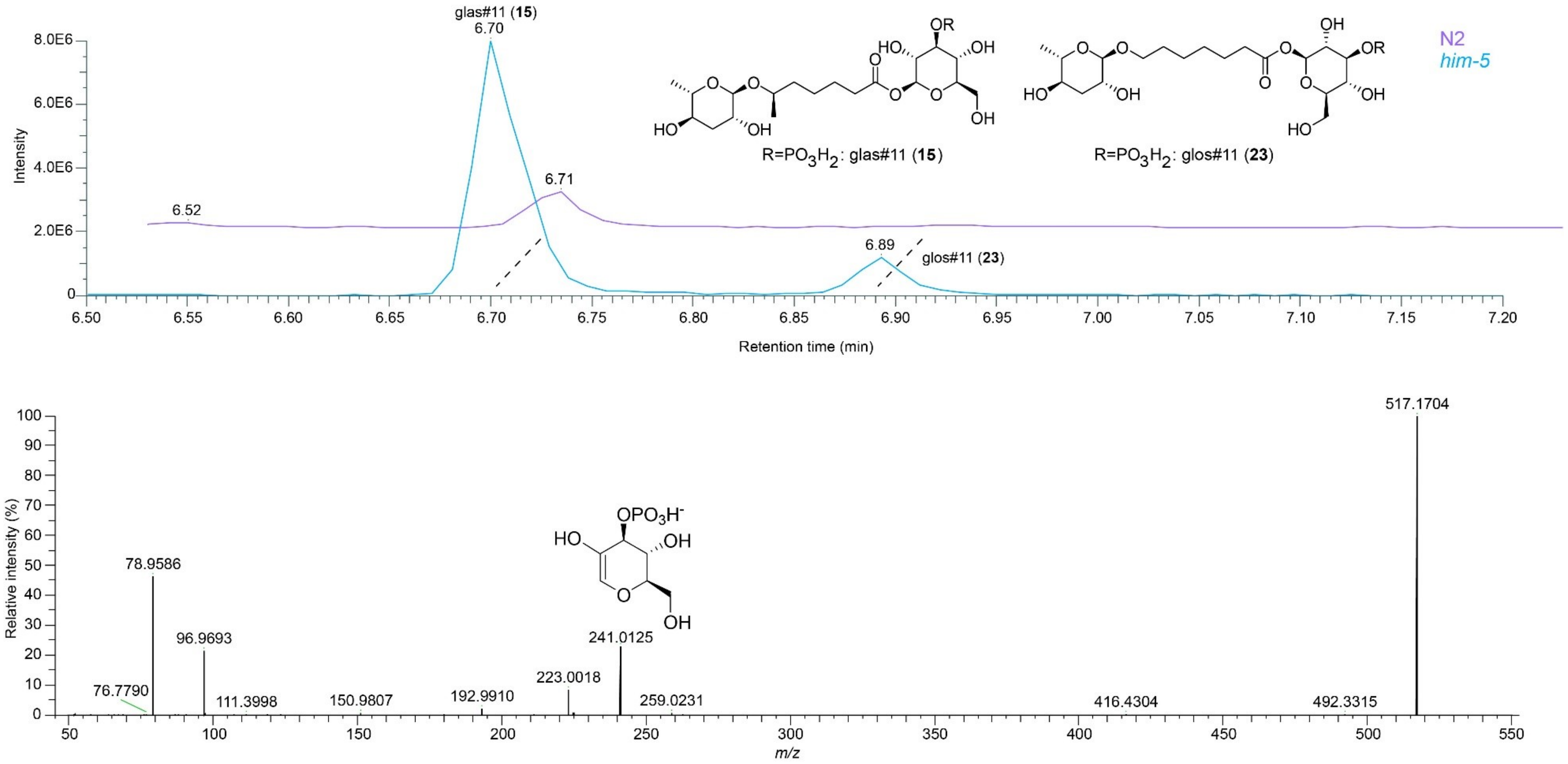


**Supplementary Figure 12. EICs and MS2 spectrum of glas#11 (15) and glos#11 (23).** EIC (ESI-) of *m/z* 517.1669 in WT and *him-5* exo-metabolome samples showing peaks for glas#11 and glos#11 and MS2 spectrum (ESI-) for glas#11 acquired from a *him-5* exo-metabolome sample. A key fragment suggesting phosphorylation on the glucose moiety is shown for *m/z* = 241.0125.

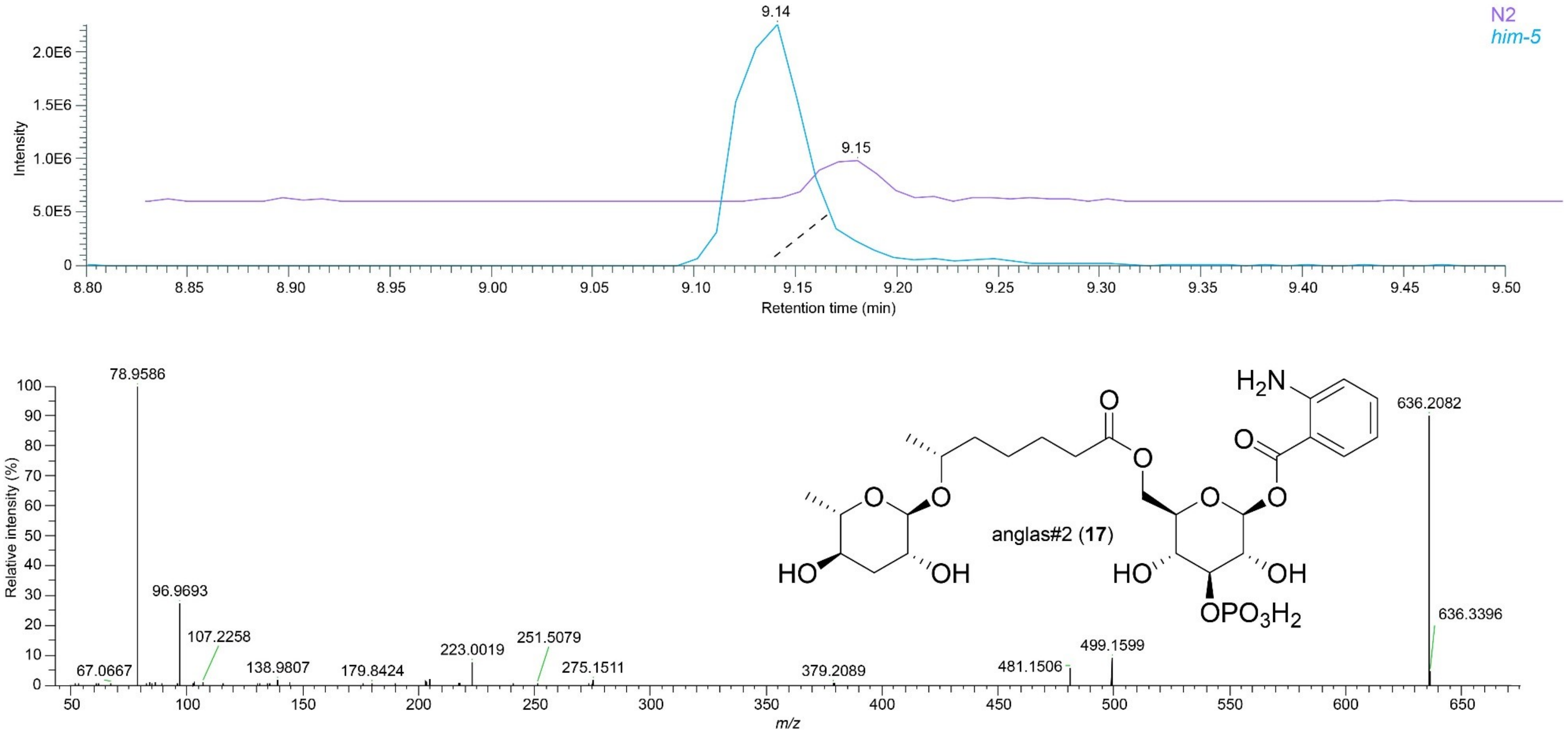


**Supplementary Figure 13. EICs and MS2 spectrum of anglas#2 (17).** EIC (ESI-) of *m/z* 636.2070 in WT and *him-5* endo-metabolome samples showing peaks for anglas#2 and MS2 spectrum (ESI-) for anglas#2 acquired from a *him-5* endo-metabolome sample.

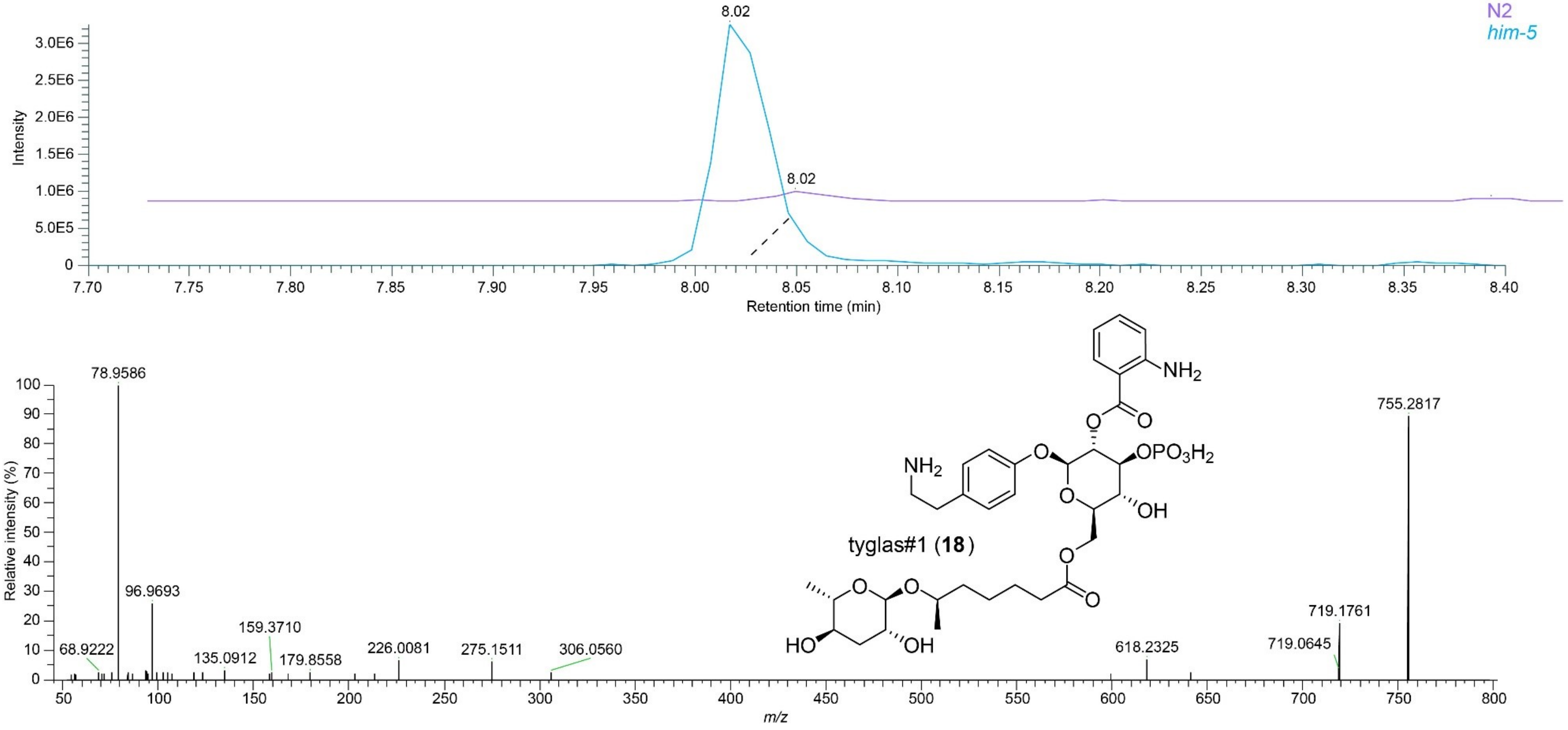


**Supplementary Figure 14. EICs and MS2 spectrum of tyglas#1 (18).** EIC (ESI-) of *m/z* 755.2807 in WT and *him-5* endo-metabolome samples showing peaks for tyglas#1 and MS2 spectrum (ESI-) for tyglas#1 acquired from a *him-5* endo-metabolome sample See reference for ESI+ MS2 spectrum[11].

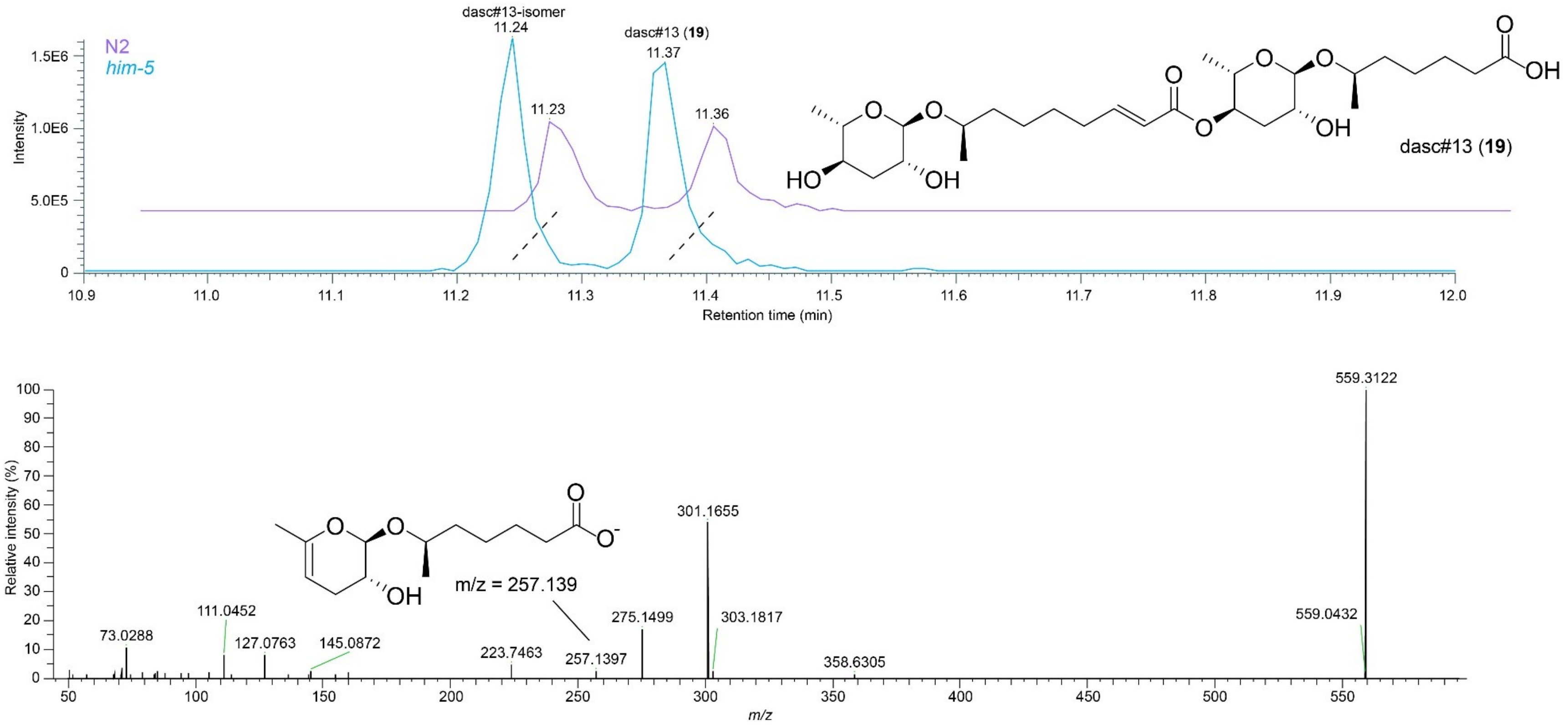


**Supplementary Figure 15. EICs and MS2 spectrum of dasc#13 (19).** EIC (ESI-) of *m/z* 559.3129 in WT and *him-5* exo-metabolome samples showing peaks for dasc#13 (11.37 min) and MS2 spectrum (ESI-) for dasc#13 (**19**) acquired from a *him-5* exo-metabolome sample. The earlier eluting isomer (11.2 min) is a dasc#13 structural isomer via MS2 analysis. Key fragment (*m/z* = 257.1397) is displayed, suggesting the loss of ascr#3 via elimination at either the 2' or 4'-position (displayed as 4').

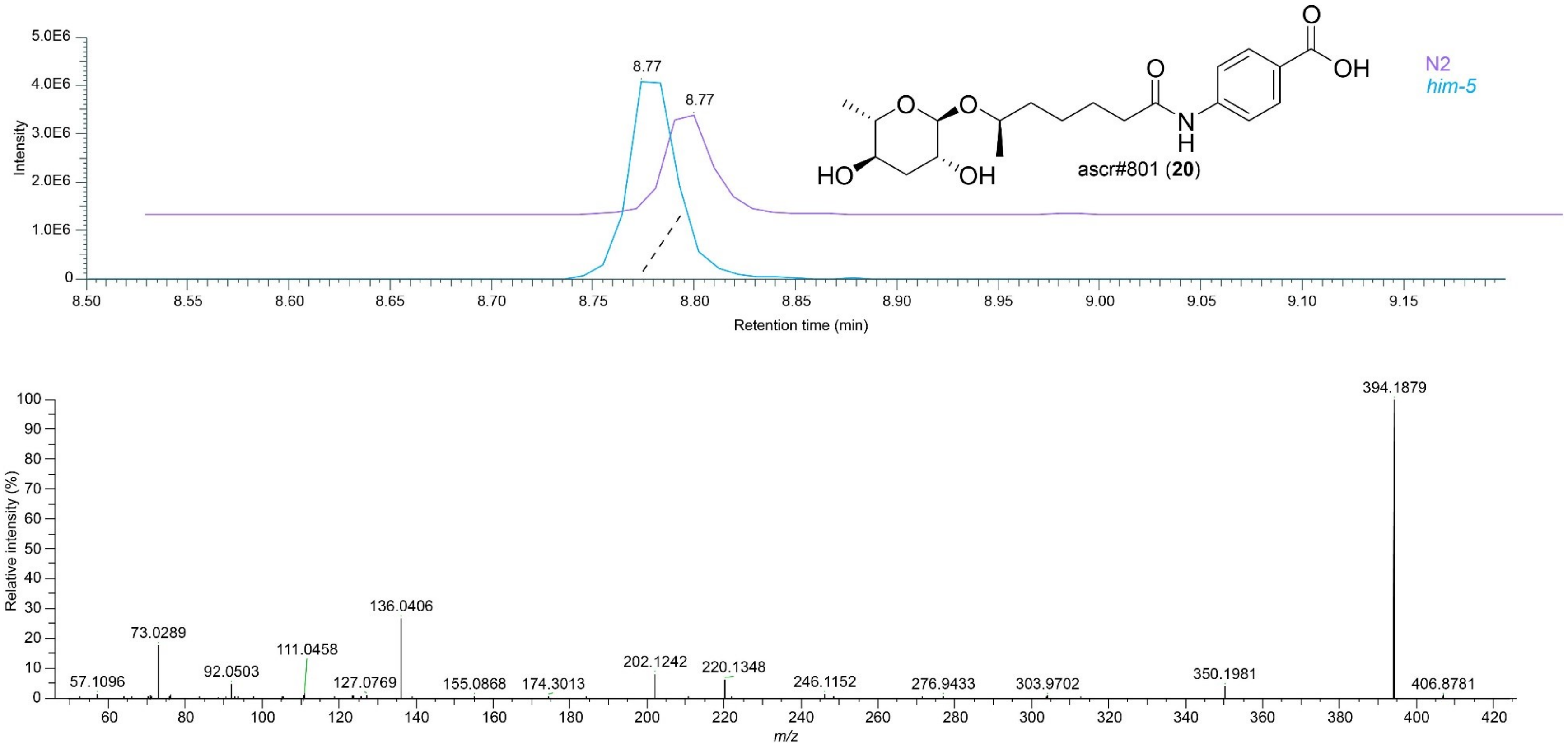


**Supplementary Figure 16. EICs and MS2 spectrum of ascr#801 (20).** EIC (ESI-) of *m/z* 394.1875 in WT and *him-5* exo-metabolome samples showing peaks for ascr#801 and MS2 spectrum (ESI-) for ascr#801 acquired from a *him-5* exo-metabolome sample.

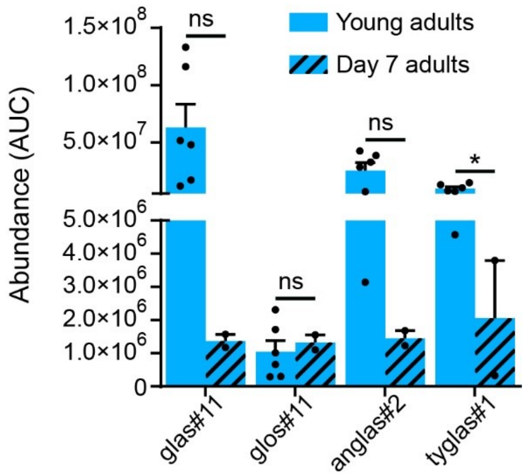


**Supplementary Figure 17.** Abundances of *him-5*-enriched ascr#1 derivatives are higher in *endo*-metabolome samples from young adults than from day-7 adults. Data are presented as mean ± s.e.m. *, p≤0.05; ns, not statistically significant.

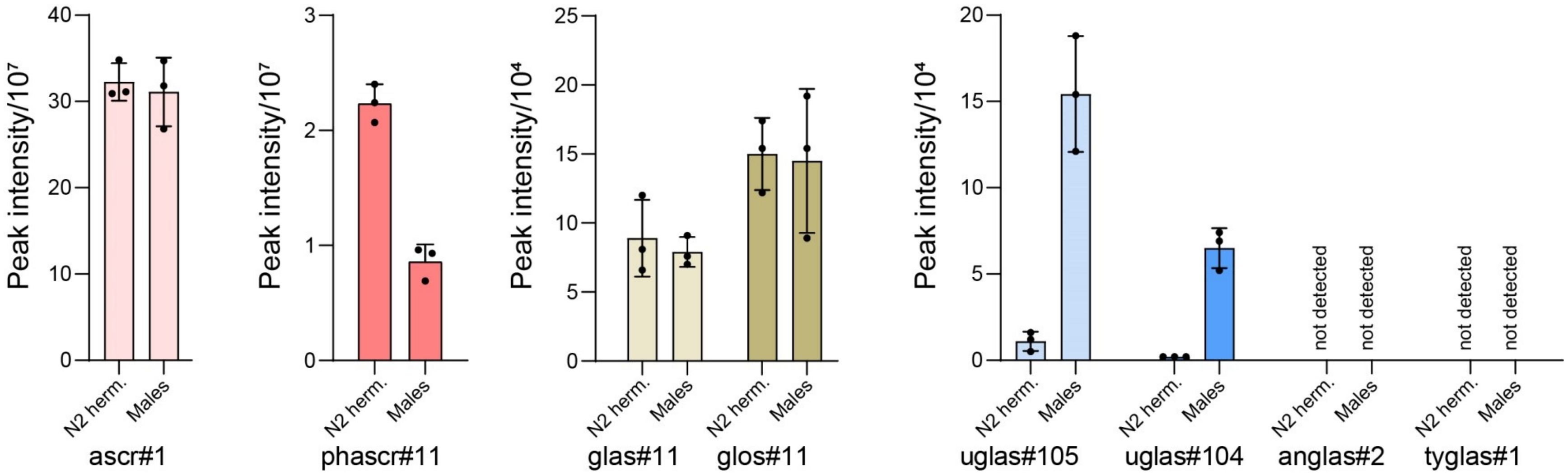


**Supplementary Figure 18.** Abundances of ascr#1 and ascr#1 derivatives enriched in *him-5 endo*-metabolomes are not enriched (ascr#1, phascr#11, glas#11, glos#11) or not detected (anglas#2, tyglas#1) in *endo*-metabolome samples of hand-picked, pure males compared to pure hermaphrodites, whereas uglas#104 and uglas#105 are consistently enriched in males.

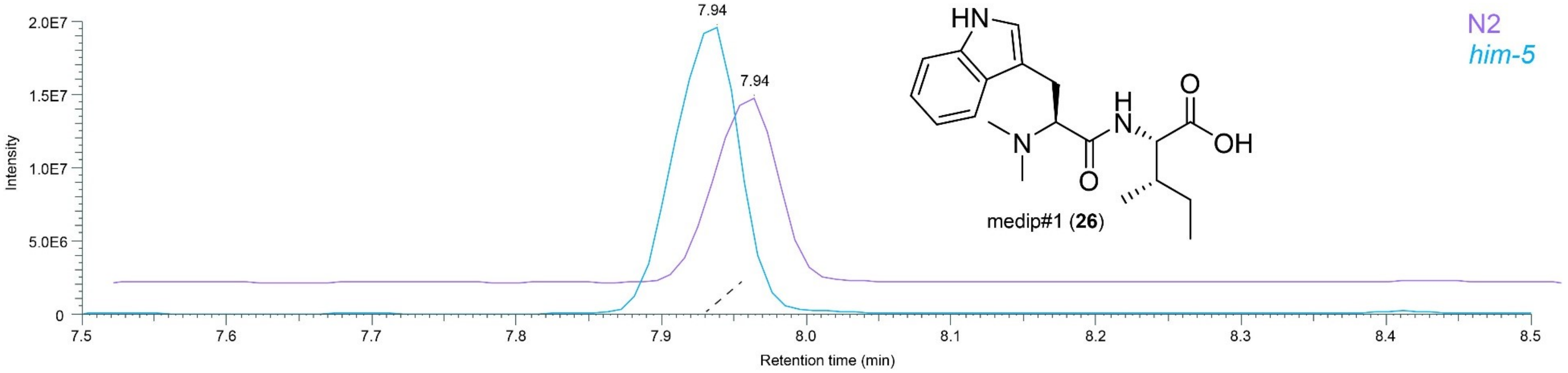


**Supplementary Figure 19.** EICs (ESI+) of *m/z* 346.2125 in WT and *him-5* exo-metabolome samples showing peaks for medip#1.

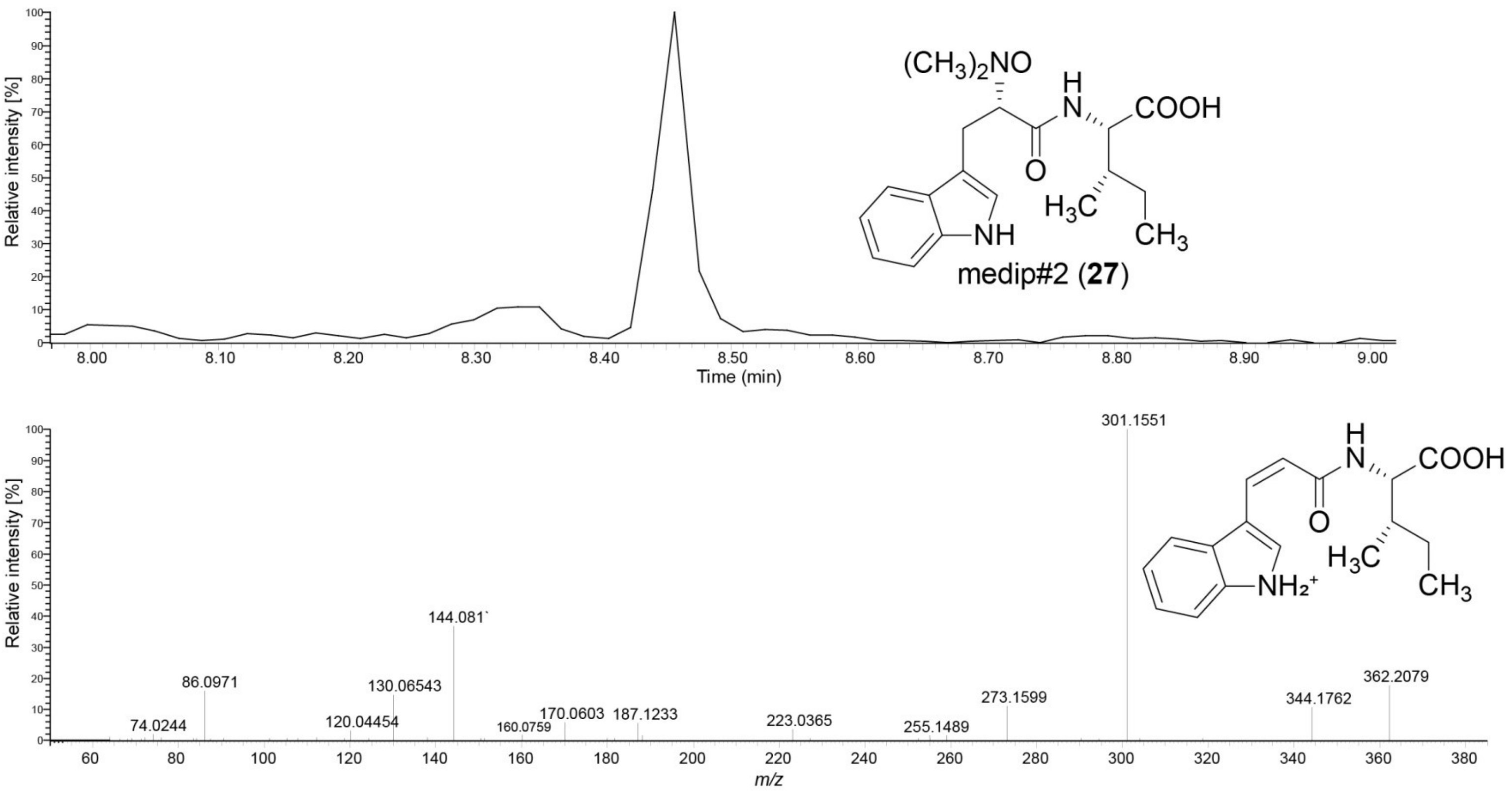


**Supplementary Figure 20. EIC and MS2 spectrum of medip#2 (27).** EIC (ESI+) of *m/z* 362.2074 in WT and *him-5* exo-metabolome samples showing a peak for medip#2 (top), and MS2 spectrum (ESI+) for medip#2 acquired from a *him-5* exo-metabolome sample showing key fragment at *m/z* 301.1551.

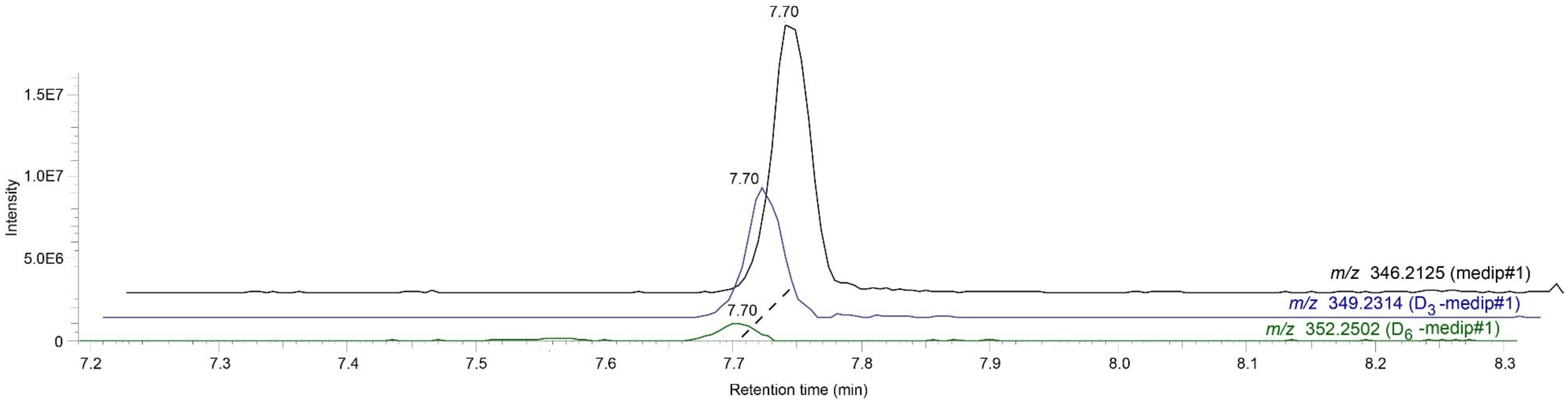


**Supplementary Figure 21. $D_3$-Methionine labeling of medip#1.** EICs (ESI+) for medip#1 (black), $D_3$-medip#1 (blue), $D_6$-medip#1 (green) in the exo-metabolome of *him-5* animals fed $D_3$-methionine.

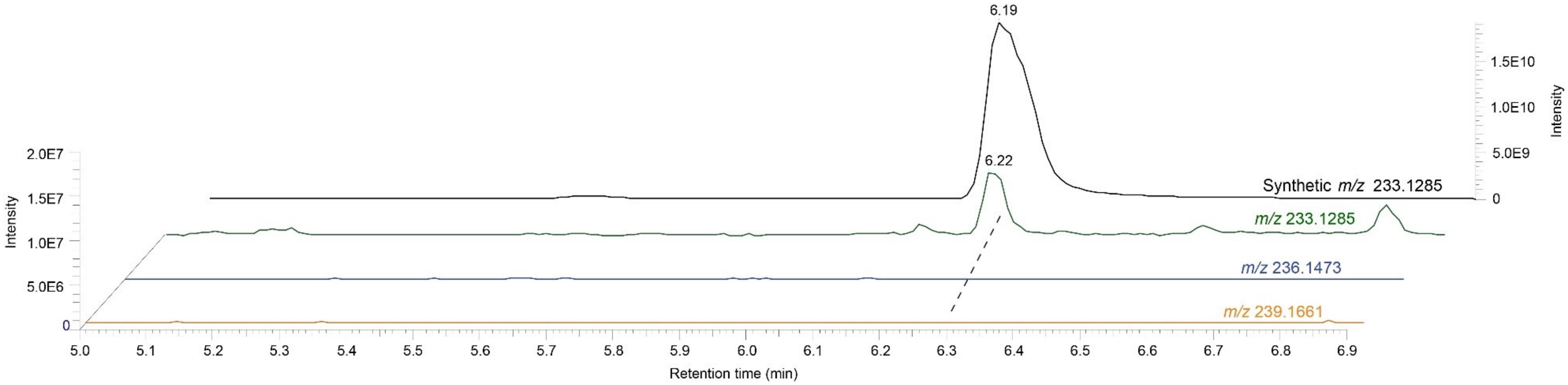


**Supplementary Figure 22. $D_3$-Methionine feeding does not label dimethyltryptophan.** EICs (ESI+) of synthetic dimethyltryptophan (black), unlabeled dimethyltryptophan in the *him-5* exo-metabolome (green), the trace for the *m/z* of $D_3$-dimethyltryptophan, lacking a co-eluting peak (blue), and the trace for the *m/z* of $D_6$- dimethyltryptophan (orange), also lacking a co-eluting peak.

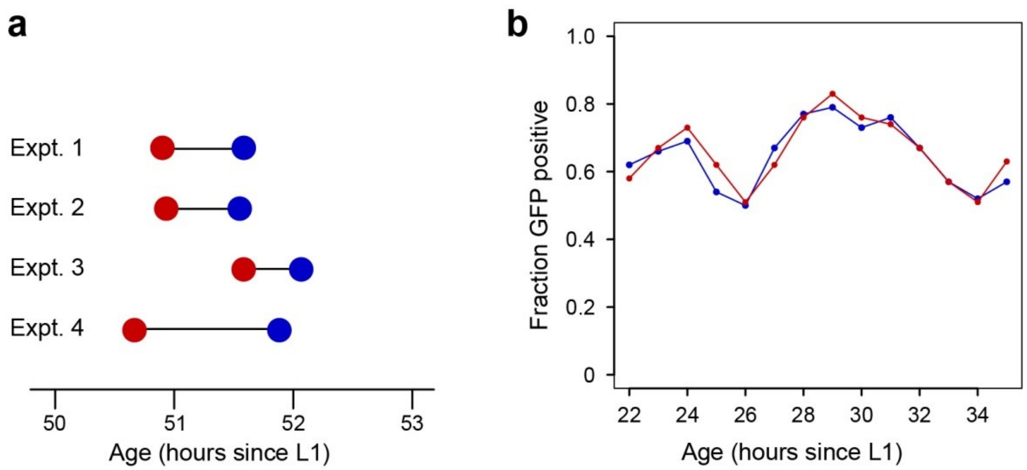


**Supplementary Figure 23. Developmental assays with medip#1. a.** Time to sexual maturation of hermaphrodites raised on 1 µM medip#1-conditioned plates (red) *vs.* paired control (blue). Median time of morphologically-defined adulthood is shown for four independent experiments. **b.** L3 larvae do not show acceleration of development. Fractions of GFP-positive *mlt-10*::GFP hermaphrodites raised on medip#1 (red) or control (blue). Two experiments were started twelve hours apart to allow the observation of animals from the L2 to L3 transition (roughly 26 h after release from L1 arrest) through the beginning of the transition from L3 to L4. At none of the points was there a significant difference between the two curves. (n = 204 control animals and 223 medip#1 animals).

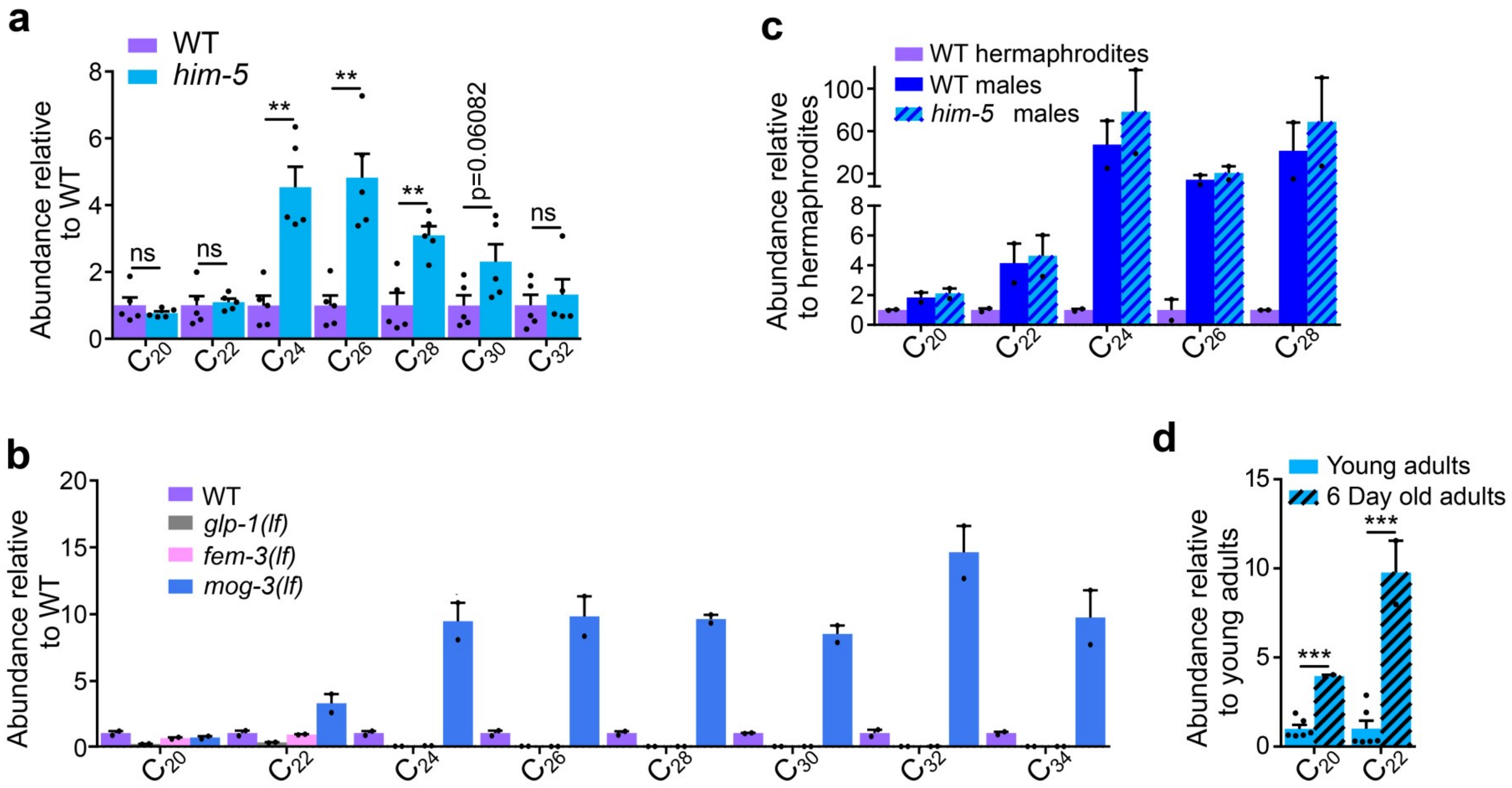


**Supplementary Figure 24. Male germline-dependent increase in polyunsaturated very long chain fatty acids (VLCPUFAs). a.** Penta-unsaturated VLCPUFAs with 24-28 carbons were significantly enriched in *him-5* cultures. **b.** VLCPUFAs with 24-32 carbons were enriched in germline-masculinized *mog-3(lf)* animals compared to WT, but absent in *glp-1 (lf)*and *fem-3(lf)* animals. Note that these measurements of lipophilic fatty acids used a different set of germline mutants than the other metabolomic comparisons (*mog-3(lf)* instead of *fem-3(gf)* and *fem-3(lf)* instead of *fem-2(lf)*. **c.** Although only two VLCPUFAs are detected in worm bodies extracted with methanol, EPA and DPA are 4- and 10-fold enriched in old worms over young worms. Data are presented as mean ± s.e.m. *, p≤0.05; **, p≤0.005; ***, p≤0.0005; ns, not statistically significant.

**Supplementary Tables**

**Supplementary Table 1. *him-5*-enriched compounds.**

If a molecular formula was not able to be determined from the *m/z* and isotope pattern, the *m/z* for the feature is included in the "Molecular Formula" column. If the compound was initially detected as enriched in the endo- or exo-metabolome of *him-5* samples it is denoted as a "P" for the pellet/endo-metabolome and/or "S" for the supernatant/exo-metabolome. If a metabolite was not detected in a sample it is denoted "ND". Fold-change values are derived from Metaboseek analysis and those provided for *fem-3* (gf) are relative to WT (N2) samples.

| Molecular Formula | Pellet/Super-natant | RT (min) | Fold Change | Identity | Up in *fem-3* (gf) | Up in Small Males | Up in 50:50? |
|---|---|---|---|---|---|---|---|
| $C_7H_{11}NO_3$ | S | 2.59 | 3.46 | | No | ND | ND |
| $C_{10}H_{13}NO_3$ | P | 4.52 | 3.04 | | No | 10.33 | No |
| $C_9H_{16}N_2SO$ | S | 5.27 | 2.14 | | 3.3 | No | ND |
| $C_9H_{19}NO_3$ | S | 5.34 | 4.11 | | No | ND | ND |
| $C_{20}H_{25}N_2PO_{11}$ | P | 5.39 | 2.07 | tyglu#501 | No | ND | No |
| $C_{16}H_{23}NO_7$ | P | 5.66 | 16.5 | No MS2 | No | No | ND |
| $C_{17}H_{25}NO_9$ | P | 5.66 | 32.1 | No MS2 | No | No | No |
| $C_{11}H_{12}N_2O_2$ | S | 5.7 | 2.4 | | No | No | ND |
| $C_{11}H_{11}NO_4$ | S | 5.78 | 2.55 | | No | ND | ND |
| $C_9H_{11}N_2PO_4$ | P | 5.8 | 2.05 | | No | No | No |
| $C_{18}H_{21}N_6PO_{10}$ | P | 5.8 | 2.76 | | No | ND | ND |
| $C_{17}H_{21}N_4PO_{10}$ | P | 5.94 | 2.16 | | No | ND | No |
| $C_{13}H_{12}N_2O_3$ | P | 5.98 | 2.19 | | No | ND | No |
| $C_{12}H_{11}N_2SO_5$ | S | 6.05 | 2.99 | | No | No | ND |
| $C_{23}H_{30}N_2PO_{15}$ | P | 6.08 | 2.28 | | No | ND | ND |
| 639.1559 [-] | P | 6.23 | 6.31 | | 1.86 | ND | ND |
| $C_{11}H_{20}O_5$ | S | 6.23 | 3.8 | | No | ND | ND |
| $C_{10}H_{17}NO_6$ | S | 6.25 | 2.8 | | No | ND | ND |
| $C_{13}H_{12}N_2O_3$ | P | 6.26 | 2.35 | | No | ND | No |
| 628.1308 [-] | P | 6.27 | 60.6 | No MS2 | No | ND | ND |
| $C_{21}H_{27}N_2PO_{10}$ | P | 6.32 | 2.22 | tyglu#2 | No | No | 1.21 |
| $C_{26}H_{35}N_2PO_{12}$ | P | 6.33 | 289.33 | No MS2 | No | ND | ND |

| | | | | | | | |
|---|---|---|---|---|---|---|---|
| 619.1799 [+] | P | 6.43 | 7.17 | | No | ND | 1.13 |
| $C_{19}H_{27}N_3PO_{11}$ | P | 6.52 | 2.16 | | No | ND | ND |
| $C_{27}H_{44}NPO_{14}$ | P | 6.58 | 3.81 | | No | ND | ND |
| $C_{23}H_{35}N_4PO_{16}$ | P | 6.58 | 20.52 | No MS2 | No | ND | ND |
| $C_{10}H_7NO_2$ | P | 6.59 | 4.87 | | No | No | 1.26 |
| $C_9H_{14}N_2O_4$ | S | 6.59 | 2.27 | | No | ND | ND |
| $C_{15}H_{32}NPO_8$ | S | 6.59 | 4.03 | | No | ND | ND |
| $C_{20}H_{36}N_4O_6$ | S | 6.65 | 2.41 | | No | ND | ND |
| $C_{16}H_{24}N_2O_3$ | S | 6.66 | 20.59 | | 2.59 | No | ND |
| $C_{13}H_{16}N_2O_4$ | P | 6.7 | 3.07 | | 4.61 | 3.74 | No |
| $C_{26}H_{53}N_{10}PO_5$ | S | 6.7 | 2.05 | | No | ND | ND |
| $C_{13}H_{25}PO_{11}$ | P | 6.71 | 25.18 | No MS2 | No | ND | ND |
| $C_{19}H_{35}PO_{14}$ | P | 6.72 | 21.94 | glas#11 | No | ND | ND |
| $C_{26}H_{28}N_3PO_{11}$ | P | 6.73 | 2.31 | | No | ND | ND |
| $C_{21}H_{26}NPO_{11}$ | P | 6.75 | 12.33 | oglu derivative? | No | 8.03 | ND |
| $C_6H_{11}PO_6$ | S | 6.76 | 3.79 | | No | ND | ND |
| $C_{13}H_{25}PO_9$ | S | 6.76 | 3.84 | phascr#11 | No | No | ND |
| $C_{14}H_{28}N_4O_{13}$ | P | 6.8 | 3.03 | | No | ND | ND |
| $C_{18}H_{25}N_4PO_{13}$ | P | 6.83 | 6.86 | gluric#2-derivative? | No | ND | ND |
| $C_{27}H_{37}N_2PO_{13}$ | P | 6.84 | 29.05 | anglas#? | No | ND | ND |
| $C_{19}H_{34}O_{11}$ | P | 6.86 | 17.11 | glas#1 | No | No | ND |
| $C_{20}H_{36}O_{13}$ | P | 6.86 | 19.39 | ascr#1-heptulose? | No | No | ND |
| $C_{18}H_{35}N_5O_7$ | S | 6.87 | 2.39 | | No | No | ND |
| $C_{24}H_{28}N_3PO_{11}$ | P | 6.89 | 2.39 | tyglu#24 | No | ND | No |
| $C_{11}H_{14}N_3P$ | S | 6.89 | 2.24 | | No | No | ND |
| $C_{28}H_{30}N_3PO_{12}$ | P | 6.9 | 2.24 | | No | ND | ND |
| $C_{13}H_{25}PO_{11}$ | P | 6.9 | 39.02 | No MS2 | No | 2.47 | ND |
| $C_{19}H_{35}PO_{14}$ | P | 6.91 | 55.36 | glos#11 | No | ND | ND |
| $C_{27}H_{35}PO_{13}$ | P | 6.95 | 2.12 | | No | ND | ND |
| $C_{18}H_{25}N_4PO_{13}$ | P | 7 | 5.44 | gluric#2-derivative? | No | ND | ND |

| | | | | | | | |
|---|---|---|---|---|---|---|---|
| $C_{19}H_{34}O_{11}$ | S | 7 | 3.52 | glos#1 | No | ND | ND |
| $C_{12}H_{22}N_2O_2$ | S | 7.02 | 3.59 | | No | No | ND |
| $C_9H_{10}N_2O$ | S | 7.04 | 2.3 | | No | ND | ND |
| $C_{18}H_{27}N_4PO_{13}$ | P | 7.07 | 77.82 | gluric#2-derivative? | No | No | 1.31 |
| $C_{20}H_{28}NPO_{11}$ | P | 7.08 | 54.3 | No MS2 | No | 1.82 | ND |
| $C_{24}H_{37}N_4PO_{16}$ | P | 7.08 | 75.18 | uglas#15 | No | 38.87 | 1.25 |
| $C_{29}H_{37}N_2PO_{13}$ | P | 7.09 | 46.07 | oglu? | No | ND | ND |
| $C_{24}H_{36}N_4O_{13}$ | P | 7.16 | 23.01 | uglas#14 | No | 6.39 | 1.48 |
| $C_{21}H_{27}N_2PO_{10}$ | P | 7.22 | 46.07 | tyglu#2? | No | No | ND |
| $C_{14}H_{22}NPO_9$ | P | 7.22 | 66.04 | No MS2 | No | ND | ND |
| $C_{19}H_{28}NPO_{11}$ | P | 7.24 | 8.36 | | No | ND | No |
| $C_{24}H_{25}N_2PO_9$ | P | 7.25 | 2.21 | | No | ND | ND |
| $C_{10}H_{19}PO_7$ | S | 7.28 | 2.11 | | No | ND | ND |
| $C_{28}H_{30}N_3PO_{12}$ | P | 7.29 | 2.74 | tyglu#6? | No | No | No |
| $C_{27}H_{44}NPO_{14}$ | P | 7.29 | 146.6 | anglas#-derivative? | No | ND | ND |
| $C_{17}H_{20}N_2PO_{11}$ | P | 7.3 | 3.28 | | No | ND | ND |
| $C_{19}H_{28}NPO_{10}$ | P | 7.31 | 2.8 | Glucoside - tyramine tiglic acid | No | 2.11 | ND |
| $C_{20}H_{24}N_5PO_9$ | P | 7.32 | 23.01 | maglu#3 | No | ND | ND |
| 613.2157 [+] | P | 7.37 | 33.85 | No MS2 | No | No | No |
| $C_{32}H_{46}N_2PO_{15}$ | P | 7.44 | 2.34 | | No | ND | ND |
| $C_{17}H_{26}N_2O_3$ | S | 7.45 | 13.07 | | No | ND | ND |
| $C_{15}H_{27}NO_5$? | S | 7.47 | 2.75 | | No | No | ND |
| $C_{19}H_{21}N_4PO_{12}$ | P | 7.5 | 11.1 | gluric#2 | No | ND | ND |
| 663.4187 [+] | P | 7.53 | 43.84 | No MS2 | No | ND | ND |
| $C_{28}H_{54}N_6O_7$? | S | 7.54 | 2.24 | | No | ND | ND |
| $C_{17}H_{29}N_3O_5$ | S | 7.54 | 2.57 | | 3.61 | 1.89 | ND |
| $C_{19}H_{30}NPO_{10}$? | P | 7.55 | 3.78 | tyglu#9 | 2.12 | ND | ND |
| $C_{19}H_{28}NPO_{14}$ | P | 7.58 | 5.67 | | 1.75 | ND | ND |
| 743.2785 [+] | P | 7.6 | 12.55 | No MS2 | No | ND | ND |

| | | | | | | | |
|---|---|---|---|---|---|---|---|
| $C_{19}H_{27}N_4PO_{13}$ | P | 7.6 | 15.11 | | No | ND | ND |
| $C_{13}H_{24}O_6$ | P | 7.63 | 8.31 | ascr#1 | No | No | 2.52 |
| $C_{20}H_{24}N_5PO_{10}$ | P | 7.63 | 8.57 | mgglu#5 | No | ND | ND |
| $C_{31}H_{51}N_{10}PO_6$ | S | 7.63 | 3.37 | | No | ND | ND |
| $C_9H_{17}N_3O_{13}$ | P | 7.64 | 5.26 | | No | No | 1.77 |
| $C_{10}H_{19}N_3O_{15}$ | P | 7.64 | 6.58 | | No | No | 3.86 |
| $C_{14}H_{13}N_5O_3$ | P | 7.64 | 7.33 | | No | No | 1.89 |
| $C_{13}H_{17}NO_4$ | P+S | 7.66 | 6.06 | angl#1 | No | ND | ND |
| $C_6H_8O_7$ | S | 7.67 | 2.36 | | No | ND | ND |
| $C_{20}H_{28}NPO_{11}$ | P | 7.7 | 10.57 | | No | 1.22 | ND |
| $C_{26}H_{37}N_5PO_{12}$ | P | 7.75 | 40.23 | No MS2 | No | ND | ND |
| 636.3602 [+] | P | 7.81 | 27.85 | No MS2 | No | ND | ND |
| $C_{22}H_{28}NPO_{10}$ | P | 7.84 | 32.83 | No MS2 | 1.61 | ND | ND |
| $C_{25}H_{31}N_2PO_{11}$ | P | 7.85 | 2.95 | | No | ND | ND |
| $C_{25}H_{38}NPO_{15}$ | P | 7.85 | 22.88 | nicotinic glucoside | No | ND | ND |
| $C_{19}H_{27}N_3O_3$ | S | 7.94 | 2.1 | medip#1 | 490.9 | ND | 13.71 |
| $C_{23}H_{33}NO_{10}$ | S | 7.94 | 4.19 | osas#9 | No | ND | ND |
| 755.2768 [+] | P | 7.97 | 7.18 | | No | ND | ND |
| $C_{10}H_{19}NSO_3$ | S | 7.97 | 2.53 | | No | No | ND |
| $C_{28}H_{45}N_5O_7$ | P | 8 | 44.52 | Peptide – LGLVT/VALVT? | No | No | ND |
| $C_{30}H_{34}N_4O_{11}$ | P | 8.03 | 62.53 | No MS2 | No | ND | ND |
| $C_{10}H_{19}NO_3$ | P | 8.04 | 2.51 | | No | No | No |
| $C_{34}H_{49}N_2PO_{15}$ | P | 8.05 | 46.64 | tyglas#1 | No | ND | ND |
| $C_{13}H_{27}N_6PO_6$ | P | 8.06 | 2.68 | | No | ND | ND |
| $C_{20}H_{32}N_4PO_{13}$ | P | 8.14 | 2.91 | | No | No | 1.13 |
| $C_{26}H_{41}N_4PO_{16}$ | P | 8.14 | 3.22 | uglas#105 | No | 2396 | 21.605 |
| $C_{25}H_{38}NPO_{15}$ | P | 8.21 | 5.77 | | No | ND | ND |
| $C_{28}H_{32}N_3PO_{11}$ | P | 8.23 | 2.6 | tyglu#4? | No | No | 5.13 |
| $C_{26}H_{40}N_4O_{13}$ | P | 8.24 | 2.21 | uglas#104 | No | 302.3 | 5.15 |

| | | | | | | | |
|---|---|---|---|---|---|---|---|
| $C_{26}H_{33}N_2PO_{11}$ | P | 8.24 | 2.6 | tyglu#8 | No | No | 45.3 |
| $C_{29}H_{48}NPO_{14}$ | P | 8.24 | 1560.59 | No MS2 | No | No | ND |
| $C_{27}H_{33}N_2PO_{12}$ | P | 8.25 | 2.22 | angl#34? | No | ND | ND |
| $C_{13}H_{23}NO_3$ | S | 8.37 | 2.31 | | No | ND | ND |
| $C_{20}H_{31}N_3O_8$ | S | 8.42 | 2.8 | | No | ND | ND |
| $C_{18}H_{29}N_3O_6$ | S | 8.44 | 2.03 | | No | ND | ND |
| 482.1238 [-] | S | 8.44 | 2.14 | | No | ND | ND |
| $C_{11}H_{18}O_4$ | S | 8.52 | 4.85 | bemeth#5.1 | 3.05 | ND | ND |
| $C_{11}H_{18}O_3$ | S | 8.54 | 3.22 | | 1.72 | No | ND |
| $C_{11}H_{20}O_4$ | S | 8.54 | 5.28 | bemeth#3.1 | 2.39 | No | 1.76 |
| $C_{11}H_{18}O_5$ | S | 8.54 | 7.44 | bemeth#6 | 1.55 | No | ND |
| $C_{22}H_{33}N_4PO_{13}$ | P | 8.64 | 2.08 | | 1.45 | ND | ND |
| $C_{11}H_{20}O_4$ | S | 8.7 | 2.72 | bemeth#3.2 | 2.37 | No | 1.47 |
| $C_{28}H_{31}N_6PO_7$ | P | 8.71 | 6.05 | | 1.2 | ND | 1.3 |
| $C_{14}H_{19}NO_4$ | S | 8.78 | 2.2 | | No | ND | ND |
| $C_{20}H_{29}NO_7$ | S | 8.78 | 2.5 | ascr#801 | No | ND | ND |
| $C_{30}H_{43}N_2PO_{12}$ | P | 8.8 | 15.79 | No MS2 | No | ND | ND |
| $C_{25}H_{46}NPO_{13}$ | S | 8.92 | 2.62 | | No | ND | ND |
| $C_{20}H_{30}NPO_{12}$ | P | 9.15 | 30.48 | No MS2 | No | No | ND |
| $C_{26}H_{40}NPO_{15}$ | P | 9.15 | 40.98 | anglas#2 | No | ND | ND |
| $C_{21}H_{33}N_3O_8$ | S | 9.16 | 2.24 | | No | ND | ND |
| 679.2625 [+] | P | 9.18 | 11.51 | | No | ND | No |
| $C_{14}H_{21}NO_3$ | S | 9.21 | 2.06 | | No | ND | ND |
| 679.2625 [+] | P | 9.26 | 13.86 | | No | ND | ND |
| $C_{14}H_{21}NO_4$ | S | 9.27 | 2.72 | | 1.4 | 1.32 | ND |
| $C_{11}H_{18}N_2SO_3$ | S | 9.37 | 2.09 | | No | No | ND |
| 743.2422 [+] | P | 9.65 | 16.3 | | No | No | ND |
| $C_{19}H_{26}NPO_{11}$ | P | 9.66 | 10.62 | No MS2 | No | No | ND |
| $C_{27}H_{48}NPO_{13}$ | S | 9.76 | 2.3 | | No | ND | ND |
| $C_{11}H_{23}NO_2$ | S | 9.81 | 2.857 | | 2.74 | No | ND |

| | | | | | | | |
|---|---|---|---|---|---|---|---|
| $C_{11}H_{16}O_4$ | S | 10.05 | 3.44 | | No | No | ND |
| $C_{25}H_{44}O_{10}$ | S | 10.23 | 2.75 | | No | ND | ND |
| $C_{18}H_{29}NO_3$ | S | 10.27 | 2.02 | | No | ND | ND |
| $C_{25}H_{44}O_{10}$ | S | 10.36 | 2.06 | | No | ND | ND |
| 681.2781 [+] | P | 10.55 | 3.93 | | 1.31 | ND | ND |
| $C_{11}H_{13}PO_5$ | P | 10.57 | 2.18 | | No | No | No |
| $C_{13}H_{16}PO_5$ | P | 10.57 | 2.64 | | No | No | 1.74 |
| $C_{15}H_{30}NPO_8$ | P | 10.74 | 10.39 | | No | ND | ND |
| 779.3510 [+] | P | 10.77 | 22.12 | No MS2 | No | ND | ND |
| $C_{16}H_{30}O_6$ | P | 10.78 | 2.47 | ascr#16? | No | ND | 14.06 |
| $C_{15}H_{30}NPO_8$ | P | 10.85 | 11.83 | No MS2 | No | ND | ND |
| 649.2480 [+] | P | 10.88 | 4.45 | | No | ND | ND |
| $C_{43}H_{52}N_2O_{10}$ | P | 10.89 | 13.92 | | No | ND | ND |
| $C_{17}H_{33}NO_4$ | S | 10.9 | 3.1 | ω-OH C11 – crotonobetaine? | 2.01 | ND | ND |
| $C_{31}H_{54}NPO_{13}$ | P | 10.91 | 5.25 | | No | ND | ND |
| 741.3670 [2+] | P | 10.92 | 85.95 | No MS2 | No | ND | ND |
| $C_{25}H_{40}NPO_{10}$ | P | 10.99 | 24.66 | tyglu#54/55 | No | ND | ND |
| $C_{27}H_{44}NPO_{10}$ | P | 11.13 | 16.17 | | No | ND | ND |
| $C_{16}H_{20}O_6$ | S | 11.19 | 2.13 | | No | ND | ND |
| $C_{13}H_{20}O_5$ | S | 11.21 | 2.07 | | No | ND | ND |
| $C_{36}H_{59}N_2PO_{13}$ | P | 11.22 | 17.18 | No MS2 | No | ND | ND |
| $C_{28}H_{48}O_{11}$ | S | 11.24 | 2.86 | | No | ND | ND |
| $C_{16}H_{20}O_6$ | S | 11.27 | 2.24 | | No | ND | ND |
| $C_{11}H_{19}NO_3$ | P | 11.33 | 8.75 | | 5.49 | 4.48 | ND |
| $C_{36}H_{59}N_2O_{13}$ | P | 11.34 | 10.13 | | No | ND | ND |
| $C_{28}H_{48}O_{11}$ | S | 11.37 | 2.94 | dasc#13 | No | ND | ND |
| $C_{30}H_{47}N_2PO_{12}$ | P | 11.46 | 6.97 | | No | ND | ND |
| $C_{38}H_{44}N_2O_8$ | P | 11.49 | 6.59 | | No | ND | ND |
| $C_{17}H_{29}PO_{10}$ | P | 11.52 | 7.24 | | 3.02 | ND | ND |

| | | | | | | | |
|---|---|---|---|---|---|---|---|
| $C_{27}H_{43}N_4PO_{13}$ | P | 11.55 | 16.92 | No MS2 | No | ND | ND |
| $C_{19}H_{32}O_6$ | P | 11.98 | 2.28 | | No | ND | ND |
| $C_{34}H_{49}N_2PO_{11}$ | P | 12.01 | 22.24 | Glucoside? | No | ND | ND |
| $C_{32}H_{50}NPO_{11}$ | P | 12.01 | 15.53 | Glucoside? | No | ND | ND |
| 299.1031 [+] | S | 12.09 | 2.17 | | No | ND | ND |
| $C_{28}H_{45}N_4PO_{13}$ | P | 12.10 | 15.49 | | No | ND | ND |
| 687.2657 [-] | P | 12.13 | 25.48 | No MS2 | No | ND | ND |
| 577.1922 [-] | P | 12.15 | 3.82 | | No | ND | ND |
| 675.2654 [-] | P | 12.25 | 10.20 | | No | ND | ND |
| $C_{12}H_{21}NO_3$ | P | 12.27 | 5.79 | | 8.22 | ND | ND |
| $C_{27}H_{44}NPO_{10}$ | P | 12.28 | 37.72 | No MS2 | No | ND | ND |
| $C_{22}H_{39}PO_{12}$ | P | 12.66 | 2.06 | | No | ND | ND |
| $C_{23}H_{43}PO_{11}$ | P | 12.67 | 5.99 | | No | ND | ND |
| 691.2925 [+] | P | 12.71 | 66.46 | No MS2 | No | ND | ND |
| $C_{11}H_{14}O_3$ | S | 12.74 | 2.71 | | No | ND | ND |
| $C_{18}H_{22}N_2O_8$ | P | 13.13 | 2.26 | | No | No | ND |
| $C_{12}H_{16}O_3$ | S | 13.42 | 2.01 | | No | ND | ND |
| $C_{24}H_{47}N_3O_2$ | P | 13.57 | 5.75 | | No | ND | ND |
| $C_{23}H_{31}N_2PO_7$ | P | 13.72 | 3.08 | | No | ND | ND |
| $C_{26}H_{37}NO_7$ | P | 13.89 | 2.09 | icas#18? | No | No | ND |
| $C_{26}H_{55}NO_8$ | P | 14.72 | 3.69 | | No | ND | ND |
| $C_{29}H_{62}N_6O_6$ | P | 15.05 | 4.71 | | No | ND | ND |
| $C_{22}H_{26}N_2O_5$ | P | 15.27 | 2.23 | | No | No | ND |
| $C_{25}H_{47}NO_4$ | P | 15.75 | 2.25 | | No | No | ND |
| $C_{29}H_{58}NPO_9$ | S | 16.18 | 4.07 | | No | ND | ND |
| $C_{25}H_{49}NO_4$ | P | 16.59 | 2.75 | | No | No | ND |
| $C_{29}H_{62}NPO_9$ | S | 16.76 | 3.29 | | No | No | ND |
| $C_{26}H_{51}NO_4$ | P | 17.05 | 2.19 | | No | 2.21 | ND |
| $C_{28}H_{55}NO_4$ | P | 17.94 | 2.82 | | No | No | ND |
| $C_{29}H_{57}NO_4$ | P | 18.32 | 2.18 | | No | No | ND |

| | | | | | | | |
|---|---|---|---|---|---|---|---|
| 746.4602[+] | P+S | 18.60 | 4.39 | PE/PC | No | ND | ND |
| $C_{28}H_{54}N_2O_5$ | S | 20.21 | 2.21 | | No | ND | ND |
| 544.4351[-] | S | 20.54 | 2.26 | | No | ND | ND |
| $C_{24}H_{30}O_3$ | P | 21.12 | 2.86 | | No | ND | ND |
| $C_{35}H_{61}N_3O_6$ | S | 21.19 | 2.67 | | No | ND | ND |
| $C_{37}H_{67}NO_4$ | S | 22.72 | 2.31 | | No | ND | ND |
| $C_{36}H_{67}NO_4$ | S | 22.89 | 2.00 | | No | ND | ND |
| $C_{33}H_{66}N_2O_9$ | P | 23.60 | 2.03 | | No | ND | ND |
| 763.5134 [+] | P | 23.68 | 674.84 | No MS2 | No | No | ND |
| $C_{37}H_{72}O_6$ | S | 24.66 | 2.13 | | No | No | ND |
| 779.6343 [+] | S | 25.31 | 2.11 | | No | No | ND |

**Supplementary Table 2.** ***fem-3*** **(gf)-enriched compounds.**

If a molecular formula was not able to be determined from the *m/z* and isotope pattern, the *m/z* for the feature is included in the "Molecular Formula" column. If the compound was initially detected as enriched in the endo- or exo-metabolome of *fem-3* (gf) samples it is denoted as a "P" for the pellet/endo-metabolome and/or "S" for the supernatant/exo-metabolome. If a metabolite was not detected in a sample, it is denoted "ND". Metabolites not enriched the metabolome of young *him-5* cultures that are enriched in the metabolome of older *him-5* cultures are denoted with a "*".

| Molecular Formula | Pellet /SN | RT (min) | Identity | Fold over N2 | Fold over *fem-2* (lf) | Fold over *glp-4* | Up in *him-5* | Up in Small Males | Up in 50:50 |
|---|---|---|---|---|---|---|---|---|---|
| $C_{10}H_{12}N_5PO_7$ | P | 2.55 | | 2.64 | 3.18 | 2.07 | No | ND | ND |
| $C_{11}H_{22}N_2O_4$ | P | 4.68 | | 15.18 | 42.6 | 46.22 | ND | ND | ND |
| $C_{15}H_{26}O_{11}$ | P | 4.79 | | 2.34 | 8558 | 421 | 5.41* | 26.29 | ND |
| $C_{15}H_{26}O_{11}$ | P | 5.19 | | 2.75 | 19746 | 2022 | 52.82* | 61.64 | ND |
| 479.1735 [+] | P | 5.24 | | 1.1 | 5250 | 2578 | 2.93* | 12.47 | ND |
| $C_{10}H_{18}N_2O_4$ | S | 5.79 | | 3.21 | 46.37 | 124.4 | 1.64 | ND | No |
| $C_{24}H_{38}O_9$ | P | 5.81 | | 9.02 | 15.44 | 21.67 | 7.32* | ND | No |
| 469.0656 [-] | P | 5.97 | | 0.79 | 3.53 | 2.1 | 13.82* | ND | ND |
| $C_{15}H_{24}NO_4$ | S | 6.61 | | 3.26 | 39.7 | 48.49 | 13.11* | No | ND |
| $C_{13}H_{16}N_2O_4$ | P+S | 6.72 | | 5.07 | 51.96 | 75.31 | 2.97 | 52.88 | No |
| $C_{13}H_{17}N_5SO_4$ | P | 6.77 | acemta#1 | 19.22 | 132.9 | 211.2 | 1.75 | 2.37 | ND |
| $C_8H_{15}NO_3$ | S | 6.91 | | 2.4 | 3.9 | 18.9 | No | No | No |
| $C_{16}H_{30}N_6O_6$ | S | 7.18 | | 4.58 | 2058 | 4684 | 2.1 | ND | 4.89 |
| $C_{21}H_{36}NPO_{17}$ | P | 7.30 | | 5.94 | 38.02 | 51 | 9.01* | ND | ND |
| $C_{20}H_{24}N_5PO_9$ | P | 7.34 | maglu#3 | 3.75 | 5.22 | 2.95 | 5.56 | ND | ND |
| $C_{13}H_{17}N_5SO_4$ | P | 7.39 | acemta#2 | 16.31 | 95.33 | 182 | 1.75 | 2.37 | ND |
| $C_{19}H_{21}N_4PO_{12}$ | P | 7.47 | gluric#3 | 6.1 | 22.89 | 3.59 | 3.44 | ND | ND |
| $C_{14}H_{27}N_5O_6$ | S | 7.49 | | 2.44 | 92.57 | 4.38 | 7.18* | ND | ND |
| $C_{14}H_{28}N_6O_5$ | P | 7.56 | | 8.85 | 252.3 | 571.9 | No | 1.79 | ND |
| $C_{24}H_{21}N_3PO_{12}$ | S | 7.57 | | 1.13 | 2.09 | 2.74 | 6.22* | ND | ND |
| $C_{19}H_{28}NPO_{14}$ | P | 7.58 | tyglu#9 | 2.12 | 4.89 | 2.71 | 3.62 | No | ND |
| $C_{20}H_{24}N_5PO_{10}$ | P | 7.61 | mgglu#5 | 2.33 | 5.41 | 4.85 | 5.67 | No | ND |
| $C_{16}H_{30}N_6O_7$ | S | 7.65 | | 4.53 | 1260 | 370 | 139.6* | 20.99 | 1.74 |

| | | | | | | | | | |
|---|---|---|---|---|---|---|---|---|---|
| $C_{19}H_{23}N_3O_4$ | S | 7.78 | | 5.41 | 5415 | 4772 | 1.59 | 39.55 | 3.31 |
| $C_{13}H_{29}PO_{12}$ | P | 7.79 | | 2.58 | 4.6 | 4.49 | No | ND | No |
| $C_{14}H_{27}N_5O_6$ | S | 7.82 | | 1.87 | 92.57 | 39.49 | 2.05 | ND | ND |
| $C_{19}H_{27}N_3O_3$ | S | 7.87 | medip#1 | 3.74 | 4526 | 380 | 1.33 | 1.97 | 15.25 |
| $C_{15}H_{29}N_5O_5$ | S | 8.07 | | 3.85 | 6.45 | 48.57 | ND | ND | No |
| $C_8H_{16}N_8O_2$ | S | 8.09 | | 2.48 | 19.59 | 19.41 | ND | ND | ND |
| $C_{16}H_{22}NO_{17}$ | P | 8.16 | | 2.5 | 4.67 | 8.74 | 34.91* | ND | ND |
| $C_{12}H_{20}N_2O_4$ | S | 8.20 | | 2.09 | 182.53 | 2797 | 38.49* | ND | No |
| $C_{11}H_{20}O_4$ | S | 8.29 | | 9.02 | 12.93 | 48.59 | 2.91 | ND | ND |
| $C_{11}H_{20}O_4$ | S | 8.35 | | 1.57 | 12.18 | 59.97 | ND | ND | ND |
| $C_{19}H_{27}N_3O_4$ | S | 8.39 | medip#2 | 7.15 | 248.3 | 48.57 | 17.03* | 1.82 | 1.72 |
| $C_{10}H_{18}NO_4$ | S | 8.45 | | 17.87 | 680.9 | 3388 | 50.27* | ND | No |
| $C_{14}H_{25}NO_5$ | S | 8.50 | alanine bemeth | 15.45 | 134.99 | 870 | 2.53 | ND | No |
| $C_{17}H_{20}N_2O_3$ | S | 8.53 | | 13.39 | 1129 | 818.2 | 42.91* | ND | ND |
| $C_{11}H_{18}O_4$ | S | 8.54 | bemeth#5.1 | 2.62 | 9.19 | 42.15 | 7.53 | ND | ND |
| $C_{11}H_{18}O_3$ | S | 8.55 | | 1.96 | 8.25 | 10.25 | 3.25 | ND | ND |
| 304.0169 [-] | S | 8.56 | | 5.41 | 578.3 | 2208 | 8.3 | ND | ND |
| $C_{11}H_{20}O_4$ | S | 8.57 | bemeth#3.1 | 2.39 | 174.5 | 513.1 | 7.53 | 2.01 | 1.76 |
| $C_{14}H_{25}NO_5$ | S | 8.69 | alanine bemeth | 15.45 | 161.7 | 566.9 | 1.84 | 6.48 | 1.9 |
| $C_{11}H_{20}O_4$ | S | 8.72 | bemeth#3.2 | 2.37 | 11.83 | 131.5 | 5.45 | 4.01 | 1.47 |
| 498.1528 [+] | P | 8.72 | | 1.35 | 2.21 | 2.12 | 32.83 | 14.76 | No |
| $C_{28}H_{31}N_6PO_7$ | P | 8.72 | | 0.68 | 2.24 | 2.12 | 6.05 | ND | No |
| 484.1005 [+] | P | 8.91 | | 2.08 | 3.46 | 5.36 | 13.56* | ND | ND |
| 482.0848 [+] | P | 8.96 | | 1.22 | 2.14 | 2.37 | 56.78* | ND | ND |
| $C_{11}H_{20}O_4$ | S | 8.99 | | 4.09 | 37.08 | 42.78 | 4.1 | ND | ND |
| $C_{11}H_{20}O_4$ | S | 9.06 | | 4.94 | 10.28 | 34.38 | 4.28 | ND | ND |
| $C_{14}H_{25}NO_4$ | S | 9.13 | | 4.72 | 12.59 | 92.82 | 3.48 | ND | 1.39 |
| $C_7H_{17}N_7$ | S | 9.33 | | 0.83 | 35.41 | 44.09 | 1.91 | No | 3.89 |

| | | | | | | | | | |
|---|---|---|---|---|---|---|---|---|---|
| $C_{11}H_{21}NO_4$ | S | 9.48 | | 19.68 | 48.17 | 5792 | 16.34* | ND | No |
| $C_{10}H_{24}N_2O_3$ | S | 9.52 | | 6.3 | 45.14 | 22.59 | 2.11 | ND | ND |
| $C_{11}H_{23}NO_2$ | S | 9.74 | | 4.39 | 16.73 | 11.12 | 3.18 | No | No |
| 574.2268 [+] | P | 9.89 | | 2.04 | 11258 | 331.2 | 16.18 | ND | ND |
| $C_{21}H_{36}NPO_{17}$ | P | 9.98 | | 508.3 | 10788 | 5299 | 229.2* | ND | ND |
| $C_{31}H_{41}N_2PO_{12}$ | P | 10.02 | | 4.69 | 55.86 | 44.83 | 52.84* | ND | ND |
| 501.1267 [+] | P | 10.02 | | 3.44 | 20.67 | 22.762 | No | ND | ND |
| $C_{22}H_{33}N_4PO_{13}$ | P | 10.06 | | 3.81 | 23668 | 11630 | 4.68 | ND | ND |
| 656.2957 [+] | P | 10.10 | | 3.95 | 2.64 | 2.69 | No | ND | ND |
| $C_{22}H_{27}N_9O_{11}$ | P | 10.14 | | 2.25 | 2085 | 486.42 | 651.8* | ND | ND |
| $C_{15}H_{22}NO_4$ | P | 10.28 | nacq#1 | 4.17 | 500 | 260.5 | 1.37 | 11.18 | 3.02 |
| $C_{10}H_{17}NO_3$ | S | 10.29 | | 9.69 | 693.1 | 4008 | 13.60 | 3.42 | No |
| 466.0899 [+] | P | 10.39 | | 2.19 | 3.98 | 5.9 | 23.16* | ND | No |
| $C_{11}H_{26}N_2O_3$ | S | 10.40 | | 11.66 | 193.7 | 120.22 | ND | No | ND |
| 498.1159 [+] | P | 10.43 | | 1.26 | 3.07 | 3.7 | 173.9* | No | ND |
| $C_{11}H_{20}O_4$ | S | 10.66 | bemeth#33 | 1.74 | 1891 | 11.2 | No | No | No |
| $C_{11}H_{20}O_4$ | S | 10.75 | bemeth#34 | 3.09 | 5.22 | 4.39 | No | No | No |
| 466.0892 [+] | P | 10.92 | | 1 | 2.57 | 5.9 | 5.69* | ND | ND |
| $C_{16}H_{27}N_8PO_{15}$ | P | 10.92 | | 1 | 2.69 | 9.27 | 14.79* | ND | ND |
| $C_{20}H_{17}NO_6$ | P | 10.92 | | 1.11 | 2.21 | 6.67 | 20.07* | ND | ND |
| $C_{15}H_{25}N_2O_4$ | P | 11.03 | | 4.31 | 8347 | 91.56 | 311.4* | ND | No |
| 514.1470 [+] | P | 11.20 | | 4.3 | 9028 | 289.9 | 15.21* | ND | ND |
| $C_{11}H_{19}NO_3$ | P | 11.23 | glycine lipid conjugate | 21.47 | 34.88 | 43.37 | 8.07* | 4.48 | No |
| $C_{14}H_{25}NO_5$ | S | 11.24 | bemeth#71 | 6.61 | 3162 | 1089 | 2.5 | 208.0 | 2.83 |
| $C_{11}H_{19}NO_3$ | P | 11.34 | | 9.47 | 11.23 | 7.59 | 8.75 | ND | No |
| $C_{14}H_{25}NO_5$ | S | 11.46 | bemeth#72 | 1.82 | 16.26 | 28.23 | 1421* | 1.53 | ND |
| 447.1386 [+] | P | 11.53 | | 5939 | 6672 | 2697 | ND | ND | ND |
| $C_{13}H_{23}NO_4$ | S | 11.68 | | 4.45 | 87503 | 683.3 | 2.39 | ND | ND |
| $C_{17}H_{29}PO_{10}$ | P | 11.77 | phosphogluc | 37.35 | 335.9 | 312.6 | 6.48 | ND | 4.31 |

| | | | | | | | | | |
|---|---|---|---|---|---|---|---|---|---|
| | | | ose conjugate | | | | | | |
| 516.1622 [+] | P | 11.90 | | 18.19 | 6945 | 3417 | No | ND | ND |
| 530.1785 [+] | P | 12.06 | | 10.29 | 58.81 | 15.79 | 67.71* | ND | ND |
| $C_{14}H_{23}NO_4$ | S | 12.08 | bemeth#81 | 20.34 | 9052 | 7344 | 2.72 | 1.44 | 1.77 |
| $C_{11}H_{18}O_3$ | S | 12.26 | bemeth | 2.62 | 24.32 | 65.54 | 15.76* | ND | ND |
| $C_{12}H_{21}NO_3$ | P | 12.35 | | 12.61 | 14.89 | 12.73 | 5.79 | ND | ND |
| $C_{14}H_{25}NO_4$ | S | 12.37 | bemeth#61 | 4 | 9413 | 1941 | 2.67* | 7.01 | 1.54 |
| 530.1781 [+] | P | 12.52 | | 143.4 | 1368 | 881.5 | 24.88* | ND | ND |
| $C_{14}H_{25}NO_4$ | S | 12.57 | bemeth | 1.32 | 1075 | 322.3 | 92.76* | ND | No |
| $C_{11}H_{20}O_3$ | S | 12.62 | bemeth#22 | 1.26 | 4.05 | 42.38 | ND | ND | ND |
| $C_{11}H_{20}O_3$ | S | 12.77 | bemeth#23 | 1.3 | 27.16 | 30.17 | 13.99* | No | No |
| $C_{14}H_{23}NO_3$ | S | 13.15 | | 3.11 | 11.52 | 11.22 | 27.07* | ND | ND |
| $C_{11}H_{22}O_3$ | S | 13.26 | | 1.91 | 2.15 | 2.3 | ND | ND | ND |
| $C_{11}H_{20}O_3$ | S | 13.83 | bemeth#24 | 3.2 | 22.32 | 27.83 | ND | ND | ND |
| $C_{18}H_{41}N_9O_9$ | P | 15.20 | | 2.67 | 5.18 | 2.1 | No | No | No |
| 570.3547 [+] | P | 15.32 | | 1 | 2.18 | 6.07 | No | No | No |
| 570.3547 [+] | P | 15.53 | | 2.73 | 2.55 | 3.96 | No | No | No |
| $C_{18}H_{43}N_9O_9$ | P | 15.82 | | 2.45 | 8.9 | 3.03 | ND | ND | No |
| 556.3394 [+] | P | 16.31 | | 3.66 | 12.2 | 3.17 | No | No | No |
| 650.4020 [+] | P | 16.72 | | 4.39 | 4.71 | 3.29 | ND | No | ND |
| 706.4642 [+] | P | 17.86 | | 2.83 | 2.47 | 3.21 | No | ND | ND |
| 674.4380 [+] | P | 17.87 | | 2.54 | 2.01 | 1.57 | No | ND | ND |

**Supplementary Table 3. Hand-picked males over hand-picked hermaphrodites.**

If a molecular formula was not able to be determined from the *m/z* and isotope pattern, the *m/z* for the feature is included in the "Molecular Formula" column. If a metabolite was not detected in a sample, it is denoted "ND". Fold-change values are derived from Metaboseek analysis and those provided for *fem-3* (gf) are relative to N2 samples.

| Molecular Formula | RT (min) | Fold Increase | Identity | Up in *him-5* | Up in *fem-3* (gf) | Up in 50:50 |
|---|---|---|---|---|---|---|
| $C_8H_{13}NO_4$ | 5.77 | 2.04 | | ND | No | ND |
| $C_7H_{13}NO$ | 6.5 | 48.8 | | No | No | ND |
| $C_{14}H_{18}NPO_8$ | 6.73 | 5.59 | iglu#2? | ND | ND | ND |
| $C_{14}H_{17}NO_5$ | 7.29 | 3.84 | iglu#1 | No | No | No |
| $C_8H_{14}N_5$ | 7.62 | 9.89 | | No | No | No |
| $C_{25}H_{42}NPO_7$ | 13.63 | 2.35 | LPE 20:5? | No | No | No |
| $C_{15}H_{29}NO_2$ | 14.04 | 5.29 | C14 lipid-ethanolamine? | No | No | 1.95 |
| $C_{23}H_{44}NPO_7$ | 14.54 | 2.27 | LPE 18:2? | No | No | No |
| 399.0445 [+] | 14.63 | 3.8 | | ND | No | ND |
| $C_{23}H_{46}NPO_7$ | 16.87 | 3.57 | LPE 18:1? | No | ND | ND |
| $C_{18}H_{38}N_2O_2$ | 17.67 | 28.68 | | No | No | No |
| 784.5828 [+] | 25.97 | 6.71 | | ND | No | No |

**NMR Spectra**

**(2*O*)- and (3*O*)-acetyl-*S*-methylthioadenosine (acemta#1 4 and acemta#2 5), $^{1}$H NMR spectrum (500 MHz, $CDCl_3$)**

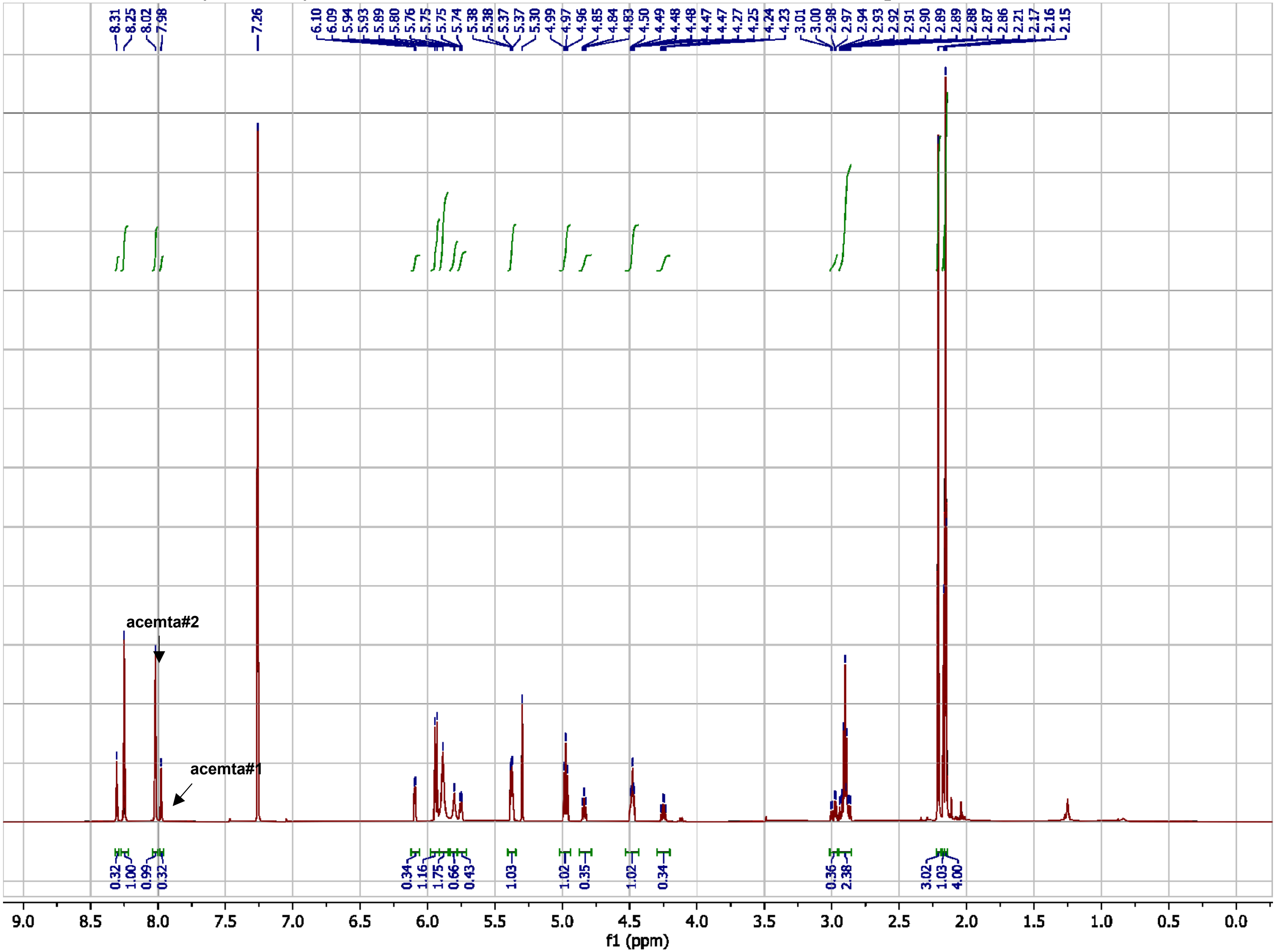

**acemta#1 and acemta#2 (4 and 5), $^{13}$C NMR spectrum (125 MHz, $CDCl_3$)**

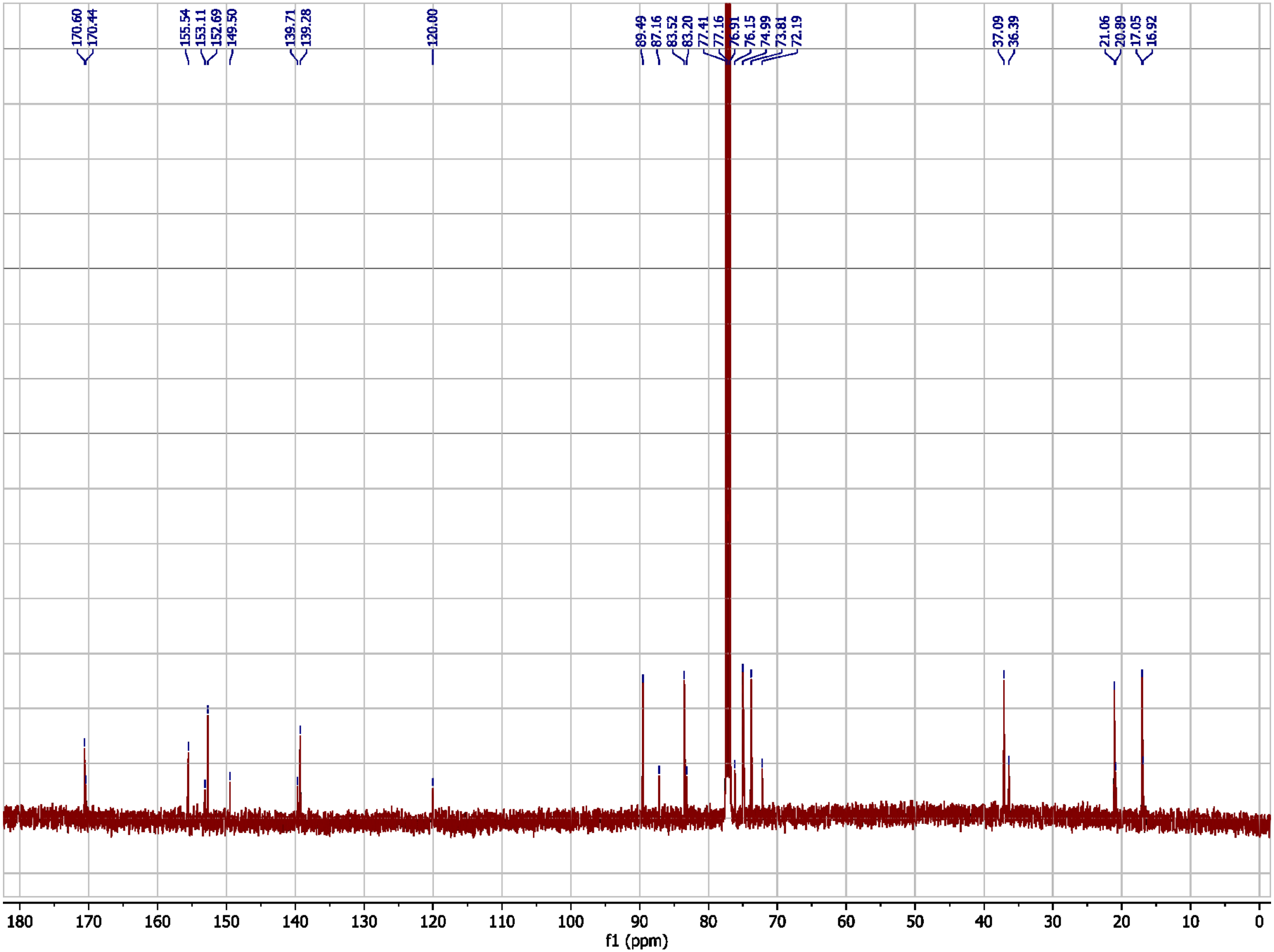

***N*,*N*-Dimethyltryptophan (24), $^{1}$H NMR spectrum (500 MHz, $D_6$-DMSO)**

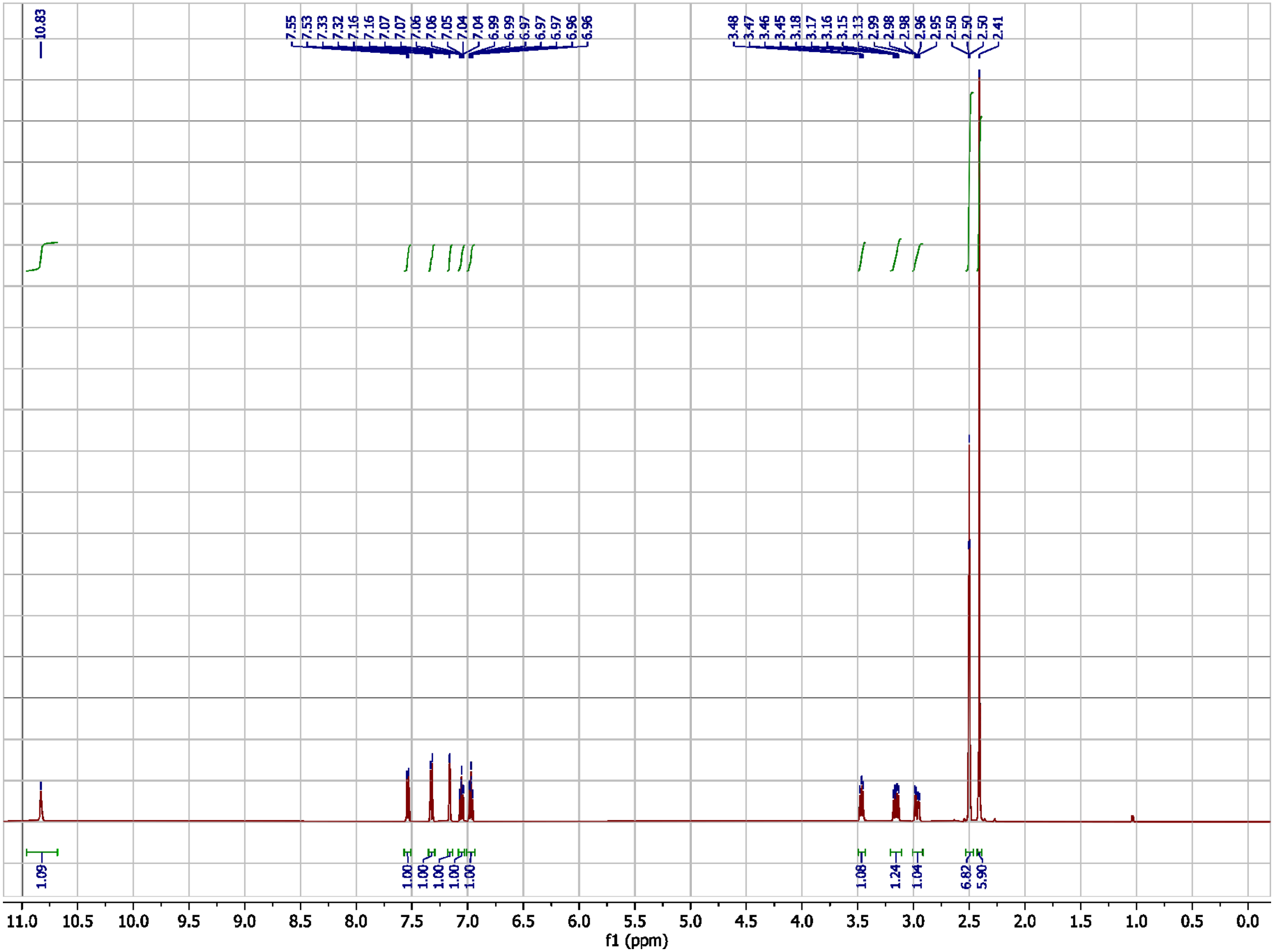

*N,N*-Dimethyltryptophan (24), $^{13}$C NMR spectrum (125 MHz, $D_6$-DMSO)

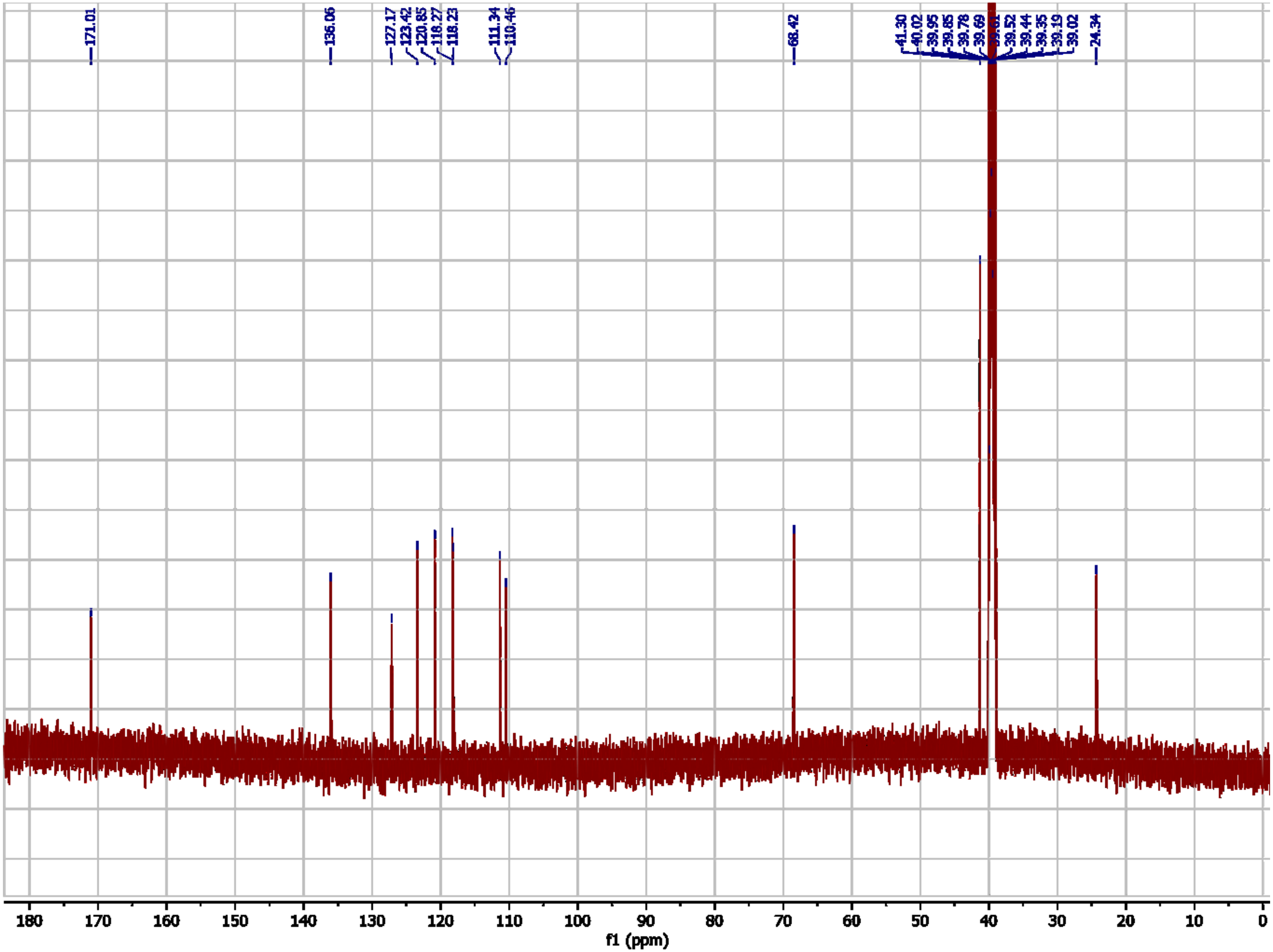

**medip#1 (26), [1]H NMR spectrum (500 MHz, $CD_3OD$)**

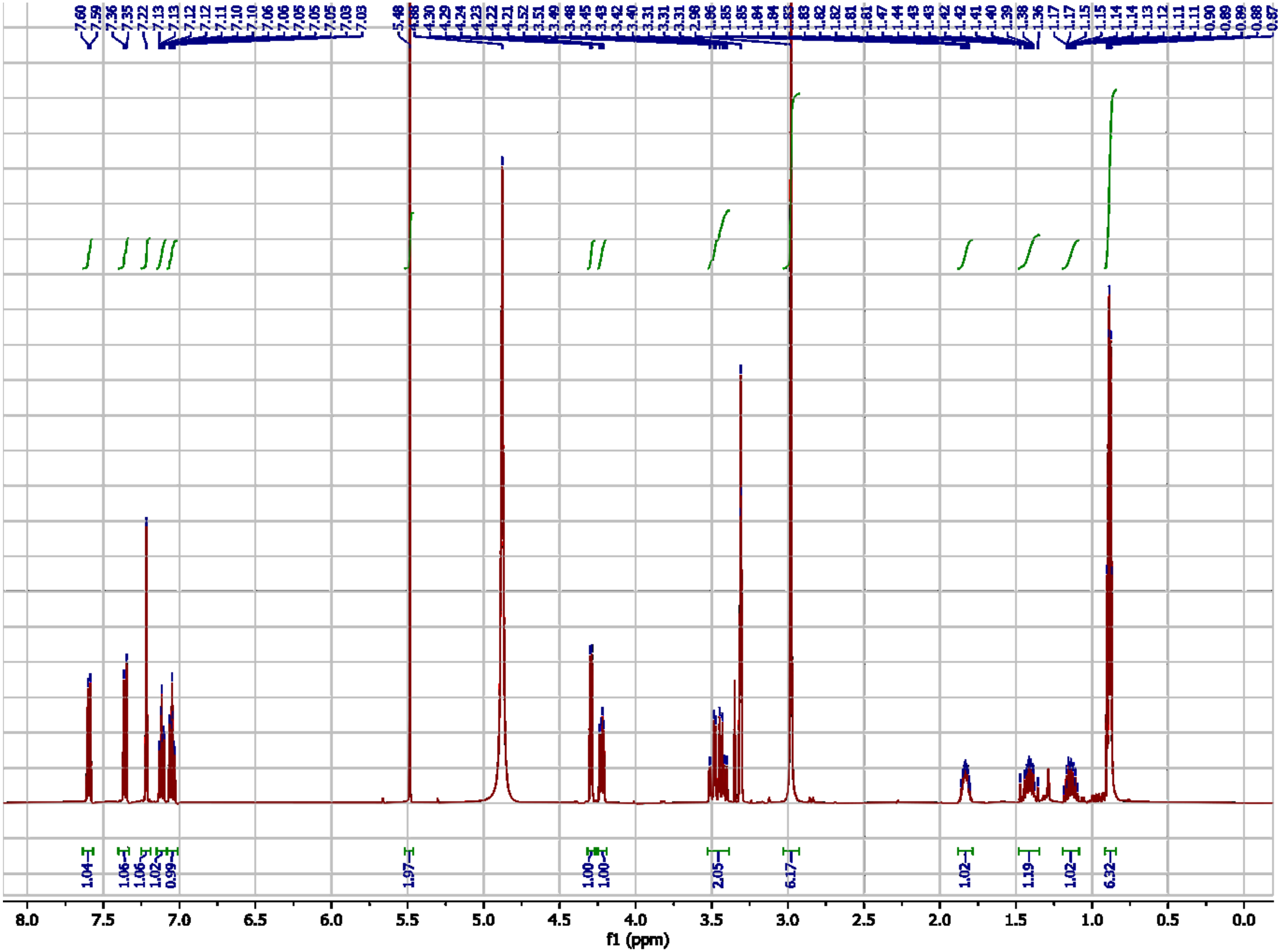

**medip#1 (26), $^{13}$C NMR spectrum (104  MHz, $CD_3OD$)**

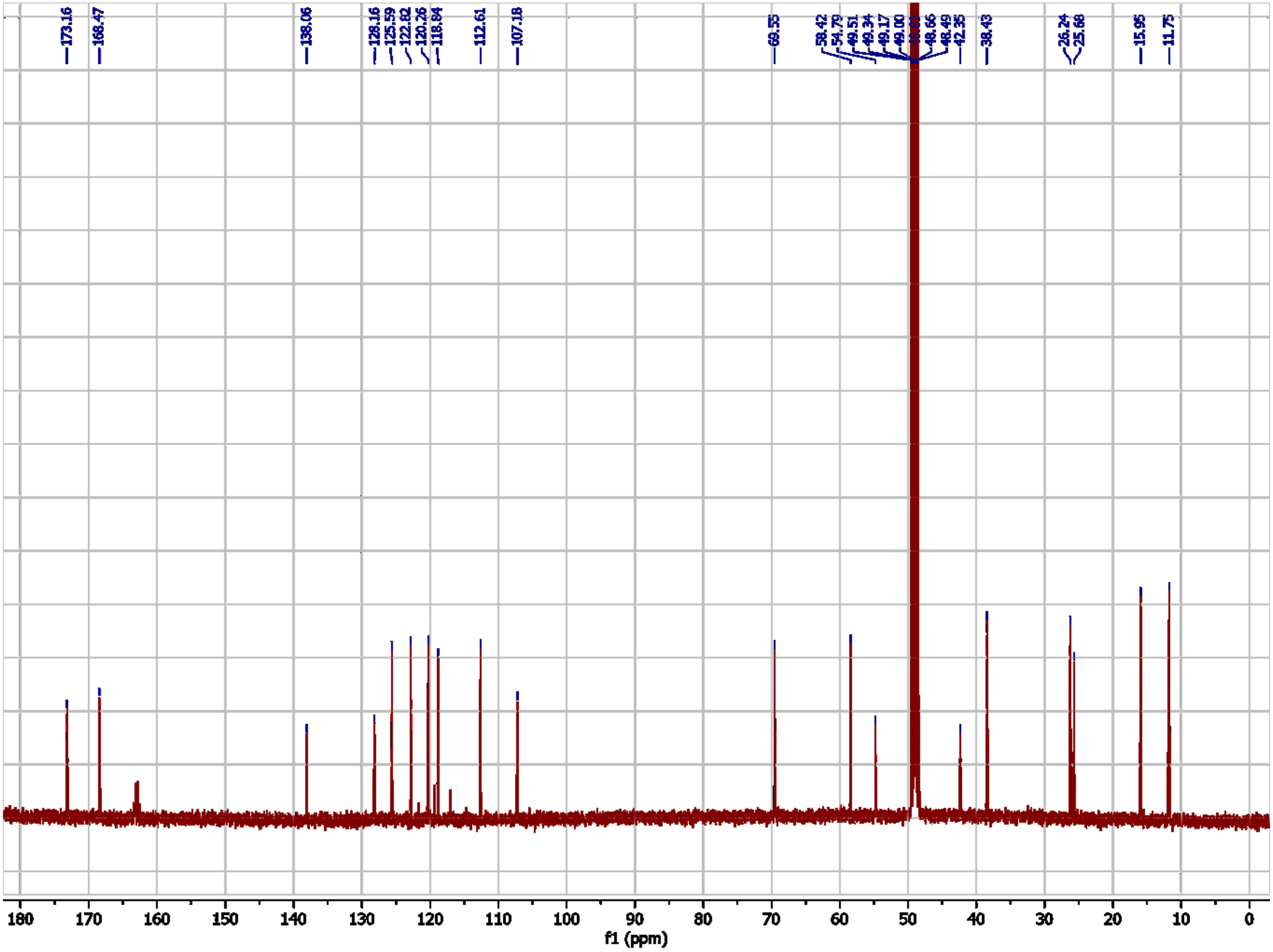
